# LRH-1/NR5A2 Activation Rewires Immunometabolism Blunting Inflammatory Immune Cell Progression in Individuals with Type 1 Diabetes and Enhances Human Islet Function in Mice

**DOI:** 10.1101/2023.09.18.558230

**Authors:** N Cobo-Vuilleumier, S Rodríguez-Fernandez, L López-Noriega, PI Lorenzo, JM Franco, CC Lachaud, E Martin Vazquez, R Araujo Legido, A Dorronsoro, R López-Férnandez-Sobrino, B Fernádez-Santos, D Salas-Lloret, N van Overbeek, M Ramos-Rodriguez, C Mateo-Rodríguez, L. Hidalgo, R Nano, AI Arroba, A Campos Caro, ACO Vertegaal, A Martin Montalvo, F Martín, M Aguilar-Diosdado, L Piemonti, L Pasquali, R González Prieto, MI García Sánchez, MA Martínez-Brocca, M Vives-Pi, BR Gauthier

## Abstract

The intricate etiology of type 1 diabetes mellitus (T1D), marked by a detrimental cross-talk between the immune system and insulin-producing β-cells, has impeded effective disease-modifying therapies. The discovery that pharmacological activation of the nuclear receptor LRH-1/NR5A2 can reverse hyperglycemia in mouse models of T1D by attenuating the autoimmune attack coupled to β-cell survival/regeneration, prompted us to investigate whether immune tolerization could be achieved in individuals with T1D by LRH-1/NR5A2 activation as well as improving islet function/survival after xenotransplantation in mice. Pharmacological activation of LRH-1/NR5A2 induced a coordinated genetic and metabolic reprogramming of T1D macrophages and dendritic cells, shifting them from a pro-to an anti-inflammatory/tolerogenic phenotype. Regulatory T-cells were also expanded resulting in the impediment of cytotoxic T-cell proliferation. LRH-1/NR5A2 activation enhanced human islet engraftment and function in hyperglycemic immunocompetent mice. In summary our findings demonstrate the feasibility of re-establishing immune tolerance within a pro-inflammatory environment, opening a new therapeutic venue for T1D.

## INTRODUCTION

Contrary to most autoimmune diseases, Type 1 Diabetes Mellitus (T1D) is gender impartial and stands out as one of the most prevalent chronic pediatric illnesses affecting 1.75 million individuals under the age of 20 years (https://diabetesatlas.org/atlas/t1d-index-2022/). A recent study estimated that in 2021 there were 355 900 new cases of T1D in children and adolescents worldwide of which only 56% were diagnosed. This number is projected to increase to 476 700 by 2050 (*1*). Classically, T1D has been considered a T-cell-mediated autoimmune disease caused by a disruption in the balance between T-regulatory cells (Tregs) and T-effector cells (Teffs; CD4^+^ and CD8^+^ cytotoxic T-cells) that respond to islet-associated self-antigens (*2, 3*). This breakdown in immune homeostasis or ‘tolerance’ leads to β-cell destruction resulting in insulin deficiency, hyperglycemia, and the lifelong necessity for insulin supplementation in afflicted individuals (*4*). Based on this immune origin, several immunosuppressive therapies have been developed and their safety demonstrated in clinical trials. One of these, teplizumab (anti-CD3 derivative) was recently approved by the Food and Drug Administration (FDA) and delays the development of T1D in individuals ‘at-risk’ by 2 years. While representing a significant breakthrough, the limited effectiveness timeframe underscores the disease’s more complex root cause (*5, 6*). Considering the latter, the contribution of β-cells in the pathogenesis of T1D has gained momentum, as evidenced by the expression of several T1D susceptible gene variants in the β-cells. These gene variants modulate pro-inflammatory signals and cause vulnerability to endoplasmic reticulum (ER) and oxidative stress, triggering cell dysfunction and apoptosis. Due to the highly vascularized islet microenvironment, β-cell stress signals reach circulating immune cells, initiating a cross-talk. This cross-talk may ultimately result in the destruction of β-cells and is further influenced by other genetic and environmental factors (*7–9*). Therefore new effective disease-modifying therapies (DMT) for T1D, should aim to resolve this detrimental dialogue by simultaneously targeting both immune and islet cells (*10*).

Nuclear receptors (NRs) play pivotal roles in a wide range of physiological and pathological processes (*11*). They regulate metabolic pathways that control cellular energy balance, survival, and adaptability to the environment. NRs are also crucial for whole-organism functions such as development, metabolism, reproduction, immune response, and tissue regeneration. The fact that NRs activities can be controlled by ligands has made them attractive targets for drug development, with potential therapeutic applications (*12, 13*). One such NR is the liver receptor homolog 1 (LRH-1 *a.k.a.* NR5A2), which has emerged as a promising drug target for diseases like diabetes, pancreatic cancer, non-alcoholic fatty liver disease, and metabolic syndrome. (*14–20*). Our previous work provided an early proof-of-concept that the specific pharmacological activation of LRH-1/NR5A2 using a small chemical agonist (BL001) could therapeutically impede the progression of hyperglycemia in 2 mouse models of T1D (NOD and RIP-B7.1) without long-term adverse effects, validating the benefits of targeting this NR (*21*). BL001 coordinated *in vivo* the resolution, rather than the suppression, of the autoimmune attack, by increasing the number of anti-inflammatory M2 while decreasing the number of pro-inflammatory M1 macrophages, and concomitantly increasing the number of tolerogenic dendritic cells and Tregs. In parallel, BL001 stimulated β-cell regeneration via trans-differentiation and improved cell survival, the latter involving the PTGS2/PGE_2_/PTGER1 signalling cascade (*21–23*).

Given the strong disease-modifying properties of LRH-1/NR5A2 activation in mouse models of T1D, and aiming for the clinical applicability of this strategy, herein we expanded our studies to primary immune cells obtained from individuals with T1D. Importantly, LRH-1/NR5A2 expression has been described in macrophages, dendritic and T-cells arguing for a direct and specific impact of BL001 on these immune cells (*24–26*). Our endpoint was to define the molecular mode of action of LRH-1/NR5A2 agonistic activation in T1D cells, which is especially relevant considering that the diabetic milieu impedes the anti-inflammatory characteristics of both human macrophages and dendritic cells (*27–29*). Furthermore, we sought to determine whether LRH-1/NR5A2 activation could facilitate long-term human islet engraftment and function in immunocompetent hyperglycemic mice.

## RESULTS

### BL001 treatment reduces inflammation in monocyte-derived macrophages and dendritic cells from T1D individuals

Peripheral blood mononuclear cells (PBMCs) were isolated from blood samples procured from individuals with T1D, with an average age 37 years, a body mass index (BMI) of 24, and of equal sex representation (Table 1). We focus on adults with long-established T1D as this population has been the most challenging to refrain from disease progression with novel immunomodulatory drug therapies. CD14^+^ monocytes were purified from PBMCs and derived either into naïve/primed (MDM1_0_) or pro-inflammatory macrophages (MDM1) and immature (iDC) or mature dendritic cells (mDC) (*30, 31*). Cell viability was greater than 90% for all groups. We initially profiled cell surface markers associated with either a pro-or anti-inflammatory/tolerogenic phenotype in the presence or absence of the pharmacological activation of LRH-1/NR5A2 using the small chemical agonist BL001. LRH-1/NR5A2 activation significantly reduced levels of the prototypical pro-inflammatory macrophage surface markers CD14 in both MDM1_0_ and MDM1 and CD80 in MDM1 (Fig. 1A and B). In parallel, BL001 increased the expression of the anti-inflammatory cell surface marker CD206 in MDM1_0_ (Fig. 1C). In contrast, expression levels of CD86, CD163, CD200R and CD209 remained relatively constant (Fig.S1A-D). Next, we measured the expression of several cell surface markers involved in antigen presentation and T-cell activation which are associated with a proinflammatory mDC phenotype. Agonistic activation of LRH-1/NR5A2 did not alter the expression of proinflammatory cell surface markers on iDCs but increased the expression of CD36 which is implicated in the clearance of apoptotic cells, inhibiting antigen presentation and DC maturation (Fig. 1D and Fig. S1E-M). In contrast, CD54, CXCR4, and CD25 expression levels were blunted in BL001-treated T1D mDCs (Fig. 1E-G). These findings collectively suggest that the pharmacological activation of LRH-1/NR5A2 not only reduces the expression of key pro-inflammatory cell surface markers in both T1D MDM1 (CD80 and CD14) and mDCs (CXCR4, CD54 and CD25) but also imposes an anti-inflammatory phenotype on naïve MDM1_0_ and iDCs, further limiting the recruitment/activation of additional MDM1 and mDCs. Of note, alterations in the expression of MDM and DC surface markers by BL001 were sex independent.

**Fig. 1.**
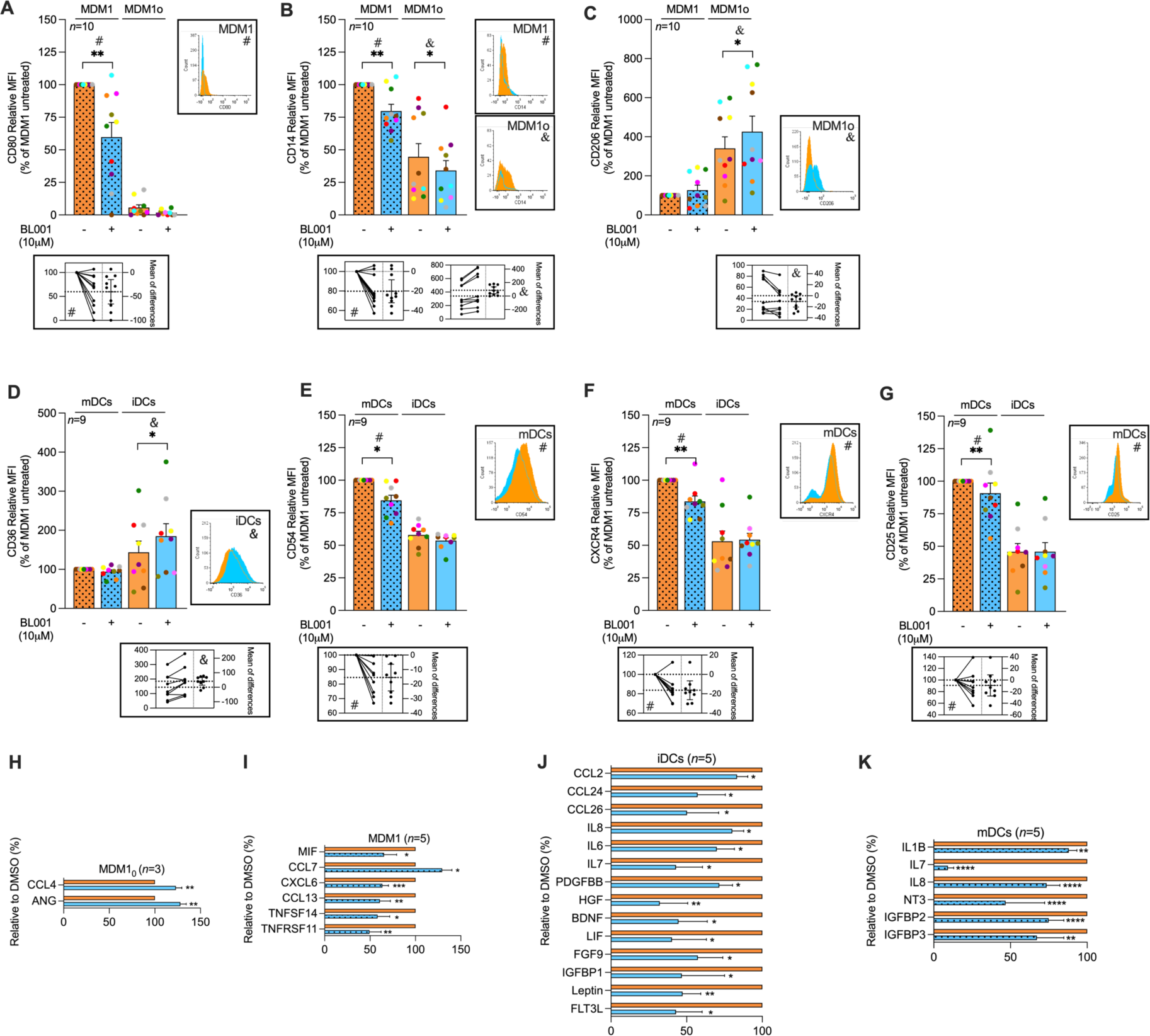
LRH-1/NR5A2 activation impedes the pro-inflammatory immune cell phenotype and cytokine secretion in T1D. Monocytes were purified from individuals with T1D, and differentiated into either resting or pro-inflammatory macrophage (MDM10 or MDM1) and immature or mature dendritic cells (iDCs or mDCs). LRH1/NR5A1 activation was achieved by administering 10 µM BL001 every 24 hours for a total duration of 48 hours with a final dose given 30 minutes before further analysis. MDM cell surface markers (**A**) CD80, (**B**) CD14 (**C**) CD206 and DC cell surface markers (**D**) CD36, (**E**) CD54, (**F**) CXCR4, (**G**) CD25 were then assessed by flow cytometry. Measurements were normalized to the mean fluorescence intensity (MFI) of untreated MDM1 or mDC for comparison. A total of *n=10* independent individuals with T1D were analyzed for MDM markers, while *n=9* independent individuals with T1D were evaluated for mDCs markers. Each donor is colour-coded. Data are presented as means ± SEM and compared to either untreated MDM1 or mDC. Paired Student t-test * p<0.05, and ** p<0.01 as compared to untreated cells (MDM10, MDM1, iDC and mDC). Flow cytometry histograms (untreated: orange, and BL001 treated: blue) as well as estimation plots (linked to plots by either & or #) are shown only for the markers with statistically significant differences. The cytokine secretion profile was assessed for (**H**) MDM10, (**I**) MDM1, (**J**) iDCs, (**K**) mDCs. A total of *n=3* independent donors were analyzed for MDM10, while *n=5* independent individuals were evaluated for MDM1, iDCs, mDCs. Percent changes in cytokine secretion are presented relative to DMSO-treated counterparts for each cytokine. Only significantly altered cytokines are shown. Data are presented as percent changes compared to DMSO for each cytokine. Unpaired Student t-test * p<0.05, ** p<0.01, *** p<0.001, **** p<0.001 compared to DMSO

**Table 1.**
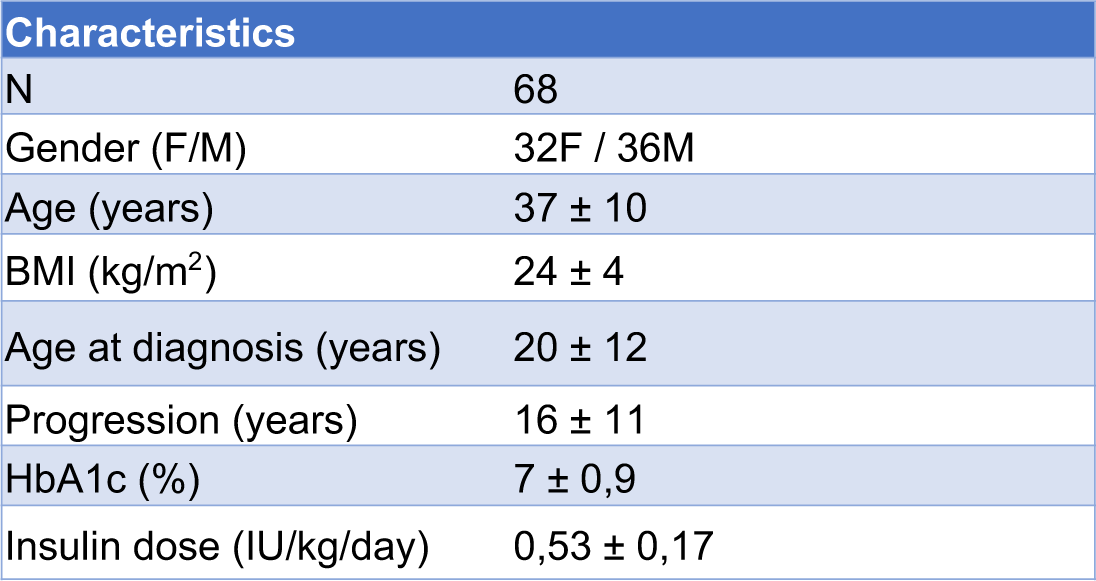
Demographic and clinical characteristics of donors with T1D.

The reduction of pro-inflammatory surface markers in both MDMs and DCs prompted us to assess whether the pharmacological activation of LRH-1/NR5A2 would also modulate their cytokine secretion profile, further favouring an anti-inflammatory environment. We found that BL001 treatment of T1D MDM1_0_ significantly increased the secretion of angiogenin (ANG) and CCL4 (Fig. 1H). Angiogenin is associated with an anti-inflammatory and wound-healing response in macrophage (*32*). CCL4 is recognized as a chemokine-attracting T-cells, including Tregs (*33*), but has recently also been shown to promote angiogenesis via angiopoietin-2 in oral squamous cell carcinoma (*34*). BL001 treatment significantly altered the secretion of 6 cytokines in T1D MDM1 (Fig. 1I). Noteworthy is the decreased secretion of CXCL6 and CCL13, both of which stimulate the chemotaxis of M1 macrophages (*35*). Additionally, BL001 reduced the secretion from of TNFSF14, a member of the TNF family that triggers the NF-κB signalling pathway leading to the induction of various chemokines including CXCL6 (*36*). The pharmacological activation of LRH1/NR5A2 also diminished the secretion of macrophage migration inhibitor factor (MIF) and TNFRSF11 (*a.k.a.* osteoprotegerin, OPG), two cytokines known for their pro-inflammatory activity.

Activation of LRH-1/NR5A2 in T1D iDCs significantly decreased the secretion of several members of the CC-chemokine subfamily (CCL2, CCL24, and CCL26), all of which play roles in various inflammatory diseases by recruiting leukocytes to sites of inflammation (Fig. 1J) (*37*). Correlating with this decline in chemokine secretion, FGF9, which is known to increase expression of pro-inflammatory chemokines like CCL2 in the central nervous system (*38*) exhibited lower levels in T1D iDCs treated with BL001. In parallel, the release of the pro-inflammatory cytokines IL-6, LIF (member of the IL-6 family), IL-7, and IL-8 as well as leptin, HGF, IGFBP1, FLT3 ligand, BDNF, and PDGFBB was reduced in BL001-treated T1D iDC (Fig. 1J). Of interest, Leptin is known for its role in driving DCs maturation, leading to Th1 priming, while the blockade of HGF is associated with the resolution of the pro-inflammatory phase, considered a critical step in restoring tissue homeostasis (*39, 40*). BL001-treated T1D mDCs also exhibited a reduction in the secretion of IL-7 and IL-8 (Fig. 1K). Two independent studies have demonstrated that blocking the IL-7 receptor could reverse autoimmune diabetes in NOD mice by repressing effector T-cells (Teffs) (*41, 42*) while high circulating levels of IL-8 have been associated with poor metabolic control in adolescents with T1D (*43*). More importantly, BL001 treatment decreased the secretion of the key cytokine IL-1B, which determines the inflammatory milieu during T1D (Fig. 1K) (*44*).

Taken together, our results indicate that LRH-1/NR5A2 activation reduces the secretion of several pro-inflammatory cytokines/chemokines in MDM1, mDCs, and iDCs, thereby attenuating the propagation of pro-inflammatory cells. Moreover, LRH-1/NR5A2 activation increases the secretion of pro-angiogenic cytokines ANG and CCL4 in MDM1_0_ while simultaneously decreasing the secretion of the anti-angiogenic factor HGF in iDCs. This intricate response underscores the multifaceted role of LRH1/NR5A2 activation in shaping the immune microenvironment towards an anti-inflammatory and regenerative state.

### LRH-1/NR5A2 agonistic activation attenuates the pro-inflammatory genetic signature of T1D MDMs

To investigate the molecular consequences resulting from LRH-1/NR5A2 pharmacological activation and understand its immunomodulatory effects at the cellular level, we conducted RNAseq analysis on T1D-isolated MDMs. Remarkably, MDM1_0_ displayed a greater number of differentially expressed genes (DEGs) than MDM1 upon BL001 treatment (1376 versus 671) (Fig. S2 and Fig. 2A and D). Gene set enrichment analysis (GSEA) revealed that common DEGs shared by MDM1_0_ and MDM1 subpopulations clustered into KEGG pathways predominantly related to hematopoietic cell lineage commitment, cholesterol, and fatty acid metabolism. In contrast, DEGs specific to MDM1_0_ clustered into cell cycle, genome dynamic, and RNA metabolism, whereas genes altered only in MDM1 clustered into inflammatory pathways and apoptosis (Fig. S2). Accordingly, functional enrichment analysis of DEGs in BL001-treated MDM1_0_ compared to untreated cells revealed the activation of enriched pathways for ribosomes, sphingolipid metabolism as well as autophagy. In contrast, cancer-related pathways (gastric cancer, p53, and transcription. misregulation in cancer) and the cell cycle pathway were suppressed as evidenced by decreased expression of numerous histone-encoding genes (Fig. 2B and C). Of particular interest was the up-regulation of DDIT4 and GRB10, both involved in the inhibition of mTOR (Fig. 2C, brown arrows) (*45, 46*). DDIT4 conveys the anti-inflammatory effects of IL-10 on pro-inflammatory M1 macrophages through the inhibition of mTOR and the activation of autophagy/mitophagy resulting in the clearance of damaged mitochondria, lower levels of reactive oxygen species and blunted NLRP3 inflammasome activation (*46*). In line with these effects, transcript levels of NLRP3 and its downstream target IL1B were reduced in BL001-treated MDM1_0_ (Fig. 2C, black arrows). Collectively, these results suggest that BL001 coerces MDM1_0_ into an anti-inflammatory and non-proliferative phenotype that secretes higher levels of regenerative factors (Fig. 1H). Notably, in BL001 treated T1D MDM1, lysosome was the only positively enriched pathway, while a myriad of pro-inflammatory-associated pathways such as the chemokine signalling pathway, cytokine-cytokine receptor interaction, and Toll-like receptor signalling, were suppressed (Fig. 2E). It is worth noting that lysosomal dysfunction has been linked to impair autophagy flux contributing to M1 macrophage polarization under diabetic conditions (*47*). This suggests that the activation of the lysosomal pathway, which includes the ACP5 gene that was significantly up-regulated in BL001-treated T1D MDM1 (Fig. 2D and F, red arrows), may be involved in suppressing the pro-inflammatory phenotype. Primary murine macrophages lacking ACP5 display a pro-inflammatory phenotype with increased secretion of IL-1B and IL12 (*48*). These cytokines, along with the inflammasome sensor NLRP3, the TLR4, and NF-κΒ signalling pathways were all decreased in BL001-treated T1D MDM1 (Fig. 2E and F, purple arrows).

**Fig. 2.**
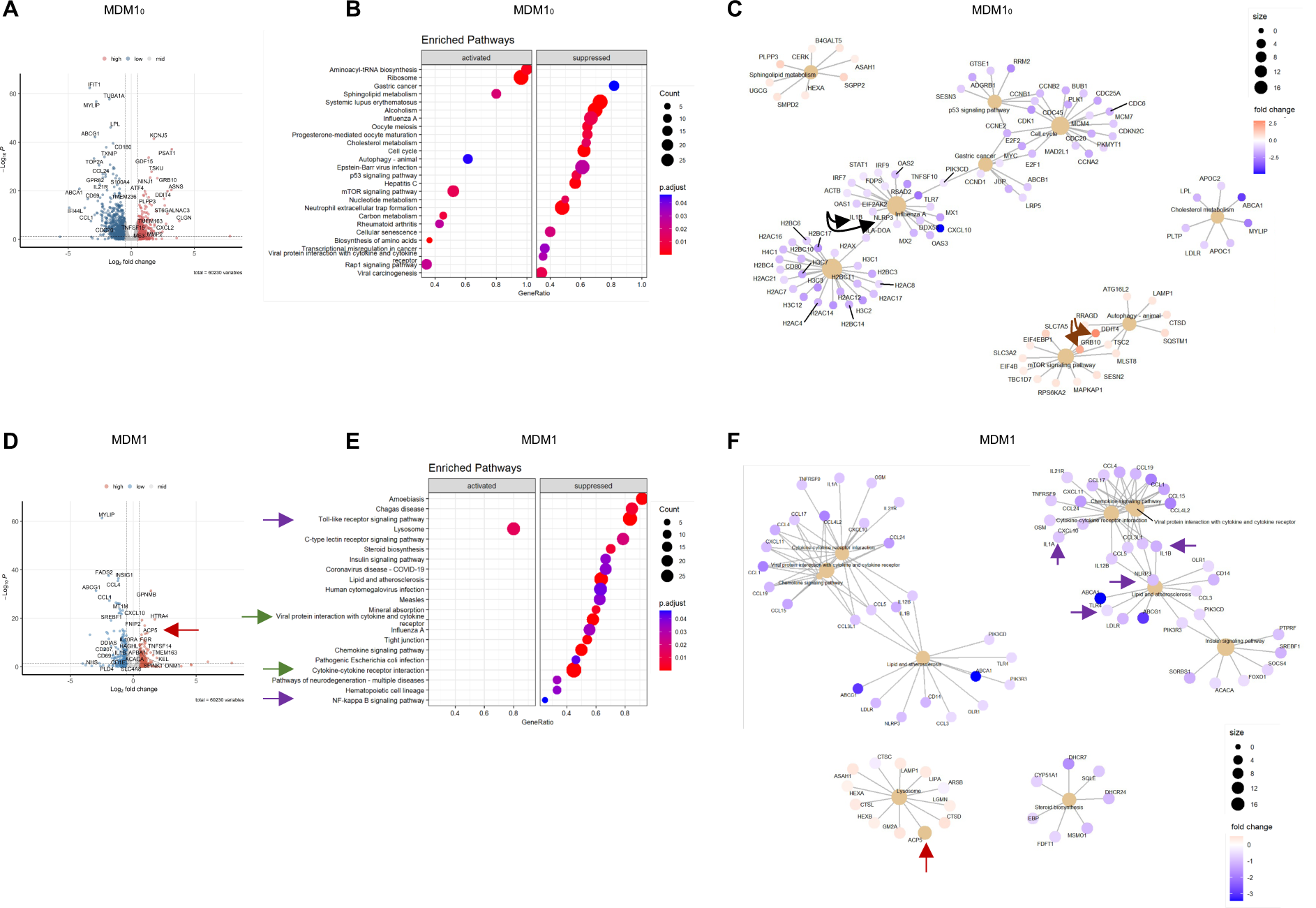
LRH-1/NR5A2 agonism mitigates the pro-inflammatory genetic program in T1D monocyte-derived macrophages (MDM). (**A**) Volcano plot of differentially expressed genes in BL001-treated vs untreated MDM10 (*n=6* independent donors). (**B**) Dot plot of KEGG pathways enriched in BL001-treated MDM10 and (**C**) Cnetplots of selected KEGG pathways. (**D**) Volcano plot of differentially expressed genes in BL001-treated vs untreated MDM1 (*n=6*, independent donors). (**E**) Dot plot of KEGG pathways enriched in BL001-treated MDM1 and (**F**) Cnetplots of selected KEGG pathways. Arrows highlight genes of interest which are described in the results.

### LRH-1/NR5A2 activation inhibits a subset of mitochondrial proteins in T1D MDM1_0_ and MDM1

To map global changes including post-transcriptional/translational alterations induced by LRH-1/NR5A2 activation, we determined the proteomic profile of T1D MDM1_0_ and MDM1 treated or not with BL001. This analysis revealed the presence of 188 differentially expressed proteins (DEPs) (87 up- and 91 down-downregulated proteins, *p<0.05*) out of a total of 1890 quantifiable proteins in MDM1_0_ (Fig. 3A, Table S1 and S2). Similarly, in T1D MDM1 we identified 287 DEPs (151 up- and 136 down-downregulated proteins, *p<0.05*) out of 1953 quantifiable proteins (Fig. 3B, Table S3 and S4). The relatively limited number of DEPs identified posed a challenge to the analysis of enriched pathways. Consequently, we investigated the protein-protein interaction (PPI) network among these DEPs. Consistent with the enrichment of the aminoacyl-tRNA biosynthesis and ribosome pathways observed in GSEA (Fig. 2B), several interacting proteins among the up-regulated DEPs in BL001-treated T1D MDM1_0_ were related to translation initiation and tRNA synthesis (Fig. 3C, grey shaded area). Remarkably, a PPI cluster of mitochondrial proteins was identified in the down-regulated DEPs of BL001-treated MDM1_0_ which was not identified by the GSEA (Fig. 3D, pink shaded area). Next, we performed an integrative analysis of the transcriptome and proteome to highlight common DEGs and DEPs. This analysis revealed 14 up- and 16-down regulated transcripts/proteins shared by both omics (Fig. 3E, green circles and F). Of particular interest, the transferrin receptor (TFRC), common to both omics, was the most up-regulated DEP in BL001-treated MDM1_0_ (Fig. 3A, C and F purple arrow). Deletion of this receptor in murine macrophages was shown to promote a M1-like polarization driven by IFNγ, suggesting that its increased expression may sway an anti-inflammatory phenotype (*49*). Similarly, BST2 was the most down-regulated DEP, consistent with a significant decrease in transcript levels in BL001-treated MDM1_0_ (Fig. 3A, D and F green arrow). BST2 is an anti-viral agent that induces pro-inflammatory gene expression via NF-κB (*50*), indicating that its suppression will likely impede MDM1_0_ activation towards an MDM1 phenotype.

**Fig. 3.**
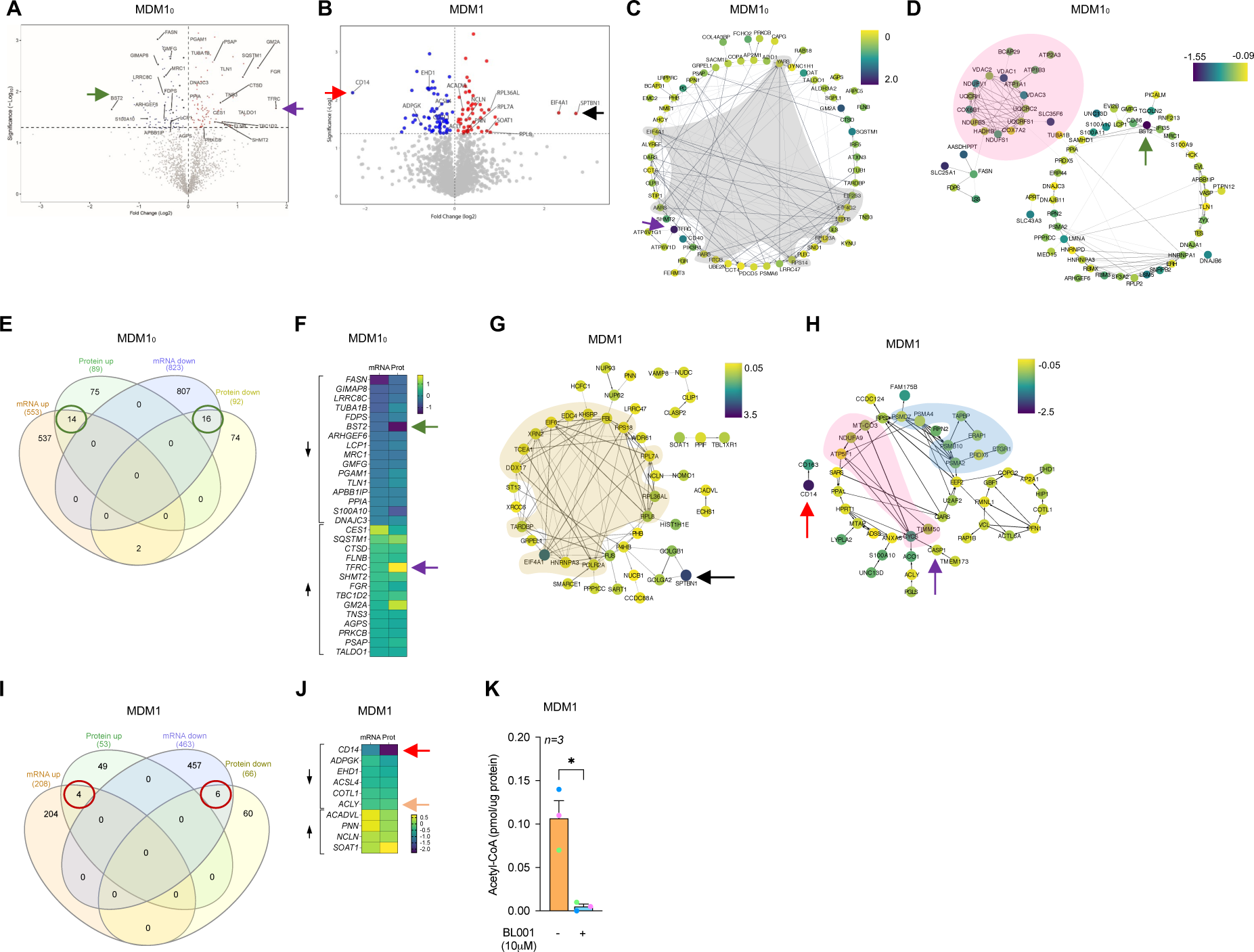
Proteomic alterations induced by BL001 in T1D monocyte-derived macrophages. Volcano plot displaying the most significantly differentially expressed proteins in (**A**) BL001-treated versus untreated MDM10, derived from *n=3* independent donors and (**B**) BL001-treated versus untreated MDM1, from *n=3* independent donors. Arrows point to genes of interest which are described in the results section. Cytoscape circular layout of significantly (**C**) up-regulated proteins and (**D**) down-regulated proteins in BL001-treated MDM10 compared to untreated controls. RNA-associated proteins are highlighted within the grey-shaded area while mitochondrial proteins are emphasized by the pink-shaded area. (**E**) InteractiVenn diagram of differentially expressed transcripts/proteins that are significantly altered in either the RNAseq or proteomic analysis of BL001-treated versus untreated MDM10. (**F**) Heatmap of differentially expressed transcripts/proteins common to both the RNAseq and proteomic analysis in BL001-treated MDM10 versus untreated MDM10 marked by green circles in (**E**). Cytoscape circular layout of significantly (**G**) up-regulated proteins and (**H**) down-regulated proteins in BL001-treated MDM1 as compared to untreated MDM1. Proteins involved in transcriptional/translational processes are within the French beige-shaded area while mitochondrial proteins are highlighted in the pink shaded area and proteasome-associated proteins are in the blue-shaded area. (**I**) InteractiVenn diagram of differentially expressed transcripts/proteins that are significantly altered in either the RNAseq or proteomic analysis of BL001-treated versus untreated MDM1. (**J**) Heatmap of differentially expressed transcripts/proteins common to both RNAseq and proteomic analysis of BL001-treated MDM1 versus untreated MDM10(red circles in I). Arrows point to genes of interest which are described in the results section. (**K**) Bar graph representing acetyl CoA levels in MDM1 treated or not with BL001 for *n=3* independent donors colour-coded and each performed in triplicate. Data are presented as means ± SEM. Student t-test * p<0.05 as compared to untreated.

In contrast to the GSEA in BL001-treated MDM1, which revealed lysosome as the only activated pathway (Fig. 2E), most of the up-regulated DEPs interacting with each other, clustered in global gene transcription/translation, including RNA polymerase (POLR2A), translation initiation factors (EIF4A1, the second most up-regulated DEP) and ribosomal subunits (RPL8, RPL7A, etc.) (Fig. 3G, French beige shaded area). The most up-regulated DEP, SPTBN1, was shown to inhibit inflammatory responses and hepatocarcinogenesis in hepatocellular carcinoma via down-regulation of the NF-κB signalling pathway, which was also suppressed in T1D MDM1 (*51*) (Fig. 3B and G, black arrow and Fig. 2E). Significantly, CD14 exhibited the strongest down-regulation among DEPs in BL001-treated MDM1, aligning with the flow cytometry results and providing robust evidence of BL001’s potent anti-inflammatory effect (Fig. 3B, H and J, red arrow and Fig. 1B) (*52, 53*). This conclusion is substantiated by the observed decrease in proteins associated with the proteasome, for which inhibition results in a conversion to an anti-inflammatory phenotype (Fig. 3H, blue shaded area) (*54*). Similar to MDM1_0_, several mitochondrial proteins were repressed in BL001-treated MDM1, evidencing that LRH-1/NR5A2 activation affects mitochondrial function (Fig. 3H, pink shaded area) (*20, 55*). We then compared and contrasted the transcriptome and proteome of BL001-treated MDM1 and found that 4 up- and 6 down-regulated genes were common to both omics approaches (Fig. 3I, red circles and J). CD14 was the most down-regulated gene/protein, emphasizing that BL001 blocks the initial proinflammatory signalling receptor to prevent further stimulation by LPS (*56*) (Fig. 3J, red arrow). Of particular interest, ATP-citrate lyase (ACLY) was also consistently decreased in both omics and resulted in lower levels of Acetyl CoA in BL001-treated MDM1s (Fig. 3J, orange arrow and K). LPS promotes histone acetylation via increased ACLY activity and higher levels of Acetyl CoA, which licences the transcription of pro-inflammatory genes such as IL1B (*57*). Our data suggest that lower levels of ACLY and Acetyl CoA could result in decreased histone acetylation, leading to the repression of the pro-inflammatory gene response.

### LRH-1/NR5A2 activation drives mitohormesis to enforce a pro-inflammatory resistant state in T1D MDM1

A hallmark of LPS-induced pro-inflammatory reprogramming of macrophages is a time-dependent shift in ATP production from oxidative phosphorylation (OXPHOS) to glycolysis (*58*). As a consequence of these metabolic changes and as a feedback mechanism to prevent cell impairment, macrophages trigger a stress response called mitohormesis (*59*). This response, which includes a cross-talk between the nuclei and the mitochondria, attempts to re-establish mitochondrial homeostasis and induce an LPS tolerance state, thereby avoiding an exacerbated and perpetual pro-inflammatory response (*60*). As LRH-1/NR5A2 pharmacological activation suppressed the expression of mitochondrial proteins in both MDM1_0_ and MDM1, we wondered whether BL001 could induce mitohormesis in macrophages. To validate this premise, we assessed the mitochondrial metabolic flux in T1D MDMs. The addition of LPS to MDM1_0_ to generate MDM1 significantly reduced the basal oxygen consumption rate (OCR) as compared to non-LPS-treated MDM1_0_, consistent with the switch from OXPHOS to glycolysis to support a pro-inflammatory phenotype (Fig. 4A and B). BL001 further decreased the basal OCR in MDM1 as well as in MDM1_0_ even to lower levels than those found before the LPS treatment (Fig. 4A and B). Accordingly, transcript and protein levels of ATF4, the key activator of mitohormesis, as well as transcript levels of its downstream target GDF15 were significantly increased (Fig. 4C-E) whereas the expression levels of IL1B and the inflammasome sensor NLRP3 were blunted in BL001-treated T1D MDM1 (Fig. 4F and G). Noteworthy, protein levels of caspase-1 (CASP1), a downstream target of the NLRP3-inflammasome responsible for IL-1B activation (*61*), were also reduced in BL001-treated MDM1 cells (Fig. 3H, purple arrow). Taken together, these results suggest that LRH-1/NR5A2 agonistic activation likely triggers mitohormesis to enforce a pro-inflammatory immune-paralyzed state with a concomitant inhibition of the inflammasome blunting further activation of cytokines. We then assessed whether this immune-paralyzed state could impact stimulation of naïve CD4^+^ and CD8^+^ T-cell proliferation, which is typically triggered by professional antigen-presenting cells such as macrophages and DCs (*62, 63*). CD4^+^, but not CD8^+^ T-cell proliferation was dampened by BL001-treated MDM1 derived from T1D individuals, indicating that BL001 reprogramming of MDM1 partly inhibits T-cell proliferation (Fig. 4H and I).

**Fig. 4.**
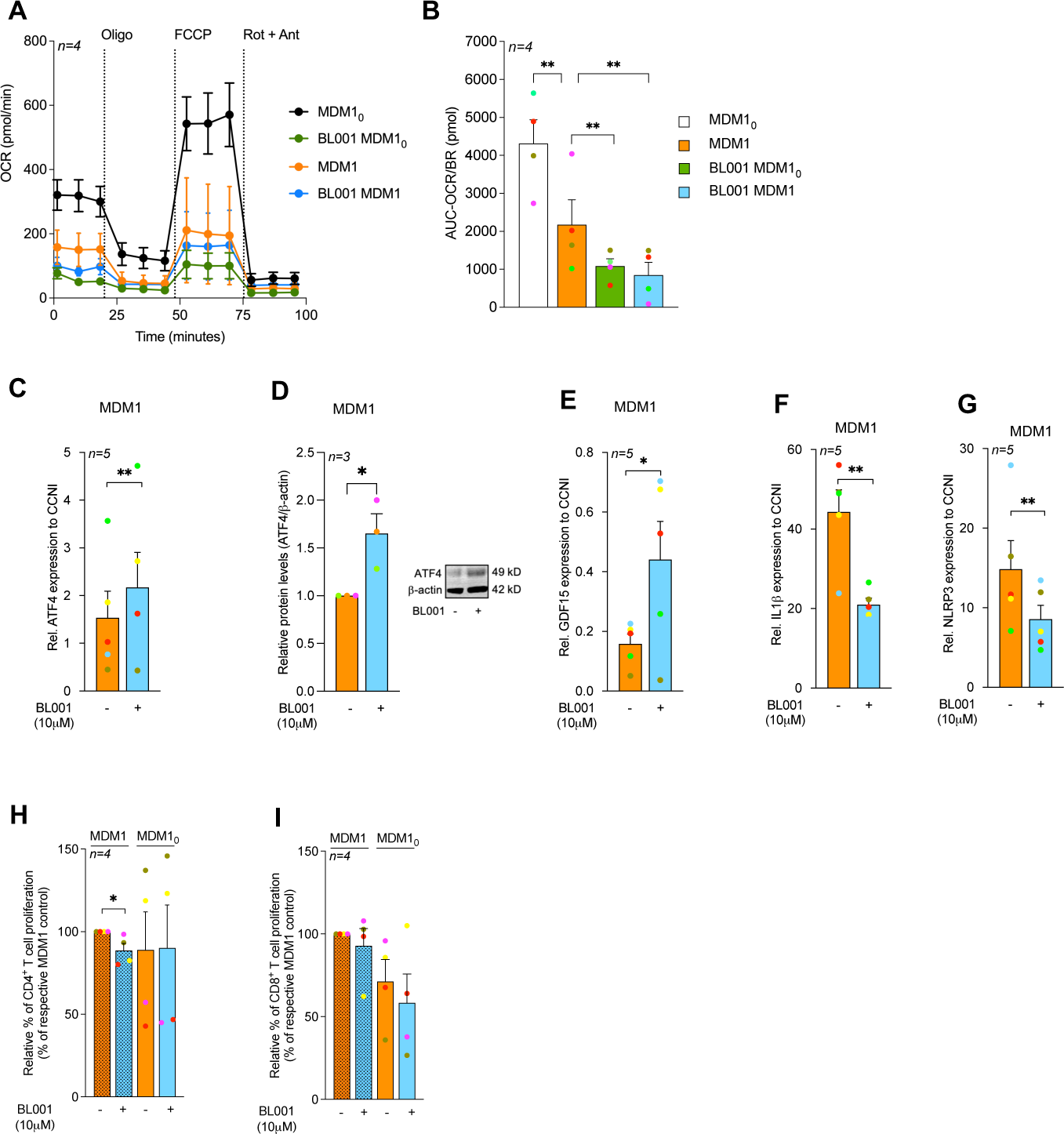
LRH-1/NR5A2 activation stimulates mitohormesis to enforce LPS-tolerance in T1D monocyte-derived macrophages. (**A**) Mitochondrial stress test performed on T1D MDM10 and MDM1 (namely LPS/IFNg-treated MDM1_0_) treated with or without BL001. *n=4* independent donors. (**B**) Calculated basal oxygen consumption rates (OCR-BR) Abbreviations: Oligo-Oligomycin, FCCP-Carbonyl cyanide-p-trifluoromethoxyphenylhydrazone, Rot-Rotenone, Ant-Antimycin A. Paired Student t-test **p<0.01 as compared to MDM10 or MDM1. *n=4* independent colour-coded donors. (**C**) Transcript levels of the mitohormesis-associated gene ATF4, in BL001 treated or not MDM1 cells from T1D donors, normalized to the housekeeping gene CCNI (https://housekeeping.unicamp.br/). *n=5* independent colour-coded donors. (**D**) Protein expression levels of ATF4 in BL001 treated or not MDM1 cells from T1D individuals, and normalized to the housekeeping protein b-actin. *n=3* independent colour-coded donors. The figure includes a representative western blot image. Transcript levels of (**E**) GDF15, (**F**) IL-1b, and (**G**) NLRP3 in T1D MDM1 treated with or without BL001. Transcript levels were normalized to the housekeeping gene CCNI (https://housekeeping.unicamp.br/). *n=5* independent colour-coded donors. Data are presented as means ± SEM. Paired Student t-test * p<0.05, **p<0.01 as compared to untreated cells. Relative proliferation of autologous (**H**) CD4^+^ and (**I**) CD8^+^ T cells in response to co-culture with healthy or T1D MDMs treated or not with 10 µM BL001. *n=4* independent colour-coded donors. Paired Student t-test *p<0.05 as compared to untreated cells.

### NR5A2 promotes mitochondrial turnover favouring tolerization of T1D mDCs

Having defined the molecular mode of action of LRH-1/NR5A2 in macrophages, we next focused on the genetic adaptations induced by BL001 in DCs. To this end, we performed an RNAseq analysis in T1DM iDCs and mDCs treated or not with BL001. Interestingly, BL001 altered the expression of only few genes in T1DM iDCs, with no significant changes detected in either KEGG pathways or GO enrichment terms (Fig. 5A). In contrast, BL001 treatment altered the expression of numerous genes in T1DM mDCs (Fig. 5B). Functional enrichment analysis of BL001-treated T1D mDCs revealed the activation of some pathways, including the lysosomal pathway, and suppression of others such as ‘viral protein interaction with cytokine and cytokine receptor’ and ‘cytokine and cytokine receptor interaction’, both of which were also decreased in T1D MDM1 treated with BL001 (Figure 5C, E and Fig. 2E, green arrows). The most activated pathway in BL001-treated T1D mDCs was OXPHOS (Fig. 5C red arrow and E). Similar to macrophages, a switch from OXPHOS towards glycolysis is the main trigger activating mDCs (*64*). Thus, stimulation of OXPHOS via increased fatty acid oxidation (FAO) driven by the PPAR signalling pathway (also among the top activated pathways-Fig. 5C blue arrow) may induce an anti-inflammatory and tolerizing phenotype in mDCs. Supporting this premise, several key tolerogenic genes such as VEGFA and B, TGFB1 and 2, PTGS2, and CD36 were all significantly up-regulated in BL001-treated mDCs (Fig. 5D). To ascertain this hypothesis, we assessed the mitochondrial metabolic flux of DCs isolated from T1D individuals treated with or without BL001 and palmitate. Similar to MDM, maturation/activation of iDCs resulted in decreased basal OCR (Fig. 5F and G). However, the addition of BL001 alone or together with palmitate did not increase/potentiate basal OCR as compared to untreated mDCs (Fig. 5F and G). A broaden analysis revealed upregulation of the lysosomal and phagosome pathways in BL001-treated mDCs, along with genes associated with increased acidification and hydrolase activity (Fig. 5C, pink arrows and E). These findings suggest a potential induction of mitophagy to eliminate dysfunctional mitochondria, coinciding with the generation of new functional mitochondria as indicated by the increased expression of mitochondrial-related genes (Fig. 5E). To corroborate this premise, mitochondrial mass and membrane potential were assessed by flow cytometry using the probes MitoTracker Green (binding covalently to mitochondrial proteins, thus providing an assessment of mass) and MitoTracker Red (taken up by polarized mitochondria, thus gauging function) (*65*). BL001-treated T1D mDCs displayed a reduced proportion of cells with non-functional mitochondria (MitoTracker Green^High^ and MitoTracker Red^low^) compared to untreated cells (Fig. 5H and I). A comparative proteomic profiling was next conducted on BL001-treated mDCs compared to vehicle-treated cells, aiming to evaluate if DEPs associated with mitochondrial biogenesis. Two hundred and forty five DEPs (201 up- and 41 down-downregulated proteins, *p<0.05*) were identified out of a total of 6181 quantifiable proteins in mDCs (Table S5 and S6). Functional classification through GO analysis demonstrated that a significant portion of up-regulated DEPs were enriched in biological processes related to mitochondrial homeostasis - turnover, biogenesis and function (Fig. 5J red biological processes and Table S7). In parallel, down-regulated DEPs segregated to several GO biological processes link to oxidative stress and mitochondria death (Fig. 5K, red biological processes). Taken together, these results support the premise that LRH-1/NR5A2 activation in mDCs facilitates mitochondrial turnover, which is associated with the emergence of a tolerogenic phenotype. In line with this, BL001 impeded mDCs-mediated stimulation of autologous CD4^+^ and CD8^+^ T-cell proliferation in individuals with T1D (Fig. 5L and M). Furthermore, BL001-treated T1D iDCs also displayed reduced capacity to stimulate CD8^+^ T-cell proliferation, which correlated with increased levels of CD36 in these cells (Fig. 5M and Fig. 1D).

**Fig. 5.**
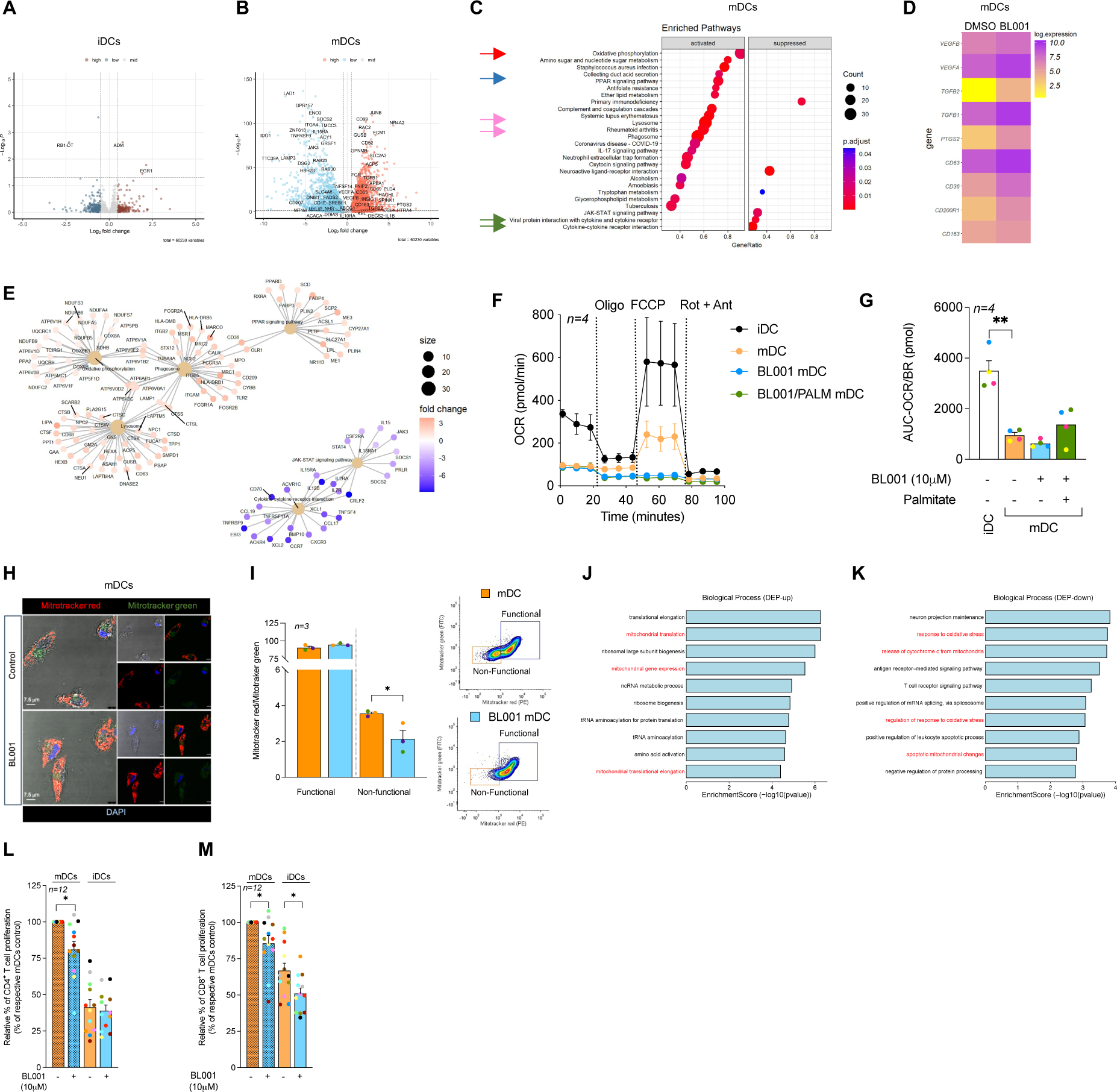
LRH-1/NR5A2 Promotes a Tolerogenic Phenotype to T1D mDC, Suppressing Autologous T-cell Proliferation. Volcano plot of differentially expressed genes in BL001-treated vs untreated (**A**) iDCS (*n=5* independent donors) and (**B**) mDCs (*n=4* independent donors). (**C**) Dot plot of KEGG pathways enriched in BL001-treated mDCs. Arrows point to pathways of interest which are described in the results section. (**D**) Heatmap of selected genes associated with a tolerogenic DC phenotype. Gene expression is presented as the average log(NC). (**E**) Cnetplots of selected KEGG pathways modulated by BL001 in mDCs. (**F**) Mitochondrial stress test and (**G**) calculated basal oxygen consumption rates (OCR) in T1D iDC and mDC treated with or without BL001 and with palmitate. Abbreviations: Oligo-Oligomycin, FCCP-Carbonyl cyanide-p-trifluoromethoxyphenylhydrazone, Rot-Rotenone, Ant-Antimycin A. *n=4* independent donors (colour matched). Paired Student t-test ** p<0.01 as compared to iDCs. Each donor is colour-coded. (**H**) Representative confocal immunofluorescence images of cells labeled with MitoTracker green and MitoTracker red for mitochondria, with nuclei counterstained using DAPI. (I) Bar chart quantification for mitochondrial functionality, as determined by flow cytometry based on MitoTracker green and MitoTracker red. Representative flow cytometry plots are provided for illustration. *n=3* independent colour-code donors. Paired Student t-test * p<0.05 as compared to untreated mDC. Bar plot ranking of the top ten GO biological process terms associated with (**J**) up-regulated and (**K)** down-regulated proteins in BL001-treated mDCs. Relative autologous proliferation of (**L**) CD4^+^ and (**M**) CD8^+^ T-cells in the presence of T1D DCs treated with or without 10 µM BL001. Data are presented as means ± SEM. Paired Student t-test * p<0.05 as compared to untreated cells.

### BL001 promotes the expansion of a CD4^+^/CD25^+^/FoxP3^+^ T-cells subpopulation in individuals with T1D which inhibits CD8^+^ T-cell proliferation

We next assessed whether BL001 could also impart an anti-inflammatory landscape to T-cell subpopulations within PBMCs, including the expansion of FoxP3^+^ T-cells, which enforce peripheral self-tolerance. PBMCs treated with BL001 had a higher number of CD4^+^CD25^+^ T-cells, alongside a modest, albeit non-significant, increased in a FoxP3^+^ cell subpopulation as compared to untreated PBMCs (Fig. 6A and B). This observation suggests a potential trend towards an enhanced regulatory Treg profile. Concurrently, a decrease in Th1 cell population and an increase in Th2 cells were noted, albeit not reaching statistical significant, indicating a shift in helper T-cell dynamics (Fig 6C and D). No changes were observed in the populations of Th17/Th22 or CD8^+^ T-cells, pointing to a selective impact of BL001 on specific CD4^+^ T-cell subsets (Fig 6E and F). Upon activation of CD4^+^ T-cells, a significant increase in the CD4^+^CD25^+^FoxP3^+^ T-cell subpopulation was further observed in the presence of BL001, suggesting that the pharmacological activation of LRH-1/NR5A2 may promote a regulatory phenotype under conditions of T-cell activation (Fig. 6G). Silencing of LRH1/NR5A2 led to a decrease in FoxP3 levels in CD4^+^ T-cells suggesting that the nuclear receptor regulates the expression of FoxP3 in these cells (Fig. 6H). Although CD4^+^ T-cells proliferation showed no significant changes, CD8^+^ T-cells proliferation was decreased by BL001 treatment of activated T-cells (Fig 6I and J). Taken together these results emphasize the pivotal role of LRH-1/NR5A2 in the regulation of Treg-associated markers and the maintenance of a regulatory T cell phenotype.

**FIG. 6.**
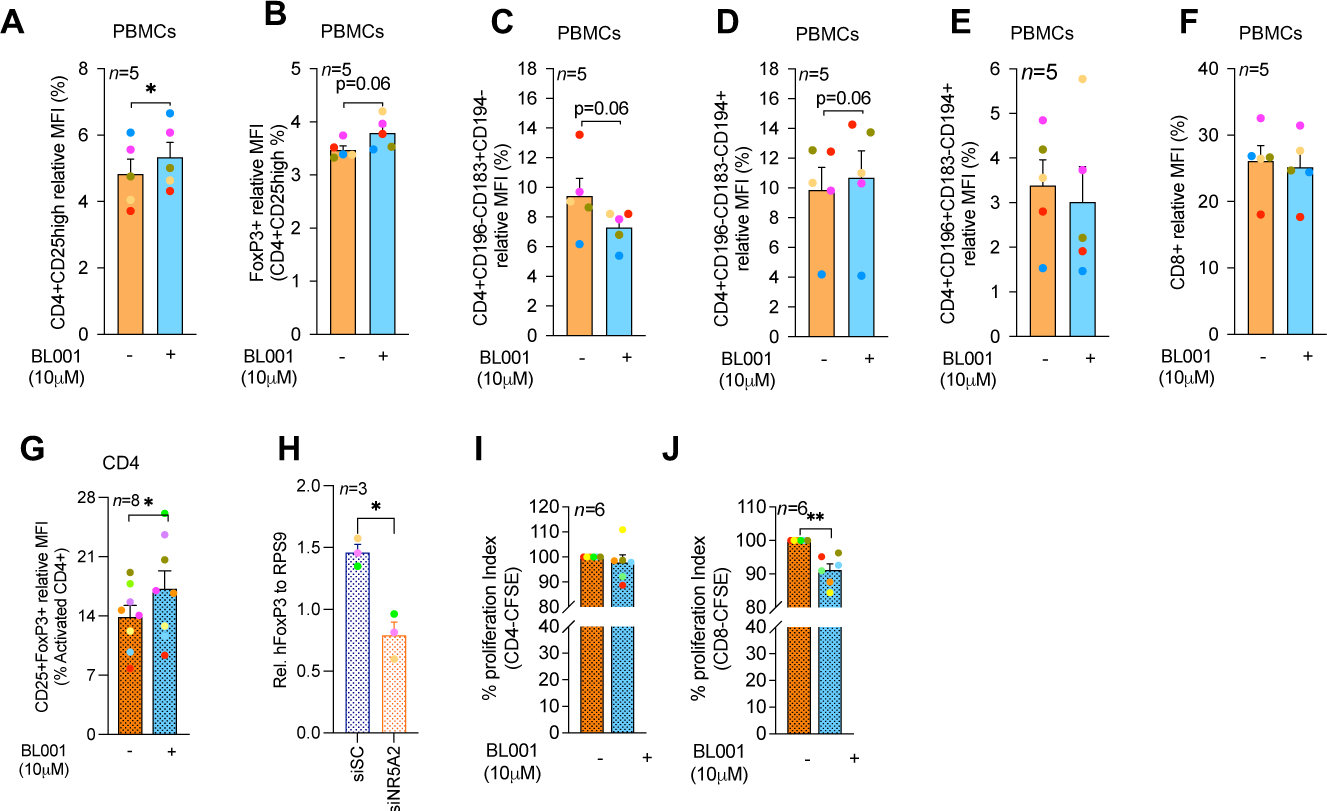
LRH-1/NR5A2 agonism promotes the expansion of a CD4^+^/CD25^+^/FoxP3^+^ cell subpopulation, leading to reduced CD8^+^ T-cell proliferation in T1D individuals. PBMCs were purified from individuals with T1D and exposed to 10 µM BL001 every 24 hours for a total duration of 48 hours with a final dose given 30 minutes before further analysis. Cells were then analysed by flow cytometry using a combination of cell surface markers and gated for subpopulations of (**A**) CD4^+^CD25^+^, (**B**) Tregs; CD4^+^CD25^+^FoxP3^+^ (**C**) Th1; CD4^+^CD196^-^CD183^+^CD194^-^, (**D**) Th2; CD4^+^CD196^-^CD183^-^CD194^+^, (**E**) Th17/22; CD4^+^CD196^+^CD183^-^CD194^+^ and (**E)** CD8^+^. *n=5* independent individuals with T1D. (**G**) CD4^+^ cells were isolated from PBMCs and treated with BL001 as described above. Cells were then analysed by flow cytometry for the cell surface markers CD25^+^FoxP3^+^ and results plotted as the percentage of CD25^+^FoxP3^+^ cells within the CD4^+^ subpopulation. *n=8* independent individuals with T1D. All cytometry measurements were normalized to the mean fluorescence intensity (MFI). (**H**) Relative FoxP3 transcript levels in either siScrambled (siSc) or siNR5A2-treated PBMCs. Data were normalized to the housekeeping gene RSP9. *n=3* T1D independent donors. CD14^-^ PBMCs were labelled with CFSE and stimulated/expanded using a polymeric nanomatrix structure composed of CD3 and CD28 for 24 hours before the addition of CD4^+^ T-cells (at a 1:2 ratio, respectively, from the same donor). Proliferation of (**I**) CD4^+^/CFSE^+^ and (**J**) CD8^+^/CFSE^+^ subpopulations was assessed by flow cytometry 4 days post co-culturing. *n=6* T1D independent donors. Each donor is colour-coded. Data are presented as means ± SEM. Paired Student t-test * p<0.05, and ** p<0.01 as compared to untreated cells.

### BL001 improves human islet graft survival and function in STZ-treated immune-competent mice

We have demonstrated that pharmacological activation of LRH-1/NR5A2 confers anti-inflammatory/tolerogenic properties to human macrophages, DCs, while promoting expansion of CD4^+^/CD25^+^/FoxP3^+^ T-cells. The current results extend our previous findings on BL001 to promote immune tolerance and enhance islet cell survival in the RIP-B7.1 and streptozotocin-induced T1D mouse models (*21*). Given these common benefits on both murine and human immune cells, we next proceeded to evaluate whether BL001 could attenuate graft rejection and improve the survival and function of human islets transplanted into hyperglycemic and immunocompetent C57BL/6J mice. To this end, we performed suboptimal human islet transplantations (750 human islet equivalents; IEQ) under the kidney capsule of mice rendered diabetic with a single high dose of streptozotocin (STZ). One week after transplantation, mice were treated or not with BL001. The rationale for using suboptimal amounts of IEQ was to examine whether any potential improvements in human islet cell survival and insulin secretion could lead to improved glycemia, endowed by the BL001 treatment. Three independent experiments were performed with 3 different donors, two of which the BL001 regimen was for up to 4 weeks and 1 for up to 8 weeks (Fig. 7A and L). Although hyperglycemia persisted at either 4 and 8 weeks, blood glucose levels were lower in transplanted mice treated with BL001 for up to 8 weeks correlating with a higher survival rate as compared to vehicle treated mice at either 4 or 6 weeks, time at which all mice had to be euthanized due to pre-defined health criteria (Fig. 7B, C, M and N). Although not statistically significant, transplanted mice treated for 4 weeks with BL001 displayed a mild improvement in glucose tolerance as compared to vehicle mice (Fig. 7D and E). Human C-peptide was detected in BL001-treated mice, but not in vehicle-treated mice, at 4- and 8 weeks post-treatment, confirming some functionality of the transplanted human islet (Fig. 7F and O). In contrast, mouse C-peptide blood levels were marginally discernable in either BL001-or vehicle-STZ-treated mice, as compared to untreated mice, 4 and 8 weeks post-treatment (Fig. 7G and P). Both BL001- and vehicle-treated mice sacrificed at 4-weeks retained human islet xenotransplants with a similar number of INS and GCG expressing cells as well as CD4^+^ T-cell infiltration (Fig. 7H-K). The 8-week extended BL001-treatment preserved both the β- and α-cell mass along with CD4^+^ T-cell infiltration in xenotransplants correlating with improved glycemia while vehicle-treated mice that were sacrificed at 6 weeks (due to pre-defined health criteria) exhibited a significantly lower number of INS expressing cells and a higher number of GCG expressing cells as compared to BL001-treated mice (Fig. 7Q-T). Both vehicle and BL001-treated mice exhibited similar pancreatic islet insulin (INS) staining patterns, indicating that the mild recovery of glycemia in BL001-treated mice was not due to remnant endogenous islet β-cells but rather attributed to preserved islet graft mass and improved islet transplant outcomes conveyed BL001 administration (Fig. 7U). Taken together, these results indicate that BL001 favours the survival, engraftment, and function of a marginal mass of human islets in STZ-treated immunocompetent C57BL/6J mice.

**Fig 7.**
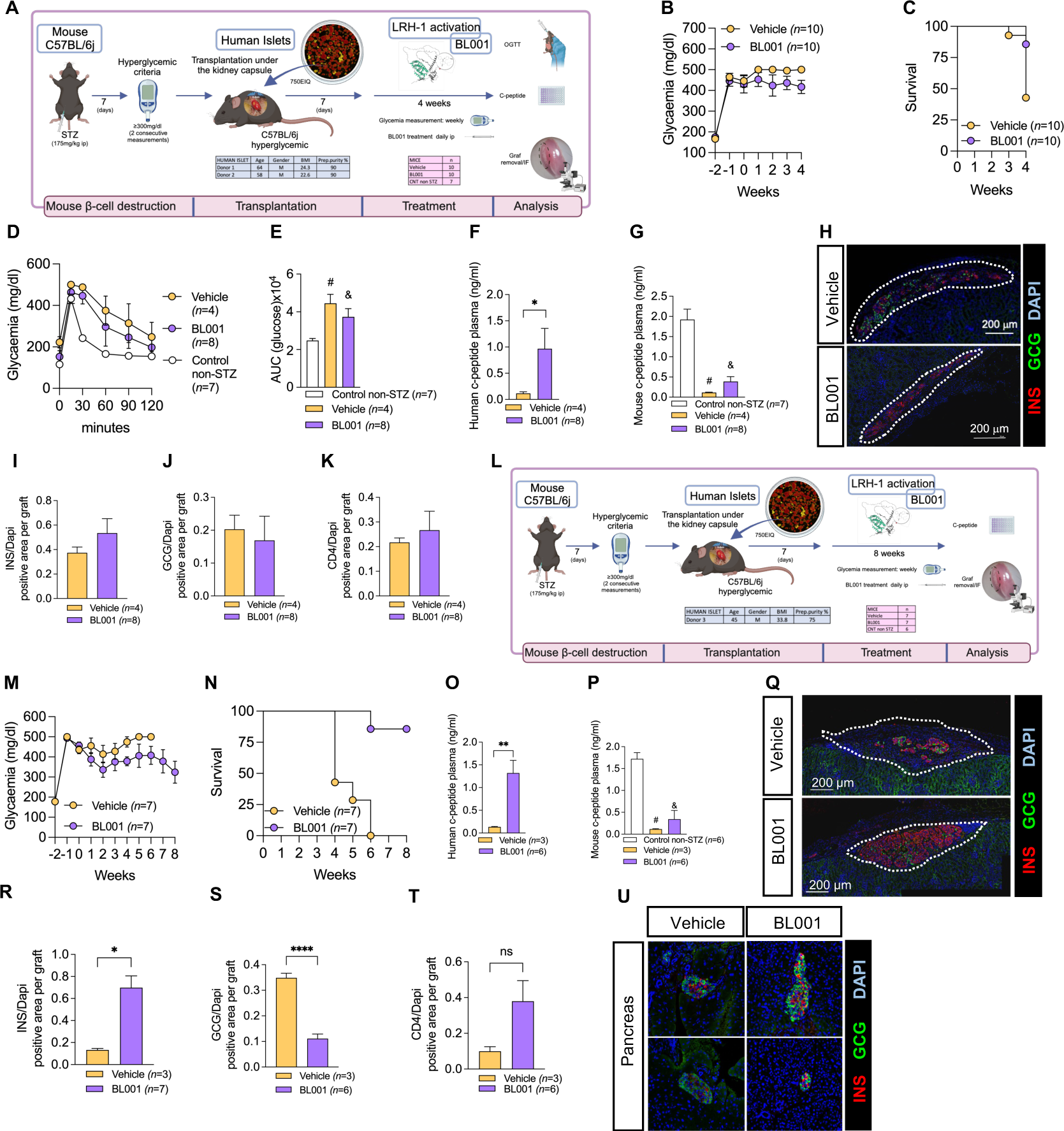
BL001 improves human islet graft survival and function in STZ-treated immunocompetent C57BL/6j mice. (**A**) Experimental design of the 4-week BL001 treatment post-xenotransplantation experiment. (**B**) Weekly measurement of non-fasting blood glucose and (**C**) Kaplan-Meier survival curve. (**D**) OGTT performed at 4 weeks post-BL001/Vehicle treatment. Mice were fasted for 6 hours before the OGTT. (**E**) Area under the curve (AUC) corresponding to the OGTT. Student t-test, # *p=0003* and & *p=0.0037* as compared to control non-STZ mice. (**F**) Human C-peptide plasma levels at 4 weeks post-BL001/Vehicle treatment. Data are presented as means ± SEM. *p<0.05 student t-test. (**G**) Mouse C-peptide plasma levels at 4 weeks post-BL001/Vehicle treatment. Data are presented as means ± SEM. Student t-test, # *p=002* and & *p=0.0003* as compared to control non-STZ mice. (**H**) Representative immunofluorescence images of kidney sections from mice euthanized at 4 weeks post-BL001/Vehicle treatment, displaying staining for insulin (INS), glucagon (GCG) staining along with nuclear DAPI staining. Quantitative analysis of (**I**) insulin (INS), (**J**) glucagon (GCG), and (**K**) CD4^+^ areas, normalized to the DAPI-positive area per graft, at 4 weeks post BL001 treatment. (**L**) Experimental design of the 8-week BL001 treatment post-xenotransplantation experiment. (**M**) Weekly measurement of non-fasting glycemia and (**N**) Kaplan Meier survival curve. (**O**) Human C-peptide plasma levels at 6-week vehicle-treated mice and 8-week BL001-treated mice. Data are presented as means ± SEM. **p<0.01 student t-test. (**P**) Mouse C-peptide plasma levels at 6-week vehicle-treated mice and 8-week BL001-treated mice. Data are presented as means ± SEM. # *p=0001* and & *p=0.*0002 Student t-test as compared to control non-STZ mice. (**Q**) Representative immunofluorescence images of kidney sections from mice at 6 weeks post-vehicle treatment and at 8 weeks post-BL001 treatment, displaying staining for insulin (INS), glucagon (GCG) staining along with nuclear DAPI staining. Quantitative analysis of (R) insulin (INS), (**S**) glucagon (GCG), and (**T**) CD4^+^ areas, normalized to the DAPI-positive area per graft, at 4 weeks post BL001 treatment. Data are presented as means ± SEM. *p<0.05 and ****p<0.0001 student t-test. ns, non-significant. (**U**) Representative immunofluorescence images of pancreas sections from mice at 6 weeks post-vehicle treatment and at 8 weeks post-BL001 treatment. displaying staining for insulin (INS), glucagon (GCG) staining along with nuclear DAPI staining.

**Fig. 8.**
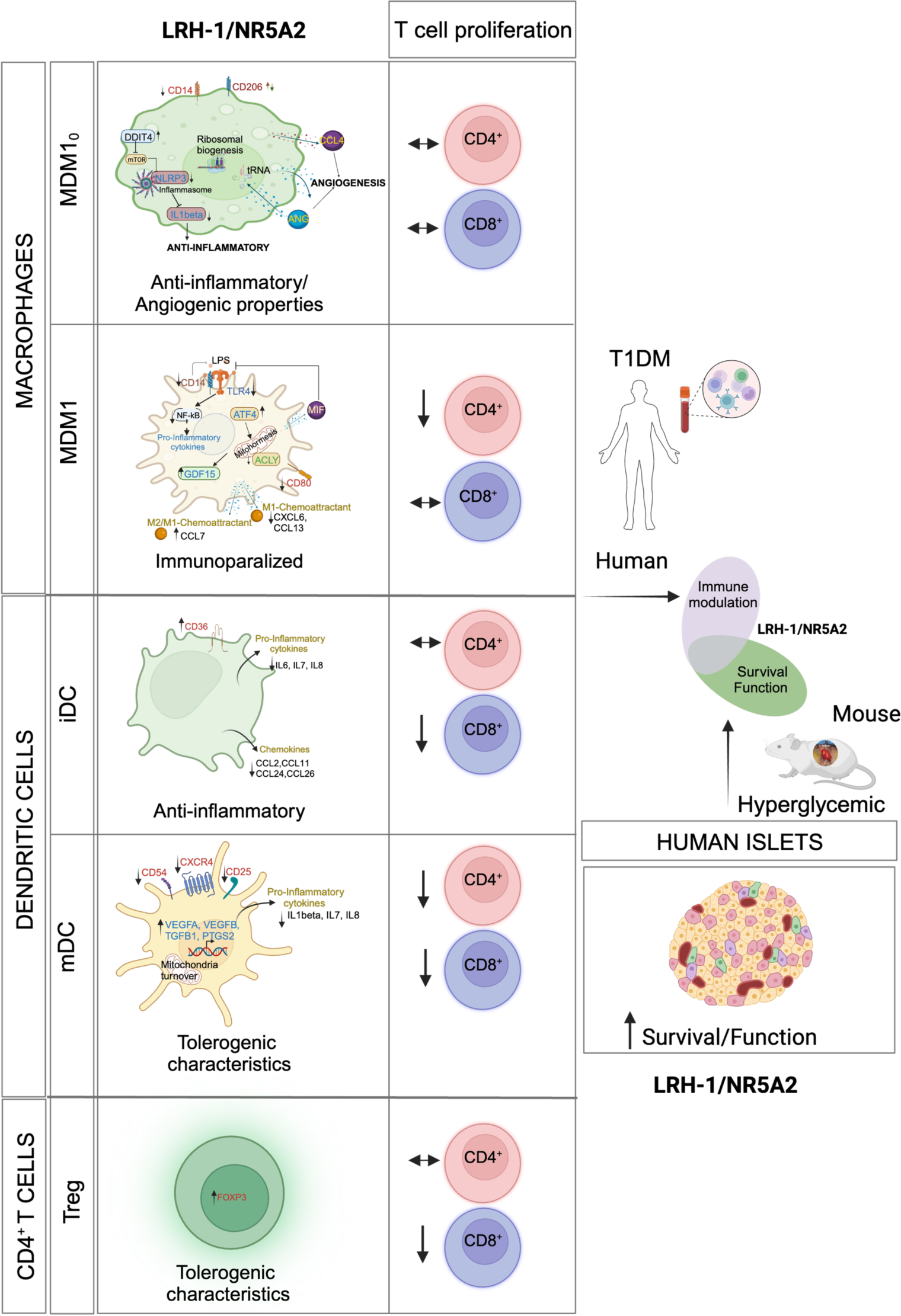
Cellular and molecular mechanism of action mediated by the pharmacological effect of LRH-1/NR5A2 in human cells. Proposed model of genetic and immune cell-tailored reprogramming induced by LRH-1/NR5A2 activation and its impact on T-effector cell proliferation. Improvement of human islet engraftment influenced by LRH-1/NR5A2 activation.

## DISCUSSION

Despite substantial research and development efforts across both academia and pharmaceutical sectors, a long-term pharmacological solution that can significantly reverse hyperglycemia in T1D has yet to be discovered. The complexity of T1D highlights the urgent need for a fundamental paradigm shift in our approach to understanding the diseasés intricate mechanisms, which involves a dynamic interaction between the immune system and pancreatic cells (*10*). This ‘symphony’ underscoring the pathogenesis of T1D (*66*), requires a conductor/target to orchestrate a coordinated resolution of the autoimmune attack/proinflammatory process, rather than suppression, leading to β-cell regeneration/survival (*10*). Herein, using MDMs, DCs, and T-cells isolated from individuals with established T1D and a combination of cell surface marker analysis, omics profiling, and functional studies, we demonstrate that the pharmacological activation of LRH-1/NR5A2 resolves the pro-inflammatory environment by: 1) locking MDM1_0_ in a naïve state while suppressing the pro-inflammatory M1-like phenotype of MDM1 through mitohormesis, 2) fostering a reprogramming of mDCs towards tolDCs by promoting mitochondrial turnover, thereby reducing CD4^+^ and CD8^+^ T-cells proliferation and 3) enhancing the expansion of CD4^+^/CD25^+^/FoxP3^+^ Tregs subpopulation, leading to diminished CD8^+^ T-cell proliferation. In parallel, our study demonstrates that the activation of LRH-1/NR5A2 can enhance the survival and function of human islets transplanted into diabetic immunocompetent mice. As such, we argue that LRH-1/NR5A2 fulfills all criteria of a ‘symphony conductor’ orchestrating resolution of the autoimmune attack coupled to improved islet cell survival and potential regeneration (*10, 21*). Given these promising results, it is conceivable that NR therapeutic targets like LRH-1/NR5A2 could also be effective in addressing other autoimmune diseases. To the best of our knowledge, our study is one of the first to demonstrate that a pharmacological compound not only imparts anti-inflammatory properties to fully activated MDM1s and mDCs of individuals with T1D but also promotes human engraftment and function in immunocompetent C57/BL6J, likely via tolerization as we previously demonstrated (*21*). Most studies have primarily concentrated on iDCs, evaluating the effects of compounds such as 1α,25-Dihydroxyvitamin D3 on DCs maturation and activation, but none have investigated their potential to transform fully mature DCs isolated from individuals with T1D into a tolerogenic phenotype (*67, 68*). Similarly, most human islet transplantation survival and functional studies have been carried out in immunodeficient mice to prevent rejection (*69*). Herein, we selected the STZ-treated C57/BL6J mouse model with the specific intention of expanding on our previous findings on BL001 tolerizing a pro-inflammatory environment, key for the sustained engraftment and long-term functionality of human islets (*21*). Consequently, our findings hold significant implications for human clinical studies, since reversing T1D will require addressing the chronic pro-inflammatory/autoimmune responses, which are primarily driven by macrophages and dendritic cells activated by islet self-antigens. These cells are pivotal in triggering T-cell proliferation and the subsequent β-cell destruction (*70*).

Our findings also emphasize the cell context-dependent nature of the genetic alterations and phenotypic outcomes induced by LRH-1/NR5A2 pharmacological activation. This is evidenced by the disparity in the molecular pathways regulated in BL001-treated MDM1s and mDCs, with many pathways suppressed in MDM1 but activated in mDCs. In MDM1s, LRH-1/NR5A2 activation suppressed multiple pathways implicated in inflammatory responses (15 out of 20), correlating with decreased expression of key cell surface markers such as CD14 and CD80, and reduced secretion of pro-inflammatory cytokines and chemokines. Interestingly, BL001 did not induce expression of M2-associated cell surface markers such as CD206, CD209, and CD200R suggesting that the action of LRH-1/NR5A2 activation is to neutralize/incapacitate the pro-inflammatory M1 phenotype rather than promoting a transition towards an M2-like phenotype. The concept of M1 incapacitation is further evidenced by the instatement of an immune-paralyzed phenotype via mitohormesis induced by BL001, which is consistent with previous studies reporting that LPS/IFNγ-mediated mitochondrial shutdown prevents IL-4 polarization of M1 towards an M2 phenotype (*71*). Additionally, BL001’s anti-inflammatory properties are highlighted by the inhibition of multiple enriched pathways related to inflammatory responses (systemic lupus erythematosus, influenza A, Epstein-Barr virus, among others), as well as pathways involved in either cell cycle or cellular senescence in BL001-treated MDM1_0._ These molecular events translated to the inhibition of CD4^+^ T-cells proliferation indicative of a resorbed Teff-mediated cell destruction. Notably, the mitohormesis-related factor GDF15, which was increased in BL001-treated T1D MDM1s, was shown to be expressed in a subset of macrophages and to promote muscle regeneration (*72*). Additionally, GDF15 expression was lower in T1D pancreas and its exogenous administration prevented insulitis progression in NOD mice and protected isolated human pancreatic islets. (*73*). Concurrently, pharmacological activation of LRH-1/NR5A2 led to upregulated expression of angiogenin and CCL4 in MDM1_0_ from T1D individuals. Both angiogenin and CCL4 induce angiogenesis, which is essential for supporting the formation of new blood vessels necessary for the delivery of nutrients and oxygen to growing tissues (*34, 74*). Taken together, these data emphasize the premise that BL001 favours a regenerative environment through increased expression of GDF15, angiogenin, and CCL4 from MDMs (*10, 21*). In this context, we have previously shown that CCL4 serum levels were increased in BL001-treated hyperglycemic RIP-B7.1 mice correlating with immune coupled α-to β-cell trans-differentiation (*21*). Angiogenin was also shown to cleave tRNAs, through its ribonuclease activity, thereby transiently inhibiting translation in cells under stress conditions to quickly adapt to their new environment (*75, 76*). Accordingly, the top two activated enriched pathways in BL001-treated T1D MDM1_0_ were aminoacyl-tRNA biosynthesis and ribosome, evidencing an autocrine effect of angiogenin to alleviate stress-induced molecular reprogramming.

Pharmacological activation of LRH-1/NR5A2 reprogrammed T1D mDCs towards a tolerogenic-like phenotype. This is evidenced by the activation of key molecular pathways inducing tolerance, such as OXPHOS, PPAR signalling, lysosome, and phagosome, as well as a specific subset of tolerogenic-associated genes, including PTGS2. Previously, we demonstrated that PTGS2 and its product, prostaglandin E_2,_ endow a tolerogenic phenotype to DCs after efferocytosis and prevent cytokine-induced apoptosis in BL001-treated islet β-cell (*22, 23, 77*). Consistent with a shift towards a tolerogenic phenotype, levels of several cell surface markers associated with mDCs (CXCR4 and CD54) were decreased, as did the secretion of IL-7 in cells treated with BL001. Of high relevance, a recent clinical trial assessing the efficacy of the humanized anti–IL-7R monoclonal antibody RN168 in subjects with T1D revealed a significant decline in CD4^+^ and CD8^+^ effector and central memory T-cells along with an increase in Tregs (*78*). BL001 treatment of mDCs from T1D donors, similarly suppressed the proliferation of both CD4^+^ and CD8^+^ T-cells suggesting a potential role of IL-7 in this process. This premise is further substantiated by the decreased levels of IL-7 in BL001-treated iDCs, which also impeded CD8^+^ T-cell expansion. Contrary to our initial hypothesis and based on RNAseq data revealing BL001-mediated activation of the PPAR and OXPHOS genetic pathways, with no net differences in mitochondrial respiration experiments, we advocate that the phenotypic switch induced by LRH-1/NR5A2 activation is likely mediated by enhanced mitochondrial turnover. The latter is supported by a decreased number of mDCs with dysfunctional mitochondria correlating with omics data evidencing mitochondrial biogenesis. Previous studies have demonstrated that mitophagy/autophagy contribute to maintaining an anti-inflammatory phenotype in immune cells (*79*).

Tregs play a key role in sustaining peripheral tolerance, and their dysfunction have been reported in individuals with T1DM (*80, 81*). Consequently, clinical trials involving autologous Tregs reinfusion following *in vitro* expansion, have shown a transient increase in these cells, correlating with enhanced islet survival and C-peptide levels, rendering Tregs as a promising cell-based therapy for T1DM (*82, 83*). Nevertheless, these cells were found to be short-lived in transfused patients, and a subsequent clinical trial that combined Tregs with IL-2 to improve survival, also resulted in an increase in cytotoxic T-cells (*82, 84*). Herein we show that BL001 stimulates the expansion of a CD4^+^/CD25^+^/FoxP3^+^ Tregs subpopulation within PBMCs as well as in CD4^+^ T-cells corroborating our previous findings in mice (*21*). The decrease in FoxP3 expression upon LRH-1/NR5A2 silencing in PBMCs directly links the observed immunomodulatory effects of BL001 to the activation of the nuclear receptor. Thus the pharmacological activation of LRH-1/NR5A2 may offer an appealing pharmacological alternative to cell therapy.

Consistent with its anti-inflammatory properties observed in both mice (*21*) and human immune cells, we demonstrate that the therapeutic administration of BL001 to hyperglycemic immunocompetent mice xenotransplanted with human islets favours graft implantation and functionality. While our prior work demonstrated that short-term BL001 treatment could protect transplanted human islets in mice against rejection, long-term engraftment, and function, as evaluated by circulating human C-peptide was not assessed in that study (*21*). Our findings have substantial implications for future immunomodulatory therapies in combination with the transplantation of either human islets or induced pluripotent stem cell (hIPSC)-derived islet-like organoids, which are also susceptible to immune rejection. Current strategies to avoid graft rejection encompass cell encapsulation and/or aggressive immunosuppressant regimens, often resulting in secondary complications such as kidney failure (*85–87*). Alternatively, hIPSC-derived islet-like organoids are being genetically modified to successfully evade the autoimmune attack in pre-clinical mouse models (*88, 89*). Nonetheless, these approaches have yet to reach ethical approval for clinical trials. Pharmacological activation of LRH-1/NR5A2 could circumvent these caveats encountered by cell therapy by attenuating the pro-inflammatory environment-without suppressing the general immune system-while increasing β-cell survival and performance (*22, 23*).

Overall, our findings introduce a new dimension in T1D therapies, where the pharmacological activation of LRH-1/NR5A2 instructs the genetic and metabolic reprogramming of human monocyte-derived macrophages, dendritic cells, and T-cells from individuals with T1D to support an anti-inflammatory and regenerative environment. This resolution of the autoimmune attack also promotes islet survival and function (**Figure 10**). Although our study was intentionally targeted to adults with long-standing T1D, the immunomodulatory benefits of BL001 should also be assessed in either young or older individuals who have recently been diagnosed with T1D in which the strong autoimmune attack may be more resilient to any drug treatment. While the mode of action of LRH-1/NR5A2 agonistic activation has been elucidated in distinct populations of macrophages and dendritic cells as well as its impact on T-cells *in vitro*, the net physiological outcome endowed by BL001 in a more complex milieu comprised of several interacting immune cell types along with islets have yet to be evaluated. Furthermore, whether regenerative factors secreted by immune cells exposed to BL001 will promote human β-cell regeneration remains to be assessed. Modelling T1D *in vitro* will address these limitations (*90*).

## MATERIALS AND METHODS

### Study design

The primary objective of the study was to validate the translational potential of BL001, a small and specific chemical agonist of LRH-1/NR5A2 conveying immunomodulatory effects in mouse models of T1D, as a prospective therapeutic avenue for individuals with T1D. Towards this goal, we assessed the capacity of BL001 to promote an anti-inflammatory phenotype to immune cells obtained from individuals with long-standing T1D and delineated the underlying cellular and molecular mechanisms conveying these benefits. Samples of 30-50 mL peripheral blood were collected by venipuncture from 68 individuals with T1D in BD Vacutainer Sodium Heparin tubes (BD Biosciences, San Jose, CA, USA). Inclusion criteria were: 20-55 years of age with equal sex representation, body mass index 18-30 kg/m^2^ and an evolution of the disease longer than 5 years. Exclusion criteria were: being under immunosuppressive or anti-inflammatory treatment, or undergoing pregnancy or breastfeeding. Patients were informed of the procedure and signed a written consent prior to blood extraction. The collection and processing of personal and clinical data from included subjects were limited to those data necessary to achieve the objectives of the study, and the data collected was processed with adequate precautions to ensure confidentiality and compliance with applicable data privacy protection laws and regulations. The Clinical/Investigation Ethical Committees of the Hospital Germans Trias i Pujol and the University Hospital Virgen Macarena and Rocio approved all experiments with human islets and blood-derived cells (PI-19-142, GAUTHIER-201306722 and 2057-N-22) which were been conducted following the principles outlined in the Declaration of Helsinki for human research. We applied a normalization approach for the analysis of cell surface markers, cytokine profiling as well as T-cell proliferation due to the inherent biological variability observed in samples derived from different individuals. This step was crucial to account for inter-individual differences and to ensure that our results reflect genuine biological effects rather than variability in baseline expression levels. It allowed us to accurately compare the relative changes induced by BL001 across all samples. For xenotransplantation experiments, C57BL/6J mice were purchased from Janvier Labs (Saint-Berthevin Cedex, FR). Due to the restricted number of cells isolated from each blood sample, individual preparation of macrophages and DCs was used in specific subsets of experiments (flow cytometry, cytokine profiling, omics, etc.). Mice were housed in ventilated plastic cages under a 12-h light/dark cycle and given food and water *ad libitum*. We opted for the C57BL/6J strain based on our prior work, which showed that BL001 could induce an anti-inflammatory and tolerogenic environment in this specific mouse background (*21*). The rationale was to circumvent the need for re-characterizing the immune cell profile in a humanized mouse model and focusing primarily on long-term human islet viability and function. Mouse experimentations were approved by the Andalusian Ministry of Agriculture, Farming, Fish and Sustainable Development (08/07/2019/120). Animal studies were performed in compliance with the ARRIVE guidelines (*91*). Human islets were either obtained from the Alberta Diabetes Institute IsletCore Laboratory (CA) (*n=2* donors) or from ECIT-San Raffaele Scientific Institute, Milan (IT) (*n=1* donor) (see Table 1 for clinical characteristics). The sample size for these *in vivo* studies to reach statistical significance was not precalculated because the survival of islet grafts was previously unknown. Mice were treated with streptozotocin and then transplanted with human islets before being randomly allocated to either the vehicle-or BL001-treated group. Although the xenotransplantation of islets and BL001 treatment were not blinded to the investigator, the subsequent analysis of blood samples and grafts were performed by blinded investigators.

### Peripheral blood mononuclear cell (PBMC) and monocyte isolation

Peripheral blood mononuclear cells (PBMCs) were isolated using either Ficoll Paque or Histopaque 1077 density gradient centrifugation. The PBMC layer was extracted and washed with PBS. Cells were resuspended in PBS, 2% FBS, 1mM EDTA buffer and monocytes were then isolated using the magnetic EasySep Human CD14^+^ Selection kit (STEMCELL Technologies, Vancouver, BC, Canada) following the manufacturer’s instructions. When the purity of CD14 marker in the selected fraction was greater that 90%, monocytes were cultured in 24-well plates (Labclinics, Barcelona, Spain) at a concentration of 10^6^ cells/ml in either X-VIVO 15 media (Lonza, Basel, Switzerland) or RPMI 1640 media (ThermoFisher Scientific), both supplemented with 2% male AB human serum (Biowest Nuaillé, France), 100 IU/ml penicillin (Sigma-Aldrich and Normon SA, Madrid, Spain), 100 µg/ml streptomycin (Sigma-Aldrich, Madrid, Spain). The negatively selected fraction of PBMCs was cryopreserved in FBS (ThermoFisher Scientific) with 10% dimethylsulfoxide (Sigma-Aldrich, Saint Louis, MO, USA) at a 10-20X10^6^ cells/ml and stored for later use.

### Monocyte-derived macrophages (MDMs) and dendritic cells (DCs)

Purified CD14^+^ monocytes were derived into either macrophages (MDM) or dendritic cells (DCs). For MDM, cells were treated with 1,000 IU/ml rhGM-CSF (Prospec, Rehovot, Israel) for 3 days to generate naïve/primed MDM1_0_ and subsequently with a cocktail containing 20ng/ml of INFγ (Immunotools, Friesoythe, Germany) and 10ng/ml of LPS (Sigma-Aldrich) to promote the pro-inflammatory MDM1 phenotype. Alternatively, monocytes were treated with 1000 IU/ml rhIL-4 and 1000 IU/ml rhGM-CSF (Prospec, Rehovot, Israel) for 6 days to obtain DCs. Media and cytokine stimuli were replenished at day 4. Dendritic cells were either cultured with 20 μg/ml human insulin (Sigma-Aldrich) to obtain immature DCs (iDCs) or adding a cytokine cocktail (CC) consisting of tumor necrosis factor (TNF)α (1000 IU/ml, Immunotools, Friesoythe, Germany), IL1β (2000 IU/ml, Immunotools) and Prostaglandin E2 (PGE2, 1 μM, Cayman Chemical, Ann Arbor, MI, USA) to obtain mature DCs (mDCs). All derived cell types were maintained at 37°C in the presence of 5% CO_2_.

### CD4^+^ T-cell isolation

The CD4^+^ T-cell subpopulation was isolated from the CD14^-^ fraction of PBMCs using the Easysep human CD4^+^ T-cell isolation kit (Stemcell Technology) as per the instructions of the manufacturer. CD4^+^ T-cells were cultured in RPMI 1640 media (ThermoFisher Scientific), supplemented with 2% male AB human serum (Biowest Nuaillé, France), 100 IU/ml penicillin (Sigma-Aldrich and Normon SA, Madrid, Spain), 100 µg/ml streptomycin (Sigma-Aldrich, Madrid, Spain). Subsequently, CD4^+^ T-cells were activated using the TransAct reagent (Milteny) for 24h before a single dose of 10 µM BL001 was added to the culture media.

### Human islet culture

Human islet preparations were washed, handpicked and subsequently maintained in CMRL-1066 (ThermoFisher Scientific) containing 5.6 mM glucose, and supplemented with 10% FCS, 100 Units/ml penicillin, 100 μg/ml streptomycin and 100 μg/ml gentamycin (all purchased from Sigma-Aldrich).

### BL001 treatment

Three consecutive doses of 10 µM BL001 was added to the culture media of PBMCs, MDM1_0_, MDM1, iDCs and mDCs at 0, 24 hours and 30 minutes prior to cell analysis or processing.

### Flow Cytometry

Subpopulations of immune cells were characterized by flow cytometry (FACSCalibur and FACSAria I, BD Biosciences, Madrid, Spain), using either Zombie Violet 421 (Biolegend) or 7aad (BD Biosciences) for viability assessment and the following antibodies for 1) MDMs phenotyping: CD14 APC/FITC, CD80 PE, CD86 APC, CD163 PE-Vio615/BV421, CD200R PE, CD206 FITC/BV711, and CD209 FITC and 2) DCs phenotyping: CD11c APC, CD25 PE, CD86 FITC, HLA class I FITC, HLA class II FITC, CD14 PE and CD40 APC, CD36 APCCy7, TIM4 APC, αvβ5 integrin PE, CD54 PECy7, CXCR4 APCCy7, CCR2 APC, DC-SIGN-APC, PD-L1 PECy7 and CCR7 PECy7 (Table 2). Regulatory T cells were evaluated using the REAfinity™ Treg Phenotyping Kit (Table 2). T helper cell phenotyping was performed using CD4 Viogreen, CD183 PE, CD194 APC, CD196 BV423. Data were analyzed using either the FlowJo V9 (Tree Star) or the FCS Express softwares (De Novo Software, Pasadena, USA).

**Table 2:**
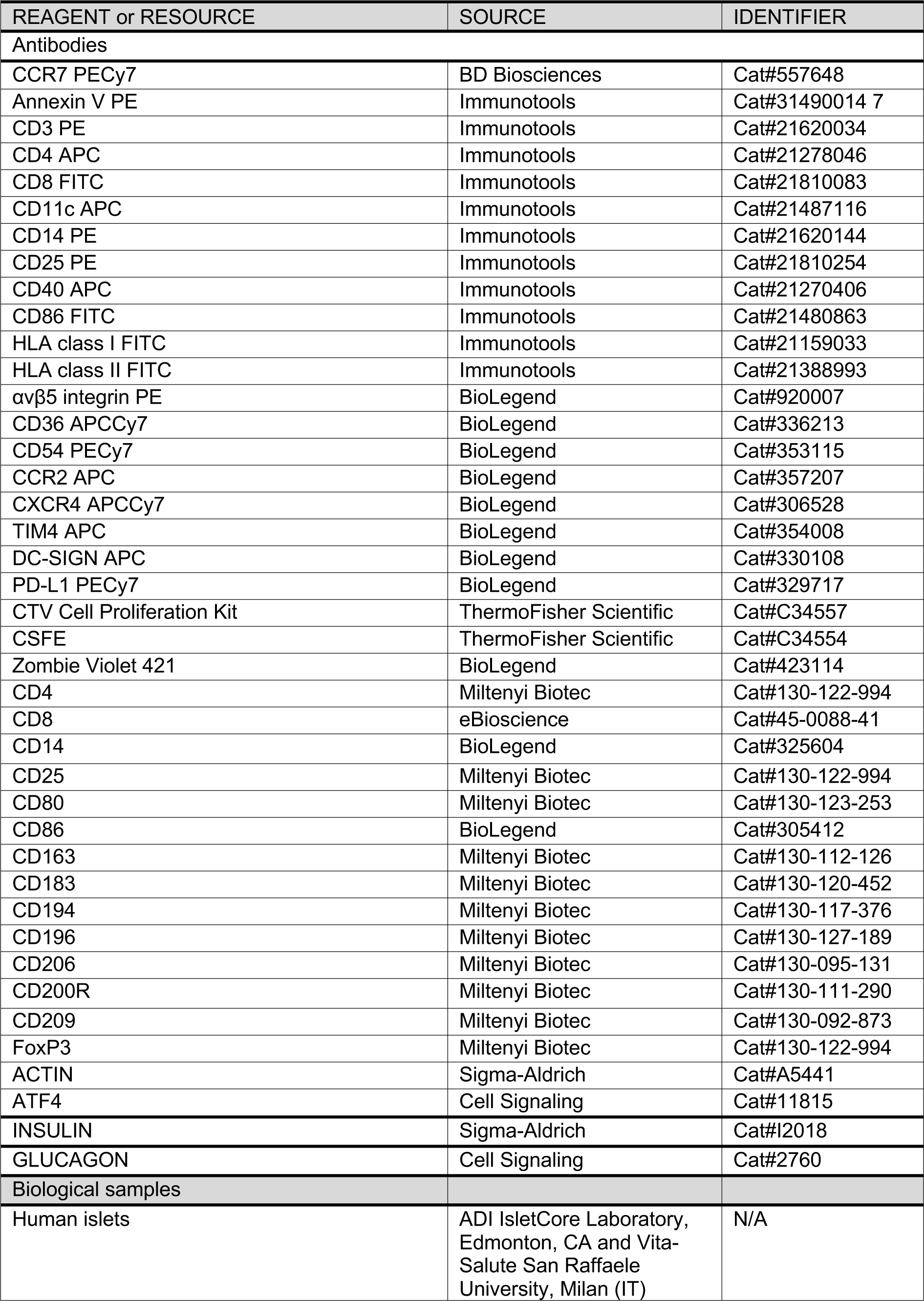
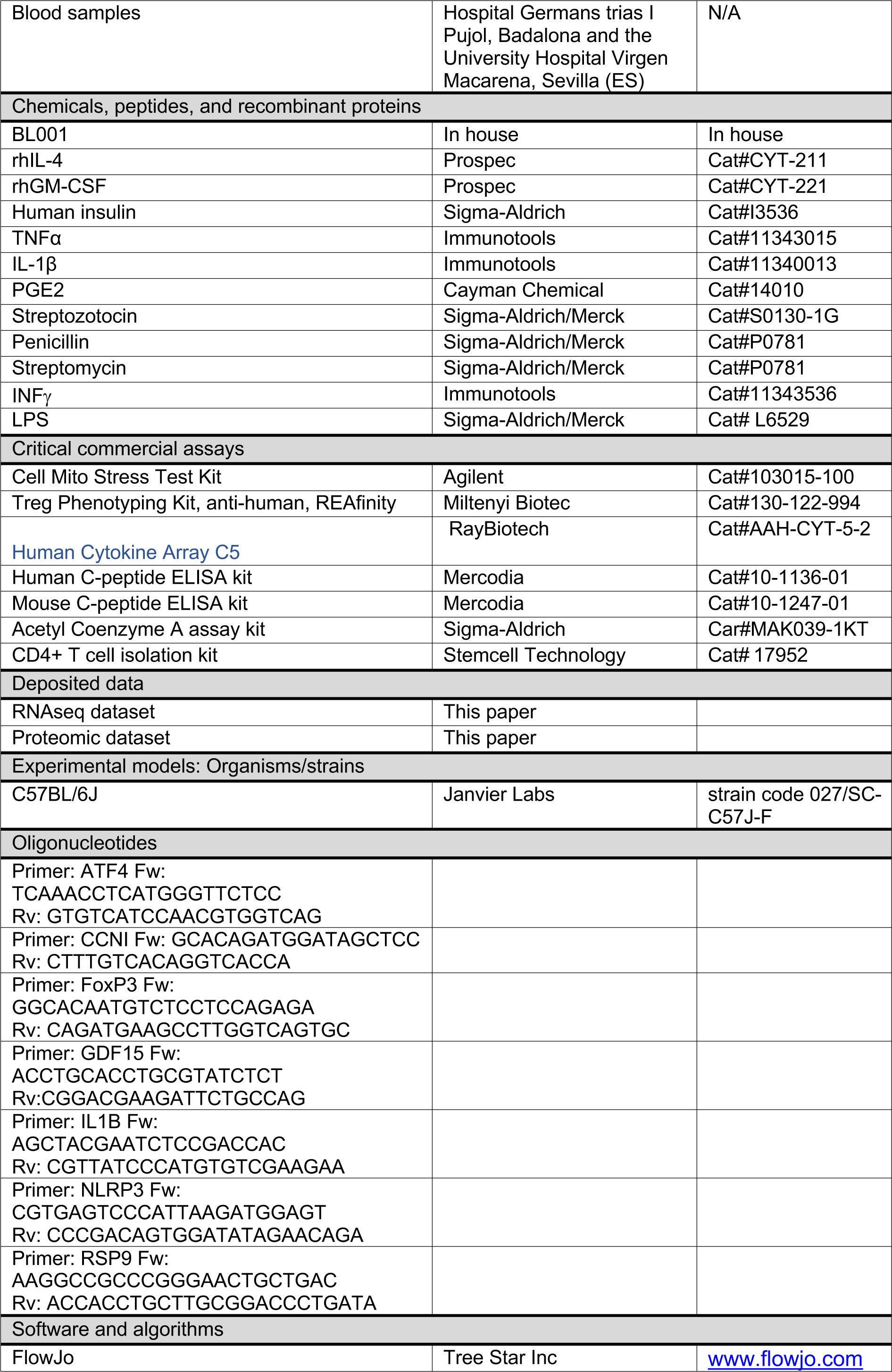
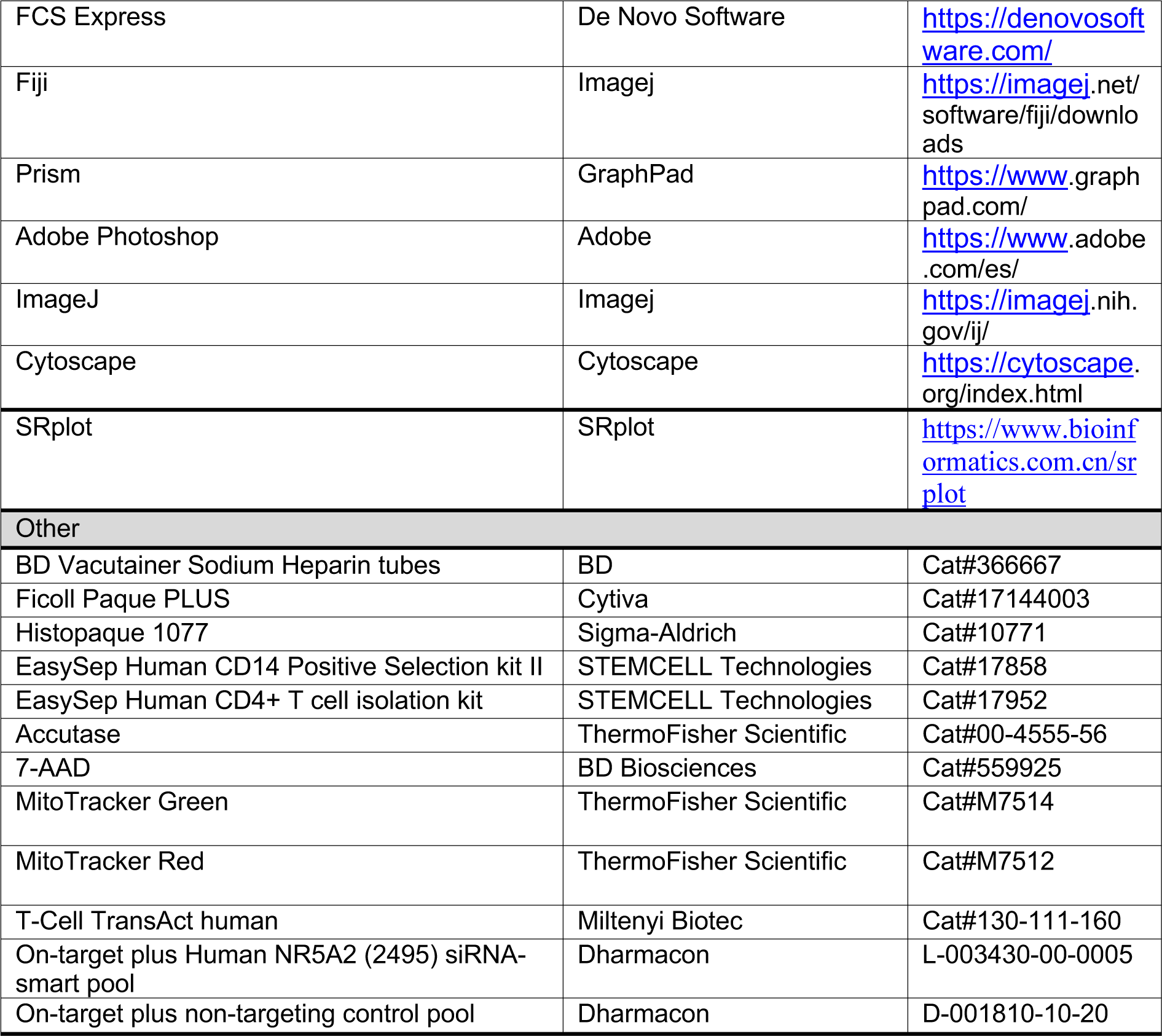
Key resources.

### Cytokine profiling

Media was collected from MDMs and DCs, treated or not with BL001, and analyzed using the RayBiotech Human Cytokine Array C5 that interrogates 80 cytokines simultaneously (RayBiotech, Norcross, USA). Membranes were scanned and analyzed using the ImageJ/Fiji software. Results were then normalized using the internal controls of the membranes, and the relative signal intensity are represented.

### Autologous T Cell Proliferation Assays

Immune cells cultured in each condition were co-cultured with autologous T lymphocytes to determine their capacity to induce T cell proliferation. Briefly, PBMCs from the same donor were thawed to be stained with the CellTrace Violet (CTV) or CFSE Cell Proliferation kit (ThermoFisher Scientific) following the manufacturer’s instructions. After staining, cells were suspended in either RPMI 1640 or X-VIVO 15 complete media at a final concentration of 10^6^ cells/ml, and 100,000 cells were plated in 96-well round bottom plates (Labclinics). T cell activation was induced using a cocktail of CD3 and CD28 antibodies (T Cell TransAct, Miltenyi biotech). Stained and activated PBMCs were co-cultured with either MDMs, DCs or CD4^+^ T-cells subject to the various experimental conditions at a 10:1 or 2:1 ratio (10^5^ PBMCs:10^4^ DCs; 2×10^4^ PBMCs: 10^4^ MDMs; 2×10^4^ PBMCs: 10^4^ CD4^+^ T-cells) in triplicates. After up to 6 days of co-culture in the incubator at 37 °C and 5% CO2, cells were washed with 150 µL PBS per well at 400×g for 5 minutes and incubated for 20 minutes at 4 °C with a staining mix containing CD3 PE, CD4 APC/Viogreen and CD8 FITC/PerCP-Cy5.5 staining and 7-AAD/Zombie Violet 421. Cells were then washed in PBS and T-cell proliferation was analyzed by flow cytometry. Data were analyzed using the FlowJo software (Tree Star Inc.).

### Transcriptome profiling

Total RNA was isolated from human primary immune cells using the RNeasy Plus Micro Kit (Qiagen). RNA integrity number (RIN) values were evaluated using Bioanalyzer® 2100 (pico assay) and their profiles were accepted for preparing libraries for NGS (RIN >8.40). RNA-seq libraries were performed by the Genomic Facility at CABIMER, using the kit Illumina Stranded TOTAL RNA preparation RIBO-ZERO PLUS and sequenced on a NovaSeq 6000 platform with an average of 30 million reads per sample. Reads were mapped and quantify using Salmon (version 1.5.0) with default parameters to the human transcriptome (assembly GRCh38) downloaded from GENCODE genome with default parameters. Differential expression was analysed using Deseq2 using a paired sample design. Gene set enrichment analysis was performed using clusterProfiler (version 4.0).

### Proteomic profiling

Monocyte derived macrophages and DCs were lysed in 8 M urea/10 mM HEPES (pH 8.0). Samples were then supplemented with 1 mM Dithiothreitol (DDT) and incubated 30 min at RT at which point 5 mM chloroacetamide (CAA) was added and further incubated for 30 min at RT. Finally, samples were supplemented with 5 mM DTT and incubated for another 30 min at RT. Five hundred ng of Lys-C (Wako) was added to the samples and incubate for 5 h. Subsequently, three volumes of 50 mM ABC were added to dilute the urea to a final concentration of 2M. A second round of digestion with 500 ng of Trypsin (Promega) was performed overnight. Resulting peptides were desalted with C18 StageTips (*92*) and analyzed by label-free quantitative mass spectrometry-based proteomics using a Q-Exactive Orbitrap mass spectrometer (Thermo Scientific, Germany) equipped with an EASY-nano liquid chromatography 1000 system (Proxeon, Odense, Denmark). The Q-Exactive was coupled to a fritted fused silica capillary (25 cm L x 75 µm ID x 360 µm OD, CoAnn Technologies, Richland, US) in-house packed with 1.9 µm C18 beads (Dr. Maisch, cat:r119.aq, Ammerburch-Entringen, Germany).

Peptides were separated by liquid chromatography using a 2h gradient from 2% to 95% solvent B (water/acetonitrile/formic acid at a ratio of 20/80/0.1) and followed by 0 % acetonitrile in 0.1% formic acid for chromatography column re-conditioning. The mass spectrometer was operated in positive polarity mode at 2.3 kV with the capillary heated to 250 °C. Data-dependent acquisition mode with a top 10 method was used to automatically switch between full-scan MS and MS/MS scans. Full scan MS spectra were acquired at a resolution of 70,000, an automatic gain control (AGC) target value of 3 × 10^6^, and a scan range of 400–1,400 m/z. Precursors were fragmented by Higher-Collisional Dissociation (HCD) with a normalized collision energy of 25. Tandem mass spectra (MS/MS) were recorded with a resolution of 17,500 and an AGC target value of 1 × 10^5^. Precursor ion masses of scanned ions were subsequently dynamically excluded from MS/MS analysis for 60 sec and only precursors with a charge state of 2-6 triggered MS/MS events. Maximum injection times for MS and MS/MS were 20 and 60 ms, respectively.

Mass spectrometry RAW files was analized using MaxQuant (v2.1.3.0) (*93*). Search was performed against an in-silico digested UniProt reference proteome for Homo sapiens including canonical and isoform sequences (29th August 2022). Database searches were performed according to standard settings with the following modifications: Digestion with Trypsin/P was used, allowing 3 missed cleavages. Oxidation (M), acetyl (protein N-term) were allowed as variable modifications with a maximum number of 3. Label-Free Quantification (LFQ) was enabled, not allowing Fast LFQ. Matching between runs was enabled with a match time window of 0.7 min and an alignment time window of 20 min.

Output from MaxQuant Data were exported and processed for statistical analysis in Perseus (*94*). LFQ intensity values were log2 transformed and potential contaminant and proteins either identify by site or only reverse peptides were removed. Samples were grouped in experimental categories and proteins not identified in every replicate in at least one condition were removed. Missing values were imputed using normally distributed values with 0.3 width and 1.8 down shift separately for each column. After imputation results were exported into in MS Excel and paired t-test were performed. Volcano plots were constructed for data visualization using the VolcaNoseR web (*95*) (https://huygens.science.uva.nl/VolcaNoseR2/). We explored the protein-protein interaction (PPI) network (PPI) among differentially expressed proteins in MDMs using the online search tool for the retrieval of interacting genes/proteins (STRING v11) that comprise a database of known and predicted protein-protein interactions (*96*). Networks were then visualized in the Cytoscape software (version 3.9.1), using the yfile circular layout algorithm (*97*). Alternatively, the SRplot webserver platform was used to interrogate GO datasets for differentially expressed proteins in DCs (*98*).

### Acetyl Coenzyme A measurement

MDM1, treated or not with BL001, were washed in PBS and then lysed in 1M Perchloric acid. The lysate was centrifuged at 10,000g for 10 min at 4°C. The deproteinized supernatant was neutralized with potassium bicarbonate and the neutral pH confirmed with Whatman® indicator papers. The Acetyl CoA content was then determined using the Acetyl coenzyme A assay kit as per the instructions of the manufacturer (Sigma-Aldrich). An Acetyl-CoA standard curve was plotted in the range of 0–100 pmol. Fluorescence was measured using a Varioskan Flash spectrophotometer. For normalization by protein content, the protein pellets were solubilized first in 0.2 N NaOH and then solubilized in a buffer containing 7M Urea, 2M ThioUrea, Tris 50mM ph 8.8, as previously described (*99*). The protein content was determined using the Quick Start Bradford Protein assay (Biorad). Acetyl CoA content was then normalized to protein content.

### Mitochondrial bioenergetic and fitness

Mitochondrial bioenergetics of MDMs and DCs were measured using the Seahorse XF Cell Mito Stress Test Kit and the XF24 Extracellular Flux Analyzer (Agilent), as previously described (*100*). Briefly, after overnight culture, cells were washed and replenished with Seahorse assay media (Seahorse Bioscience), supplemented with 1 mM pyruvate and 2 mM glutamine. Where applicable, 10 µM BL001 and 1 mM palmitate conjugated to 0.17 mM BSA (150 mM NaCl, pH 7.2), were also added to designated wells. Plates were incubated in a CO_2_-free incubator at 37 °C for 1 h to allow temperature and pH equilibration, after which oxygen consumption rate (OCR) was measured in the XF24 Extracellular Flux Analyzer over a period of 95 min. Mitochondrial processes were examined through sequential injections of oligomycin (4 μM) at min 21, carbonyl cyanide 4-(trifluoromethoxy) phenylhydrazone (FCCP; 2 μM) at min 45, 5 μM antimycin A/Rotenone at min 78. At the end of the measurement, cells in each well were counted using the Scepter™ 2.0 Cell Counter to normalize the data. Only the oxygen consumption rate (OCR) for the basal respiration rate was considered for which the area under the curve (AUC) is presented in pertinent figures. Mitochondrial fitness as assessed by the number of non-functional mitochondria in mDCs treated or not with BL001, was determined by flow cytometry using the probes MitoTracker Green that binds covalently to mitochondrial proteins, thus providing an assessment of mass and MitoTracker Red that is taken up by polarized mitochondria thus gauging function. Non-functional mitochondria were then determined by the ratio of MitoTracker Green^High^ over MitoTracker Red^low^ (*65*). Fluorescent images were acquired using a Leica TCS SP5 confocal microscope.

### siRNA silencing

On-target plus NR5A2 siRNA-smart pool or control on-target plus non-targeting pool were used for silencing studies in PBMCs (Table 2) as previously described (*21*). RNA was extracted 48 hours post transfection.

### RNA extraction and quantitative real-time PCR

Total RNA from was extracted using the RNeasy Micro Kit (Qiagen, Madrid, SP). Complementary DNA using 0.1 to 1 µg RNA was synthesized using the Superscript III Reverse Transcriptase (Invitrogen-Thermo Fisher Scientific, Madrid, Spain). The QRT-PCR was performed on individual cDNAs using SYBR green (Roche) (*21*). Gene-specific primers were selected using a human housekeeping gene database (HRT Atlas v1.0 database) (*101*) and the sequences are listed in Table 2. Expression levels were normalized to various reference genes including CCNI, ACTIN and RSP9. The relative gene expression was calculated using the standard curve-based method (*102*).

### Western blot

MDMs were disrupted in a RIPA lysis buffer containing protease (P8340) and phosphatase inhibitors (P0044, P5726). Western blots were performed according to standard methods (*100*). Antibodies employed are provided in Table 2.

### Xenotransplantation

Human islet transplantations were performed as previously described (*21*). Briefly, 8-week-old immune-competent C57BL/6J male mice were treated with a single dose of 175 mg /kg b.w. streptozotocin (STZ) prepared in 0.01 M sodium citrate at pH 4.5 to induce hyperglycemia. One week later, mice were anesthetized via an intraperitoneal (*i.p.*) injection of 100 mg/Kg ketamine and 10 mg/Kg xylazine and 750 human islet equivalents (IEQ) were transplanted under the kidney capsule using a PE50 tubing connected to a 25 µL gauged Hamilton syringe. Mice were then *i.p.* injected, or not, with 10 mg/kg body weight (b.w.) BL001. Circulating glucose levels were measured from tail vein blood samples using an Optium Xceed glucometer (Abbott Scientifica SA, Barcelona, Spain). An oral glucose tolerance test (OGTT) was performed at 5-weeks post-transplantation as previously described (*103*). Upon termination of the experiment, animals were sacrificed and transplanted kidneys extracted, fixed and embedded for further histological analysis. The entire kidney was sectioned and insulin/glucagon co-immunostaining as well as CD4 was performed at every 15^th^ slice, corresponding with an interval of ∼ 75-150 µm which matches the median size of the majority of islets.

### Immunofluorescence analyses

For immunostaining, kidneys were fixed overnight in 4% paraformaldehyde at 4°C. Tissues were dehydrated, paraffin embedded, and sectioned at 5 μm thickness. Immunostaining was then performed overnight at 4°C using a combination of primary antibodies (Table 2) in PBS 1% BSA 0.2% TritonX100. Subsequently, secondary antibodies (Table 2) were incubated for 1 hour at room temperature in PBS 0.2% TritonX100. Nuclei were stained with 0.0001% of 4′,6-diamidino-2-phenylindole (DAPI, Sigma-Aldrich) and cover slips were mounted using fluorescent mounting medium (DAKO). Epifluorescence microscopy images were acquired with a Leica DM6000B microscope and z-stack images were acquired using a Confocal Leica TCS SP5 (AOBS) microscope.

### Morphometric analyses

Images of kidney sections were automatically acquired using the Thunder imager software and processed using the Photoshop, ImageJ and FIJI softwares.

## QUANTIFICATION AND STATISTICAL ANALYSIS

Data are presented as the mean ± SEM. Student’s t-tests were used as described in figure legends. *p* values less than or equal to 0.05 were considered statistically significant. Statistical analyses were performed using the GraphPad Prism software version 8 (GraphPad Software, La Jolla, USA).

## SUPPLEMENTARY MATERIALS

Supplemental information can be found online at XXX

## ACKNOWLEDGMENTS

We thank Dr. Maria José Quintero from the Cytometry Core Facility and the team from the Genomic Core Facility of CABIMER as well as Pedro Antonio Soriano Fernandez and Noelia García Rodríguez for their excellent technical support. Special thanks to all donors and to the Biobank Node of the Hospital Virgen Macarena (Biobanco del Sistema Sanitario Público de Andalucía) which is part of the Spanish National biobanks Network (PT20/00069) supported by Instituto Carlos III and FEDER funds. We acknowledge the support of the Basic Experimentation in Diabetes study group of the Spanish Association of Diabetes as well as positive discussion through the Consolidated Research Group 2021 SGR 00002, AGAUR, Generalitat de Catalunya. Human islets for research were provided, in part, by the Alberta Diabetes Institute IsletCore at the University of Alberta in Edmonton (www.bcell.org/adi-isletcore) with the assistance of the Human Organ Procurement and Exchange (HOPE) program, Trillium Gift of Life Network (TGLN), and other Canadian organ procurement organizations. Islet isolation was approved by the Human Research Ethics Board at the University of Alberta (Pro00013094). All donors’ families gave informed consent for the use of pancreatic tissue in research.

## FUNDING

The authors are supported by grants from the Consejería de Salud y Consumo, Fundación Pública Andaluza Progreso y Salud, Junta de Andalucía (PI-0727-2010 and PI-0001-2020 to B.R.G), the Consejería de Economía, Innovación y Ciencia (P10.CTS.6359 to B.R.G and DOC_00652, FSE and Lema: Andalucía moves with Europe to L.L.N.), the Ministerio de Ciencia E Innovación (BFU2017-83588-P, PID2021-123083NB-I00 financed by MCIN/AEI/10.13039/501100011033 and by FEDER, UE to B.R.G,), The DiabetesCERO Foundation (B.R.G.), the Spanish Diabetes Society and the Juvenile Diabetes Research Foundation (17-2013-372, 2-SRA-2019-837-S-B, 3-SRA-2023-1307-S-B to B.R.G.). P.A.S.F. is contracted through an action funded by the EU-NextGenerationEU, and the Recovery, Transformation and Resilience Plan (SE/INV/0008/2022) granted to the Consejería de Empleo, Formación y Trabajo Autónomo of the Junta de Andalucía in the 2022 call of the Investigo Program, Mechanism for Recovery and Resilience.

## AUTHOR CONTRIBUTIONS

N.C.V., R.G-P., A.M.M., M.A.G.S., M.A.M.B., M.V.P., and B.R.G. were involved in research design. N.C.V., S.R.F., D.S-L., L.L.N., B.F.S., P.I.L, J.F.M., A.D., E.M.V., R.A.L., C.C.L., N.v.O., C.M.R., L.H., R.L.F.S., A.C.O.V., A.M.M., A.I.A., A.C.C. L.A.F. and F.M. contributed to conducting experiments. N.C.V., S.R.F., L.L.N., P.I.L, J.F.M., A.D., C.C.L., M.R.R, L.P., M.V.P., R.G-P, and B.R.G contributed to data analysis. R.N., L.P., and M.A.D. supplied human samples and reagents. N.C.V., and B.R.G. wrote the manuscript. All authors commented on the manuscript. B.R.G. is the guarantor of this work and, as such, has full access to all the data in the study and takes responsibility for the integrity of the data and the accuracy of the data analysis.

## COMTETING INTERESTS

Two patents (WO 2011 144725 A2 and WO 2016 156531 A1) related to BL001 have been published of which B.R.G and N.C.V. are inventors. M.V.-P. holds a patent that relates to immunotherapy for T1D and is co-founder and SEO of Ahead Therapeutics S.L., which aims at the clinical translation of immunotherapies for the treatment of autoimmune diseases. The other authors declare no competing interests related to the current study.

## DATA AND MATERIAL AVAILABILITY

All resources, reagents and raw data reported in this paper will be shared by the lead contact, Benoit R. Gauthier.The mass spectrometry proteomics data have been deposited to the ProteomeXchange Consortium via the PRIDE (*104*) partner repository with the dataset identifier PXD045222. The RNAseq data have been deposited to the NCBI BioProject and can be retrieved with the project ID number PRJNA1017470.

## Notes

### Competing Interest Statement

The authors have declared no competing interest.

### Summary of Updates

This revised version of the manuscript contains a proteomic analysis of dendritic cells as well as a more detailed analysis of BL001 impact on T-cell phenotype.

