## Supplemental data for "LRH-1/NR5A2 Activation Rewires Immunometabolism Blunting Inflammatory Immune Cell Progression in Individuals with Type 1 Diabetes and Enhances Human Islet Function in Mice"

[1]Andalusian Center of Molecular Biology and Regenerative Medicine-CABIMER, Junta de Andalucía-University of Pablo de Olavide-University of Seville-CSIC, Seville, Spain
[2]Immunology Department, Germans Trias i Pujol Research Institute, Autonomous University of Barcelona, Badalona, Spain
[3]Centro de Investigación Biomédica en Red de Diabetes y Enfermedades Metabólicas Asociadas (CIBERDEM), Madrid, Spain
[4]Cell and Chemical Biology, Leiden University Medical Centre, Einthovenweg 20, 2333ZC Leiden, The Netherlands
[5]Pompeu Fabra University, Barcelona, Spain
[6]Department of Endocrinology and Nutrition. University Hospital Virgen Macarena, Sevilla, Spain
[7]Diabetes Research Institute, IRCCS Ospedale San Raffaele, Milan, Italy
[8]Department of Endocrinology and Nutrition. University Hospital Puerta del Mar, Institute of Research and Innovation in Biomedical Sciences of Cádiz (INiBICA). University of Cádiz (UCA), Spain.
[9]Department of Cell Biology, Faculty of Biology, University of Sevilla, Spain
[10]Biobank of the Andalusian Public Health System, Node Hospital Virgen Macarena, Sevilla, Spain
[11]Lead contact

[*]These authors contributed equally to this work
[$]These senior authors contributed equally to this work.

[*]Correspondence: Benoit R. Gauthier (ORCID: 0000-0001-8146-7486)
CABIMER: Andalusian Center of Molecular Biology and Regenerative Medicine Avenida Américo Vespucio, 24 41092 Sevilla, Spain


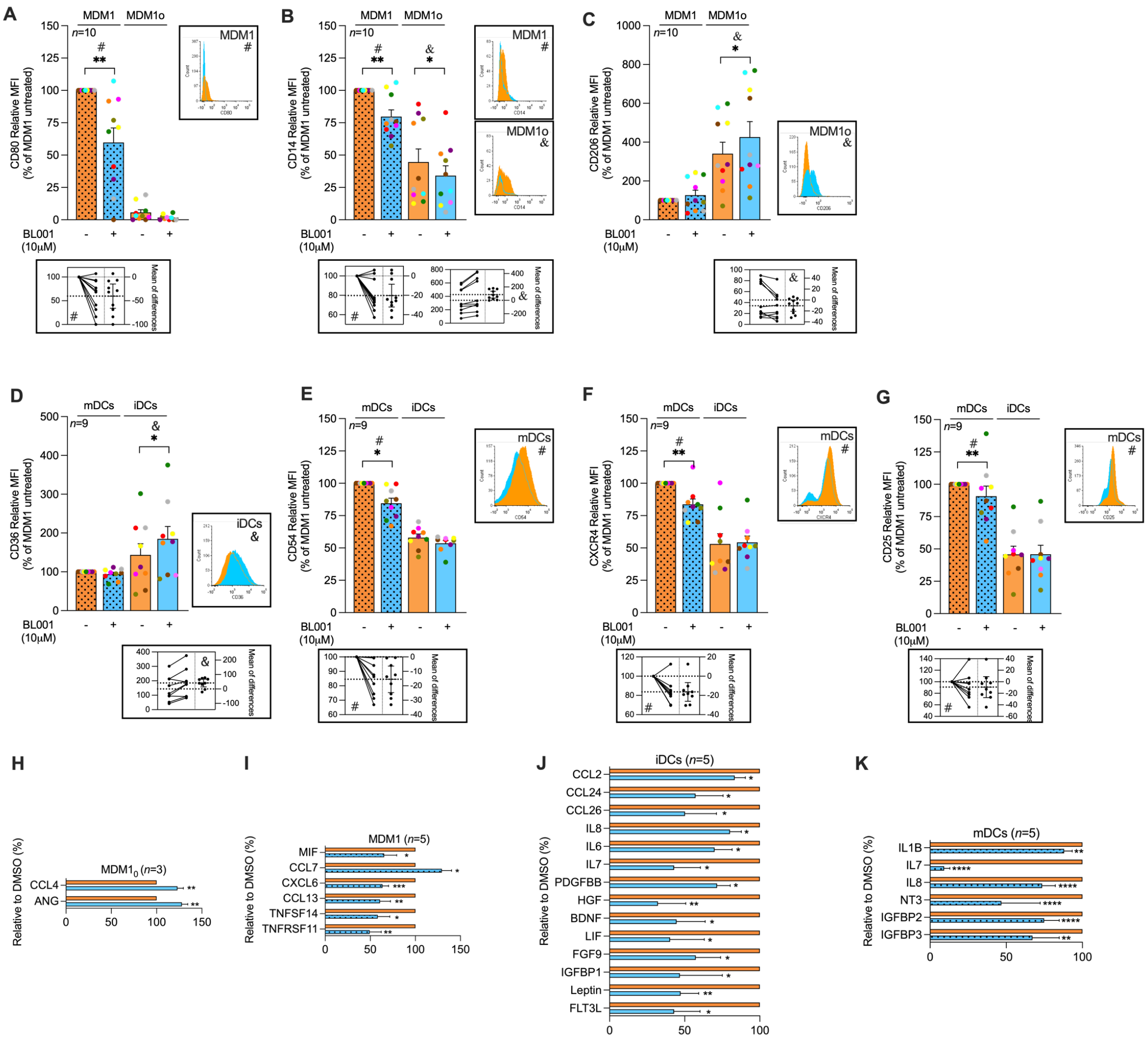


**Fig. 1. LRH-1/NR5A2 activation impedes the pro-inflammatory immune cell phenotype and cytokine secretion in T1D.** Monocytes were purified from individuals with T1D, and differentiated into either resting or pro-inflammatory macrophage (MDM1$_0$ or MDM1) and immature or mature dendritic cells (iDCs or mDCs). LRH1/NR5A1 activation was achieved by administering 10 μM BL001 every 24 hours for a total duration of 48 hours with a final dose given 30 minutes before further analysis. MDM cell surface markers (**A**) CD80, (**B**) CD14 (**C**) CD206 and DC cell surface markers (**D**) CD36, (**E**) CD54, (**F**) CXCR4, (**G**) CD25 were then assessed by flow cytometry. Measurements were normalized to the mean fluorescence intensity (MFI) of untreated MDM1 or mDC for comparison. A total of *n=10* independent individuals with T1D were analyzed for MDM markers, while *n=9* independent individuals with T1D were evaluated for mDCs markers. Each donor is colour-coded. Data are presented as means ± SEM and compared to either untreated MDM1 or mDC. Paired Student t-test * p<0.05, and ** p<0.01 as compared to untreated cells (MDM1$_0$, MDM1, iDC and mDC). Flow cytometry histograms (untreated: orange, and BL001 treated: blue) as well as estimation plots (linked to plots by either & or #) are shown only for the markers with statistically significant differences. The cytokine secretion profile was assessed for (**H**) MDM1$_0$, (**I**) MDM1, (**J**) iDCs, (**K**) mDCs**.** A total of *n=3* independent donors were analyzed for MDM1$_0$, while *n=5* independent individuals were evaluated for MDM1, iDCs, mDCs. Percent changes in cytokine secretion are presented relative to DMSO-treated counterparts for each cytokine. Only significantly altered cytokines are shown. Data are presented as percent changes compared to DMSO for each cytokine. Unpaired Student t-test * p<0.05, ** p<0.01, *** p<0.001, **** p<0.001 compared to DMSO

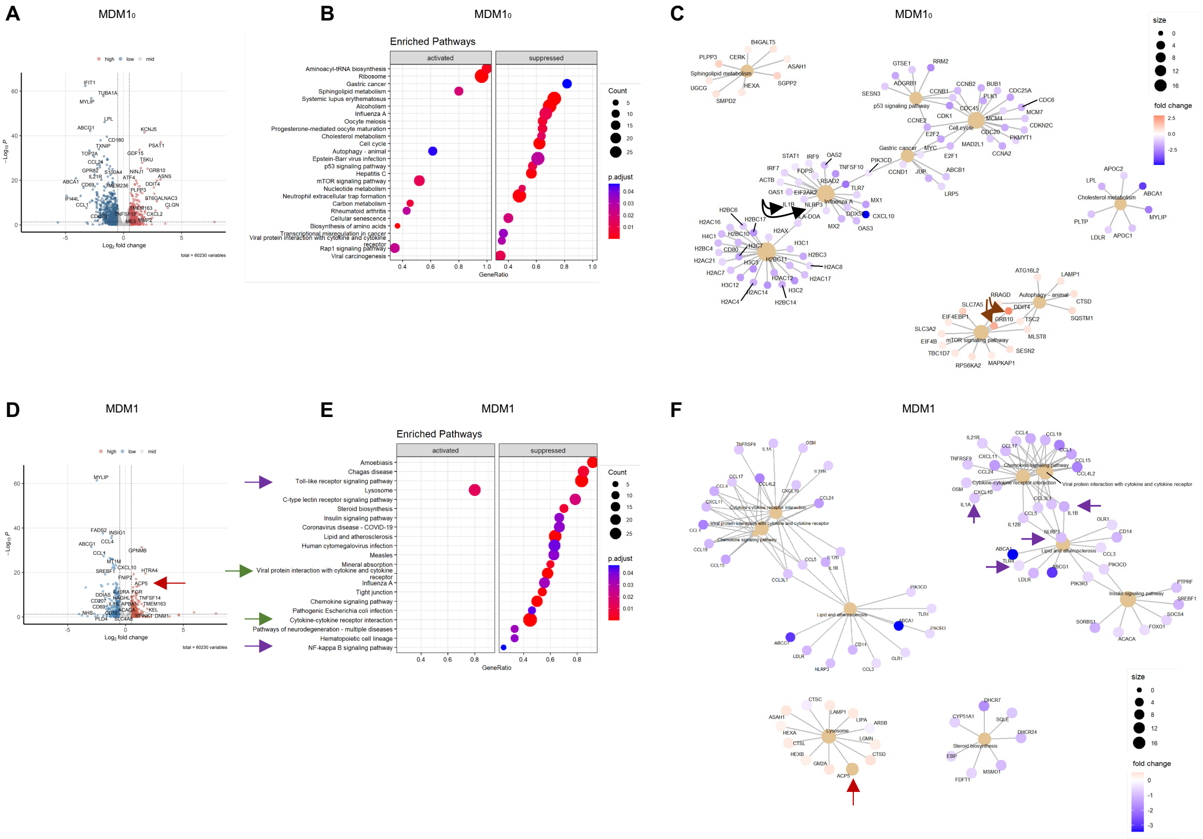


**Fig. 2. LRH-1/NR5A2 agonism mitigates the pro-inflammatory genetic program in T1D monocyte-derived macrophages (MDM).** (**A**) Volcano plot of differentially expressed genes in BL001-treated vs untreated $MDM1_0$ (*n=6* independent donors). (**B**) Dot plot of KEGG pathways enriched in BL001-treated $MDM1_0$ and (**C**) Cnetplots of selected KEGG pathways. (**D**) Volcano plot of differentially expressed genes in BL001-treated vs untreated MDM1 (*n=6*, independent donors). (**E**) Dot plot of KEGG pathways enriched in BL001-treated MDM1 and (**F**) Cnetplots of selected KEGG pathways. Arrows highlight genes of interest which are described in the results.

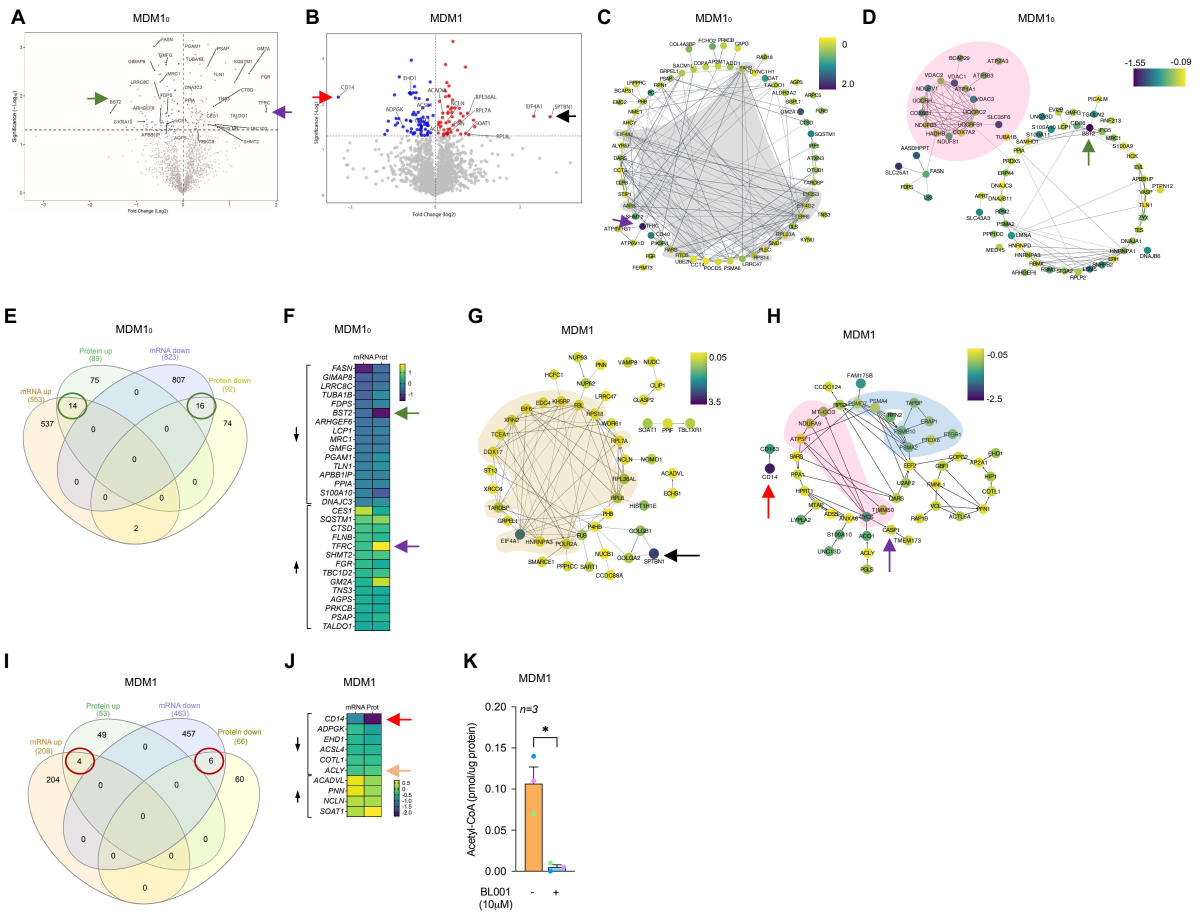


**Fig. 3. Proteomic alterations induced by BL001 in T1D monocyte-derived macrophages.** Volcano plot displaying the most significantly differentially expressed proteins in (**A**) BL001-treated versus untreated $MDM1_0$, derived from *n=3* independent donors and (**B**) BL001-treated versus untreated MDM1, from *n=3* independent donors. Arrows point to genes of interest which are described in the results section. Cytoscape circular layout of significantly (**C**) up-regulated proteins and (**D**) down-regulated proteins in BL001-treated $MDM1_0$ compared to untreated controls. RNA-associated proteins are highlighted within the grey-shaded area while mitochondrial proteins are emphasized by the pink-shaded area. (**E**) InteractiVenn diagram of differentially expressed transcripts/proteins that are significantly altered in either the RNAseq or proteomic analysis of BL001-treated versus untreated $MDM1_0$. (**F**) Heatmap of differentially expressed transcripts/proteins common to both the RNAseq and proteomic analysis in BL001-treated $MDM1_0$ versus untreated $MDM1_0$ marked by green circles in (**E**). Cytoscape circular layout of significantly (**G**) up-regulated proteins and (**H**) down-regulated proteins in BL001-treated MDM1 as compared to untreated MDM1. Proteins involved in transcriptional/translational processes are within the French beige-shaded area while mitochondrial proteins are highlighted in the pink shaded area and proteasome-associated proteins are in the blue-shaded area. (**I**) InteractiVenn diagram of differentially expressed transcripts/proteins that are significantly altered in either the RNAseq or proteomic analysis of BL001-treated versus untreated MDM1. (**J**) Heatmap of differentially expressed transcripts/proteins common to both RNAseq and proteomic analysis of BL001-treated MDM1 versus untreated $MDM1_0$(red circles in I). Arrows point to genes of interest which are described in the results section. (**K**) Bar graph representing acetyl CoA levels in MDM1 treated or not with BL001 for *n=3* independent donors colour-coded and each performed in triplicate. Data are presented as means ± SEM. Student t-test * $p<0.05$ as compared to untreated.

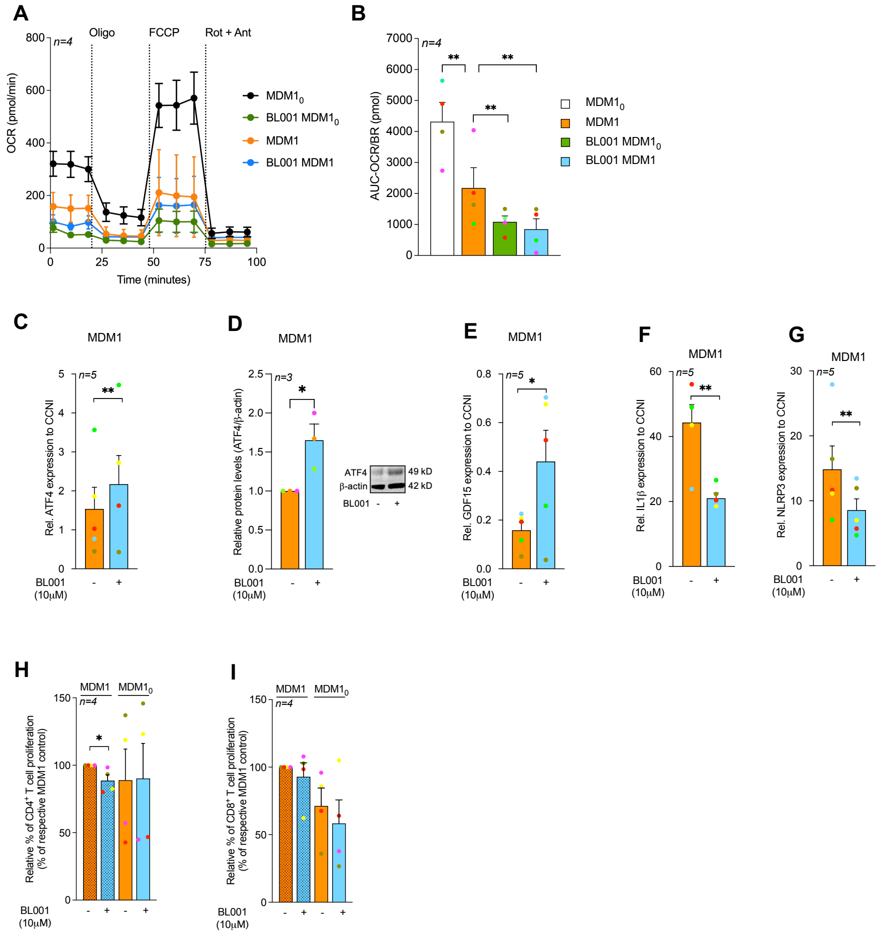

**Fig. 4. LRH-1/NR5A2 activation stimulates mitohormesis to enforce LPS-tolerance in T1D monocyte-derived macrophages.** (**A**) Mitochondrial stress test performed on T1D MDM$1_0$ and MDM1 (namely LPS/IFNg-treated MDM$1_0$) treated with or without BL001. *n=4* independent donors. (**B**) Calculated basal oxygen consumption rates (OCR-BR) Abbreviations: Oligo-Oligomycin, FCCP-Carbonyl cyanide-p-trifluoromethoxyphenylhydrazone, Rot-Rotenone, Ant-Antimycin A. Paired Student t-test **p<0.01 as compared to MDM$1_0$ or MDM1. *n=4* independent colour-coded donors. (**C**) Transcript levels of the mitohormesis-associated gene ATF4, in BL001 treated or not MDM1 cells from T1D donors, normalized to the housekeeping gene CCNI (https://housekeeping.unicamp.br/). *n=5* independent colour-coded donors. (**D**) Protein expression levels of ATF4 in BL001 treated or not MDM1 cells from T1D individuals, and normalized to the housekeeping protein b-actin. *n=3* independent colour-coded donors. The figure includes a representative western blot image. Transcript levels of (**E**) GDF15, (**F**) IL-1b, and (**G**) NLRP3 in T1D MDM1 treated with or without BL001. Transcript levels were normalized to the housekeeping gene CCNI (https://housekeeping.unicamp.br/). *n=5* independent colour-coded donors. Data are presented as means ± SEM. Paired Student t-test * p<0.05, **p<0.01 as compared to untreated cells. Relative proliferation of autologous (**H**) $CD4^+$ and (**I**) $CD8^+$ T cells in response to co-culture with healthy or T1D MDMs treated or not with 10 µM BL001. *n=4* independent colour-coded donors. Paired Student t-test *p<0.05 as compared to untreated cells.

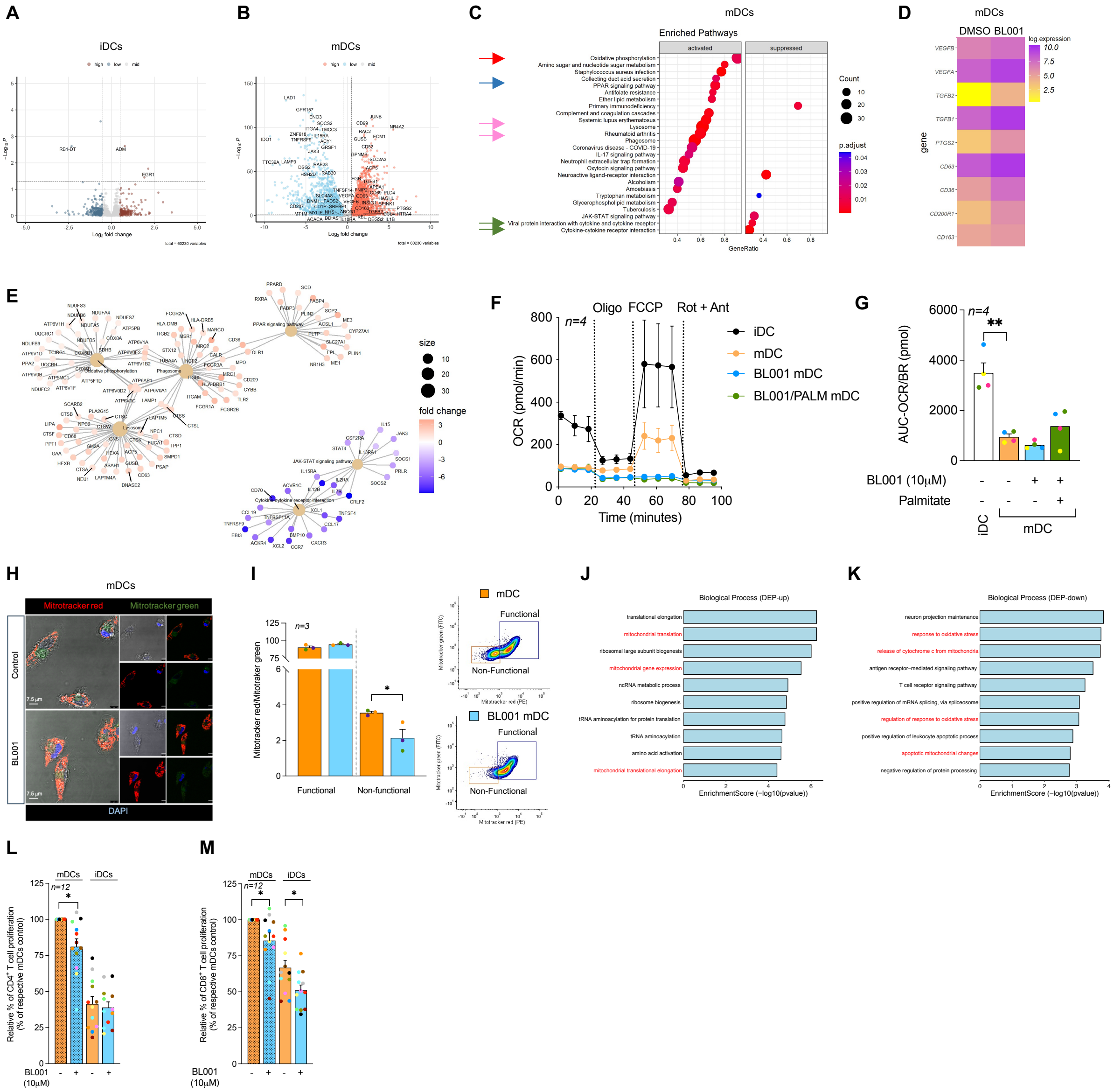

**Fig. 5. LRH-1/NR5A2 Promotes a Tolerogenic Phenotype to T1D mDC, Suppressing Autologous T-cell Proliferation.** Volcano plot of differentially expressed genes in BL001-treated vs untreated (**A**) iDCS (*n=5* independent donors) and (**B**) mDCs (*n=4* independent donors). (**C**) Dot plot of KEGG pathways enriched in BL001-treated mDCs. Arrows point to pathways of interest which are described in the results section. (**D**) Heatmap of selected genes associated with a tolerogenic DC phenotype. Gene expression is presented as the average log(NC). (**E**) Cnetplots of selected KEGG pathways modulated by BL001 in mDCs. (**F**) Mitochondrial stress test and (**G**) calculated basal oxygen consumption rates (OCR) in T1D iDC and mDC treated with or without BL001 and with palmitate. Abbreviations: Oligo-Oligomycin, FCCP-Carbonyl cyanide-p-trifluoromethoxyphenylhydrazone, Rot-Rotenone, Ant-Antimycin A. *n=4* independent donors (colour matched). Paired Student t-test ** p<0.01 as compared to iDCs. Each donor is colour-coded. (**H**) Representative confocal immunofluorescence images of cells labeled with MitoTracker green and MitoTracker red for mitochondria, with nuclei counterstained using DAPI. (I) Bar chart quantification for mitochondrial functionality, as determined by flow cytometry based on MitoTracker green and MitoTracker red. Representative flow cytometry plots are provided for illustration. *n=3* independent colour-code donors. Paired Student t-test * p<0.05 as compared to untreated mDC. Bar plot ranking of the top ten GO biological process terms associated with (**J**) up-regulated and (**K**) down-regulated proteins in BL001-treated mDCs. Relative autologous proliferation of (**L**) $CD4^+$ and (**M**) $CD8^+$ T-cells in the presence of T1D DCs treated with or without 10 μM BL001. Data are presented as means ± SEM. Paired Student t-test * p<0.05 as compared to untreated cells.

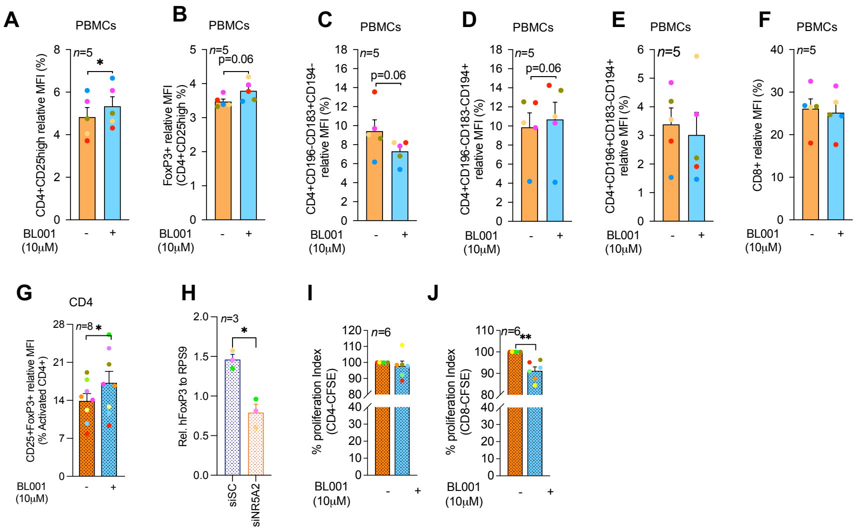

**FIG. 6. LRH-1/NR5A2 agonism promotes the expansion of a CD4⁺/CD25⁺/FoxP3⁺ cell subpopulation, leading to reduced CD8⁺ T-cell proliferation in T1D individuals.**
PBMCs were purified from individuals with T1D and exposed to 10 μM BL001 every 24 hours for a total duration of 48 hours with a final dose given 30 minutes before further analysis. Cells were then analysed by flow cytometry using a combination of cell surface markers and gated for subpopulations of (**A**) $CD4^+CD25^+$, (**B**) Tregs; $CD4^+CD25^+FoxP3^+$ (**C**) Th1; $CD4^+CD196^-CD183^+CD194^-$, (**D**) Th2; $CD4^+CD196^-CD183^-CD194^+$, (**E**) Th17/22; $CD4^+CD196^+CD183^-CD194^+$ and (**E**) $CD8^+$. *n=5* independent individuals with T1D. (**G**) $CD4^+$ cells were isolated from PBMCs and treated with BL001 as described above. Cells were then analysed by flow cytometry for the cell surface markers $CD25^+FoxP3^+$ and results plotted as the percentage of $CD25^+FoxP3^+$ cells within the $CD4^+$ subpopulation. *n=8* independent individuals with T1D. All cytometry measurements were normalized to the mean fluorescence intensity (MFI). (**H**) Relative FoxP3 transcript levels in either siScrambled (siSc) or siNR5A2-treated PBMCs. Data were normalized to the housekeeping gene RSP9. *n=3* T1D independent donors. $CD14^-$ PBMCs were labelled with CFSE and stimulated/expanded using a polymeric nanomatrix structure composed of CD3 and CD28 for 24 hours before the addition of $CD4^+$ T-cells (at a 1:2 ratio, respectively, from the same donor). Proliferation of (**I**) $CD4^+/CFSE^+$ and (**J**) $CD8^+/CFSE^+$ subpopulations was assessed by flow cytometry 4 days post co-culturing. *n=6* T1D independent donors. Each donor is colour-coded. Data are presented as means ± SEM. Paired Student t-test * p<0.05, and ** p<0.01 as compared to untreated cells.

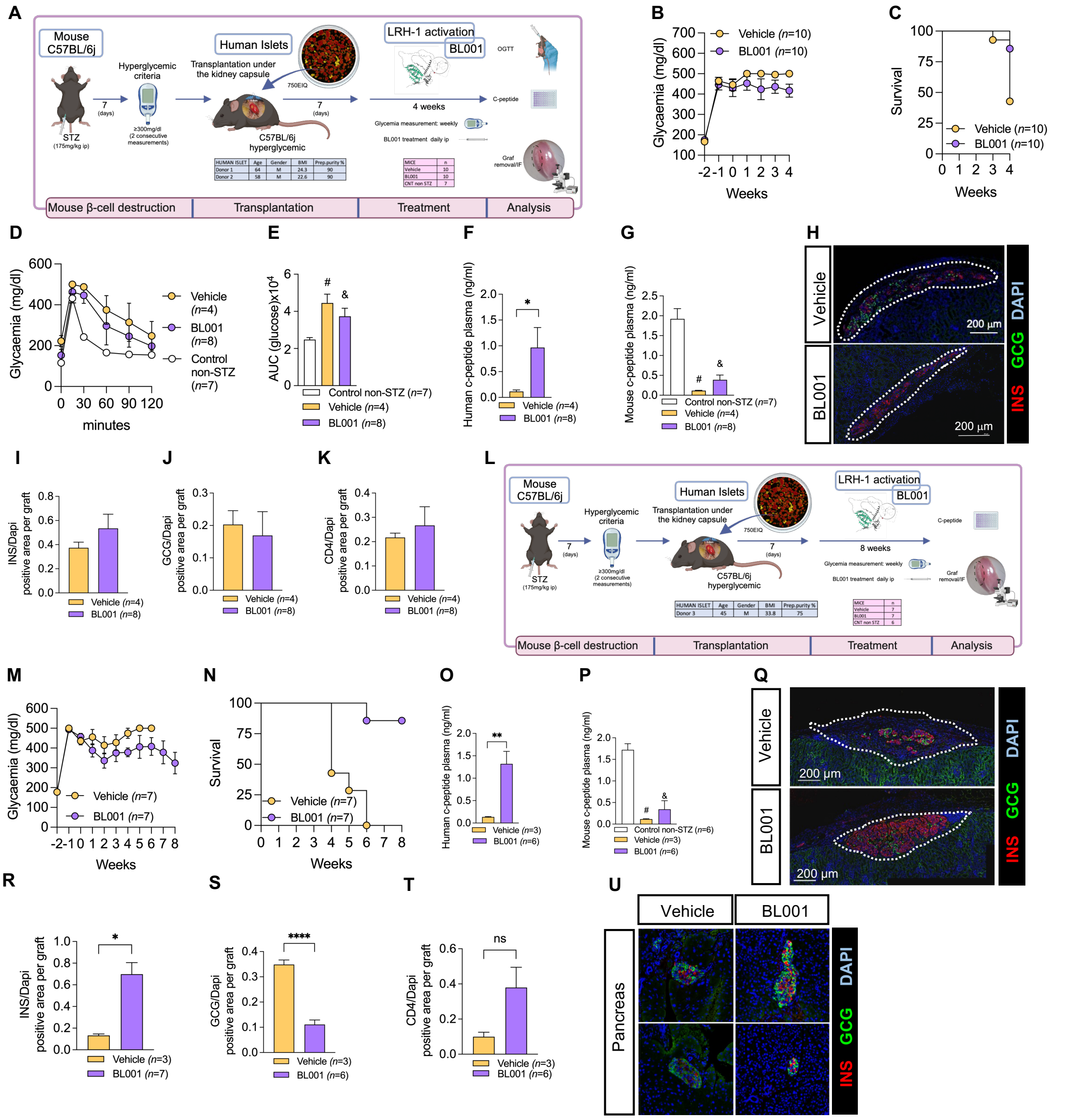

**Fig 7. BL001 improves human islet graft survival and function in STZ-treated immunocompetent C57BL/6j mice.** (**A**) Experimental design of the 4-week BL001 treatment post-xenotransplantation experiment. (**B**) Weekly measurement of non-fasting blood glucose and (**C**) Kaplan-Meier survival curve. (**D**) OGTT performed at 4 weeks post-BL001/Vehicle treatment. Mice were fasted for 6 hours before the OGTT. (**E**) Area under the curve (AUC) corresponding to the OGTT. Student t-test, # *p=0003* and & *p=0.0037* as compared to control non-STZ mice. (**F**) Human C-peptide plasma levels at 4 weeks post-BL001/Vehicle treatment. Data are presented as means ± SEM. *p<0.05 student t-test. (**G**) Mouse C-peptide plasma levels at 4 weeks post-BL001/Vehicle treatment. Data are presented as means ± SEM. Student t-test, # *p=002* and & *p=0.0003* as compared to control non-STZ mice. (**H**) Representative immunofluorescence images of kidney sections from mice euthanized at 4 weeks post-BL001/Vehicle treatment, displaying staining for insulin (INS), glucagon (GCG) staining along with nuclear DAPI staining. Quantitative analysis of (**I**) insulin (INS), (**J**) glucagon (GCG), and (**K**) $CD4^+$ areas, normalized to the DAPI-positive area per graft, at 4 weeks post BL001 treatment. (**L**) Experimental design of the 8-week BL001 treatment post-xenotransplantation experiment. (**M**) Weekly measurement of non-fasting glycemia and (**N**) Kaplan Meier survival curve. (**O**) Human C-peptide plasma levels at 6-week vehicle-treated mice and 8-week BL001-treated mice. Data are presented as means ± SEM. **p<0.01 student t-test. (**P**) Mouse C-peptide plasma levels at 6-week vehicle-treated mice and 8-week BL001-treated mice. Data are presented as means ± SEM. # *p=0001* and & *p=0.*0002 Student t-test as compared to control non-STZ mice. (**Q**) Representative immunofluorescence images of kidney sections from mice at 6 weeks post-vehicle treatment and at 8 weeks post-BL001 treatment, displaying staining for insulin (INS), glucagon (GCG) staining along with nuclear DAPI staining. Quantitative analysis of (R) insulin (INS), (**S**) glucagon (GCG), and (**T**) $CD4^+$ areas, normalized to the DAPI-positive area per graft, at 4 weeks post BL001 treatment. Data are presented as means ± SEM. *p<0.05 and ****p<0.0001 student t-test. ns, non-significant. (**U**) Representative immunofluorescence images of pancreas sections from mice at 6 weeks post-vehicle treatment and at 8 weeks post-BL001 treatment. displaying staining for insulin (INS), glucagon (GCG) staining along with nuclear DAPI staining.

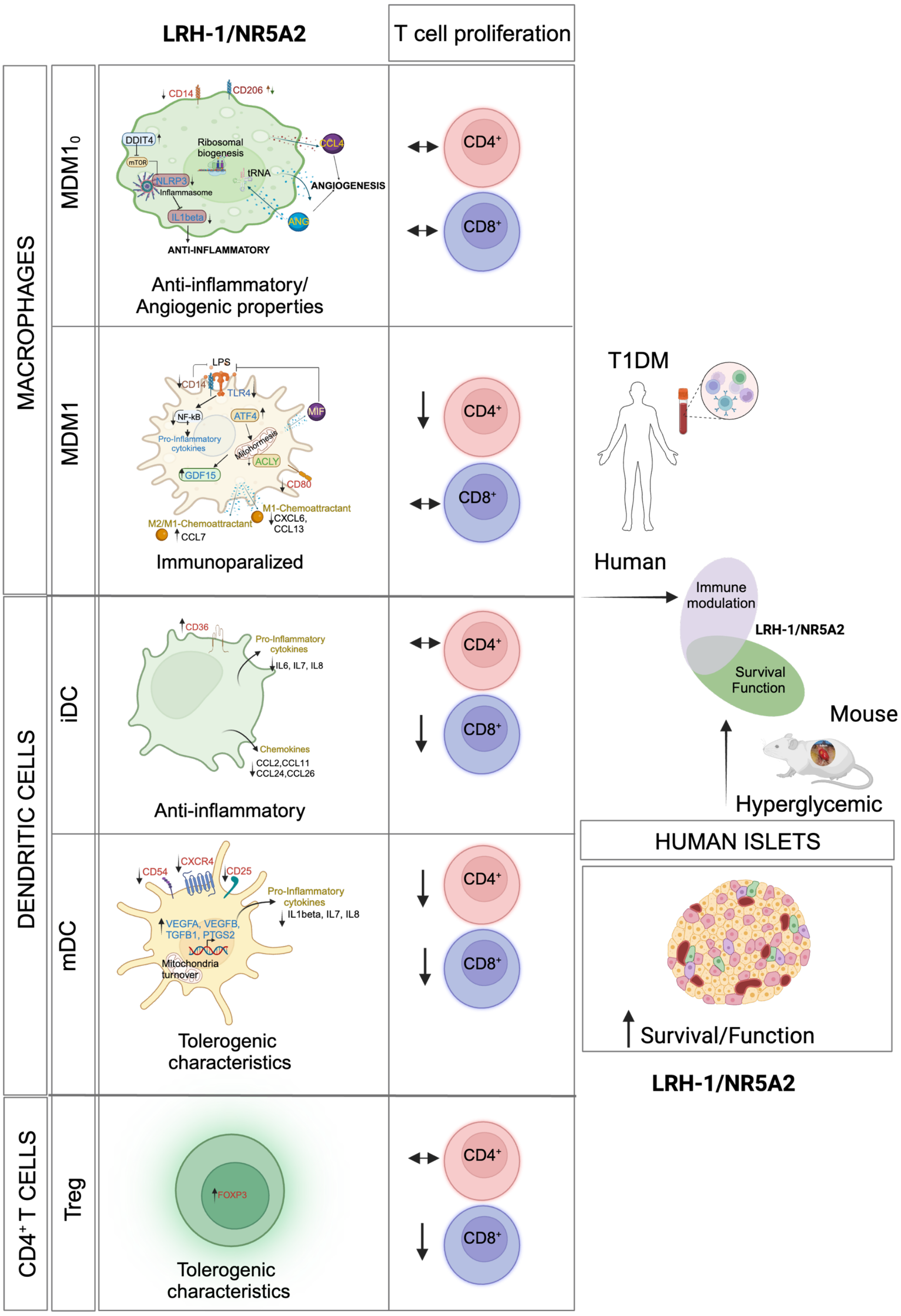


**Fig. 8. Cellular and molecular mechanism of action mediated by the pharmacological effect of LRH-1/NR5A2 in human cells.** Proposed model of genetic and immune cell-tailored reprogramming induced by LRH-1/NR5A2 activation and its impact on T-effector cell proliferation. Improvement of human islet engraftment influenced by LRH-1/NR5A2 activation.

Table 1. Demographic and clinical characteristics of donors with T1D

| Characteristics | |
|---|---|
| N | 68 |
| Gender (F/M) | 32F / 36M |
| Age (years) | 37 ± 10 |
| BMI (kg/m$^2$) | 24 ± 4 |
| Age at diagnosis (years) | 20 ± 12 |
| Progression (years) | 16 ± 11 |
| HbA1c (%) | 7 ± 0,9 |
| Insulin dose (IU/kg/day) | 0,53 ± 0,17 |

**Table 2:** Key resources

| REAGENT or RESOURCE | SOURCE | IDENTIFIER |
|---|---|---|
| Antibodies | | |
| CCR7 PECy7 | BD Biosciences | Cat#557648 |
| Annexin V PE | Immunotools | Cat#31490014 7 |
| CD3 PE | Immunotools | Cat#21620034 |
| CD4 APC | Immunotools | Cat#21278046 |
| CD8 FITC | Immunotools | Cat#21810083 |
| CD11c APC | Immunotools | Cat#21487116 |
| CD14 PE | Immunotools | Cat#21620144 |
| CD25 PE | Immunotools | Cat#21810254 |
| CD40 APC | Immunotools | Cat#21270406 |
| CD86 FITC | Immunotools | Cat#21480863 |
| HLA class I FITC | Immunotools | Cat#21159033 |
| HLA class II FITC | Immunotools | Cat#21388993 |
| αvβ5 integrin PE | BioLegend | Cat#920007 |
| CD36 APCCy7 | BioLegend | Cat#336213 |
| CD54 PECy7 | BioLegend | Cat#353115 |
| CCR2 APC | BioLegend | Cat#357207 |
| CXCR4 APCCy7 | BioLegend | Cat#306528 |
| TIM4 APC | BioLegend | Cat#354008 |
| DC-SIGN APC | BioLegend | Cat#330108 |
| PD-L1 PECy7 | BioLegend | Cat#329717 |
| CTV Cell Proliferation Kit | ThermoFisher Scientific | Cat#C34557 |
| CSFE | ThermoFisher Scientific | Cat#C34554 |
| Zombie Violet 421 | BioLegend | Cat#423114 |
| CD4 | Miltenyi Biotec | Cat#130-122-994 |
| CD8 | eBioscience | Cat#45-0088-41 |
| CD14 | BioLegend | Cat#325604 |
| CD25 | Miltenyi Biotec | Cat#130-122-994 |
| CD80 | Miltenyi Biotec | Cat#130-123-253 |
| CD86 | BioLegend | Cat#305412 |
| CD163 | Miltenyi Biotec | Cat#130-112-126 |
| CD183 | Miltenyi Biotec | Cat#130-120-452 |
| CD194 | Miltenyi Biotec | Cat#130-117-376 |
| CD196 | Miltenyi Biotec | Cat#130-127-189 |
| CD206 | Miltenyi Biotec | Cat#130-095-131 |
| CD200R | Miltenyi Biotec | Cat#130-111-290 |
| CD209 | Miltenyi Biotec | Cat#130-092-873 |
| FoxP3 | Miltenyi Biotec | Cat#130-122-994 |
| ACTIN | Sigma-Aldrich | Cat#A5441 |
| ATF4 | Cell Signaling | Cat#11815 |
| INSULIN | Sigma-Aldrich | Cat#I2018 |
| GLUCAGON | Cell Signaling | Cat#2760 |
| Biological samples | | |
| Human islets | ADI IsletCore Laboratory, Edmonton, CA and Vita-Salute San Raffaele University, Milan (IT) | N/A |

| | | |
|---|---|---|
| Blood samples | Hospital Germans trias I Pujol, Badalona and the University Hospital Virgen Macarena, Sevilla (ES) | N/A |
| Chemicals, peptides, and recombinant proteins | | |
| BL001 | In house | In house |
| rhIL-4 | Prospec | Cat#CYT-211 |
| rhGM-CSF | Prospec | Cat#CYT-221 |
| Human insulin | Sigma-Aldrich | Cat#I3536 |
| TNFα | Immunotools | Cat#11343015 |
| IL-1β | Immunotools | Cat#11340013 |
| PGE2 | Cayman Chemical | Cat#14010 |
| Streptozotocin | Sigma-Aldrich/Merck | Cat#S0130-1G |
| Penicillin | Sigma-Aldrich/Merck | Cat#P0781 |
| Streptomycin | Sigma-Aldrich/Merck | Cat#P0781 |
| INFγ | Immunotools | Cat#11343536 |
| LPS | Sigma-Aldrich/Merck | Cat# L6529 |
| Critical commercial assays | | |
| Cell Mito Stress Test Kit | Agilent | Cat#103015-100 |
| Treg Phenotyping Kit, anti-human, REAfinity | Miltenyi Biotec | Cat#130-122-994 |
| Human Cytokine Array C5 | RayBiotech | Cat#AAH-CYT-5-2 |
| Human C-peptide ELISA kit | Mercodia | Cat#10-1136-01 |
| Mouse C-peptide ELISA kit | Mercodia | Cat#10-1247-01 |
| Acetyl Coenzyme A assay kit | Sigma-Aldrich | Car#MAK039-1KT |
| CD4+ T cell isolation kit | Stemcell Technology | Cat# 17952 |
| Deposited data | | |
| RNAseq dataset | This paper | |
| Proteomic dataset | This paper | |
| Experimental models: Organisms/strains | | |
| C57BL/6J | Janvier Labs | strain code 027/SC-C57J-F |
| Oligonucleotides | | |
| Primer: ATF4 Fw: TCAAACCTCATGGGTTCTCC Rv: GTGTCATCCAACGTGGTCAG | | |
| Primer: CCNI Fw: GCACAGATGGATAGCTCC Rv: CTTTGTCACAGGTCACCA | | |
| Primer: FoxP3 Fw: GGCACAATGTCTCCTCCAGAGA Rv: CAGATGAAGCCTTGGTCAGTGC | | |
| Primer: GDF15 Fw: ACCTGCACCTGCGTATCTCT Rv:CGGACGAAGATTCTGCCAG | | |
| Primer: IL1B Fw: AGCTACGAATCTCCGACCAC Rv: CGTTATCCCATGTGTCGAAGAA | | |
| Primer: NLRP3 Fw: CGTGAGTCCCATTAAGATGGAGT Rv: CCCGACAGTGGATATAGAACAGA | | |
| Primer: RSP9 Fw: AAGGCCGCCCGGGAACTGCTGAC Rv: ACCACCTGCTTGCGGACCCTGATA | | |
| Software and algorithms | | |
| FlowJo | Tree Star Inc | www.flowjo.com |

| | | |
|---|---|---|
| FCS Express | De Novo Software | https://denovosoftware.com/ |
| Fiji | Imagej | https://imagej.net/software/fiji/downloads |
| Prism | GraphPad | https://www.graphpad.com/ |
| Adobe Photoshop | Adobe | https://www.adobe.com/es/ |
| ImageJ | Imagej | https://imagej.nih.gov/ij/ |
| Cytoscape | Cytoscape | https://cytoscape.org/index.html |
| SRplot | SRplot | https://www.bioinformatics.com.cn/srplot |
| Other | | |
| BD Vacutainer Sodium Heparin tubes | BD | Cat#366667 |
| Ficoll Paque PLUS | Cytiva | Cat#17144003 |
| Histopaque 1077 | Sigma-Aldrich | Cat#10771 |
| EasySep Human CD14 Positive Selection kit II | STEMCELL Technologies | Cat#17858 |
| EasySep Human CD4+ T cell isolation kit | STEMCELL Technologies | Cat#17952 |
| Accutase | ThermoFisher Scientific | Cat#00-4555-56 |
| 7-AAD | BD Biosciences | Cat#559925 |
| MitoTracker Green | ThermoFisher Scientific | Cat#M7514 |
| MitoTracker Red | ThermoFisher Scientific | Cat#M7512 |
| T-Cell TransAct human | Miltenyi Biotec | Cat#130-111-160 |
| On-target plus Human NR5A2 (2495) siRNA-smart pool | Dharmacon | L-003430-00-0005 |
| On-target plus non-targeting control pool | Dharmacon | D-001810-10-20 |

**Supplementary Materials**

### LRH-1/NR5A2 Activation Rewires Immunometabolism Blunting Inflammatory Immune Cell Progression in Individuals with Type 1 Diabetes and Enhances Human Islet Function in Mice

Cobo-Vuilleumier *et al*.

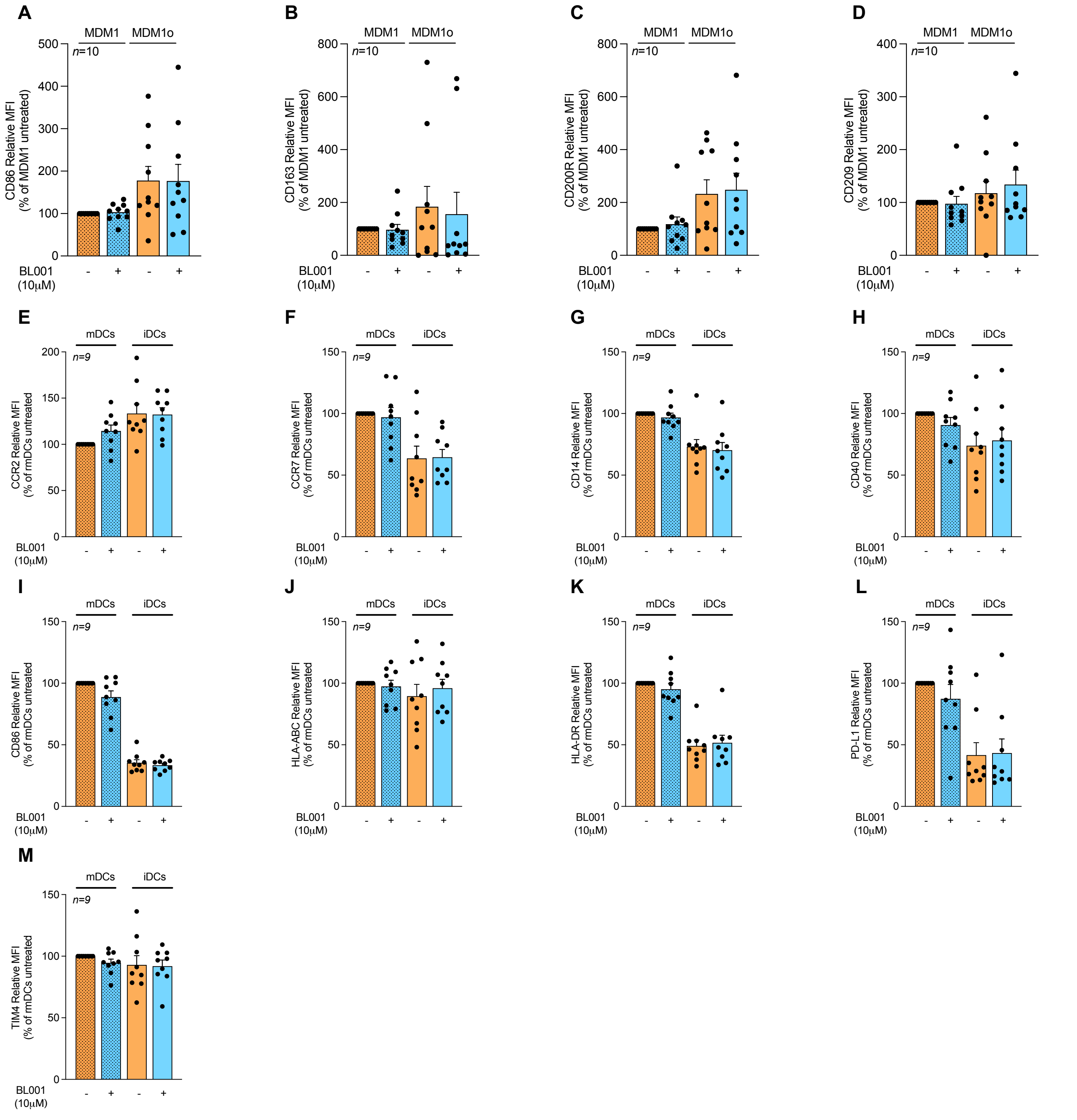


**Fig. S1: Cell surface markers of monocyte-derived macrophages (MDM) and dendritic cells (DCs) not altered by BL001 treatment.** Monocytes were purified from healthy and T1DM individuals and derived into either resting or pro-inflammatory macrophage and dendritic cells (MDM1$_0$ or MDM1 and iDCs or mDCs) and treated with 10 uM BL001 for 48 hours and then 30 minutes prior to analysis. MDM cell surface markers (**A**) CD86, (**B**) CD163, (**C**) CD200R (D) CD209 and DC surface markers (**E**) CCR2, (**F**) CCR7, (**G**) CD14, (**H**) CD40, (**I**) CDCD86, (**J**) HLA-ABC, (**K**) HLA-DR, (**L**) PD-L1 and (**M**) TIM4 were then assessed by flow cytometry.

| KEGG TERM | Adjusted p value | GENES |
|---|---|---|
| Coronavirus disease - COVID-19 | 5.824370e-10 | MX1,MX2,ISG15,OAS3,OAS2,OAS1,TLR7,STING1,EIF2AK2,FCGR2A,VWF,IRF9,STAT1,TLR8,PRKCA,FOS,IKBKE,NRP1,RPS15A,RPL3,RPL37A,RPL32,RPL9,RPS16,RPS13,RPS6,RPS3A,RPL30,RPL13A,RPL28,PRKCB,RPL23,RPS23,RPL15,RPS18,RPL13,RPL12,RPL18A,IRAK1,RPL17,RPLP0,MAPK13,CXCL8 |
| Neutrophil extracellular trap formation | 0.0000025864 | H2AC14,H3C3,H2BC17,H3C2,H3C12,H3C7,H2BC3,H2AC12,H2BC14,H2BC10,H2BC11,H2AC17,H2AC7,H2AC16,H2BC6,TLR7,H2AX,H2AC21,FCGR3A,FCGR2A,VWF,HDAC9,TLR8,PRKCA,HDAC7,FPR3,H2BC12,ACTG1,MACROH2A1,PRKCB,ITGAL,SLC25A6,MAPK13,FPR1 |
| Cell cycle | 0.0000044095 | CDK1,CDC25A,E2F2,CCNA2,CDC45,CDC20,CCNE2,PLK1,CDC6,E2F1,CCNB2,CDKN2C,BUB1,MCM7,CCNB1,MAD2L1,PKMYT1,CCND1,MYC,MCM3,PCNA,SKP2,MCM6,TFDP1,RB1,YWHAZ |
| Systemic lupus erythematosus | 0.0002105552 | H2AC14,H3C3,H2BC17,H3C2,H3C12,H3C7,H2BC3,H2AC12,H2BC14,H2BC10,H2BC11,H2AC17,H2AC7,H2AC16,H2BC6,CD80,HLADOA,H2AX,H2AC21,FCGR3A,FCGR2A,TRIM21,H2BC12,MACROH2A1 |
| Ribosome | 0.0076197549 | RPS15A,RPL3,RPL37A,RPL32,RPL9,RPS16,RPS13,RPS6,RPS3A,RPL30,RPL13A,RPL28,RPL23,RPS23,RPL15,RPS18,RPL13,RPL12,RPL18A,RPL17,RPLP0 |
| Alcoholism | 0.0086872840 | H2AC14,H3C3,H2BC17,H3C2,H3C12,MAOB,H3C7,H2BC3,H2AC12,H2BC14,H2BC10,H2BC11,H2AC17,H2AC7,H2AC16,GNG2,H2BC6,H2AX,H2AC21,PKIA,HDAC9,HDAC7,H2BC12,PRKACB,MACROH2A1,SLC29A1 |
| Epstein-Barr virus infection | 0.0124044321 | E2F2,CCNA2,CCNE2,ISG15,E2F1,OAS3,OAS2,OAS1,HLADOA,IRF7,MAP2K6,EIF2AK2,CCND1,MYC,IRF9,ENTPD1,STAT1,CALR,RBPJ,SKP2,RB1,IKBKE,ITGAL,IRAK1,NCOR2,TNFAIP3,MAPK13 |
| Viral carcinogenesis | 0.0162088000 | H2BC17,CDK1,H2BC3,CCNA2,CDC20,CCNE2,H2BC14,H2BC10,H2BC11,EGR3,H2BC6,IRF7,EIF2AK2,CCND1,HDAC9,IRF9,HDAC7,RBPJ,SKP2,H2BC12,PRKACB,RB1,PKM,YWHAZ,STAT5B,PXN,ATP6V0D2 |
| Hepatitis B | 0.0283354491 | E2F2,CCNA2,CCNE2,E2F1,EGR3,BIRC5,NFATC2,IRF7,MAP2K6,MYC,STAT1,PRKCA,PCNA,FOS,RB1,IKBKE,YWHAZ,STAT5B,PRKCB,IRAK1,MAPK13,HSPG2,CXCL8 |

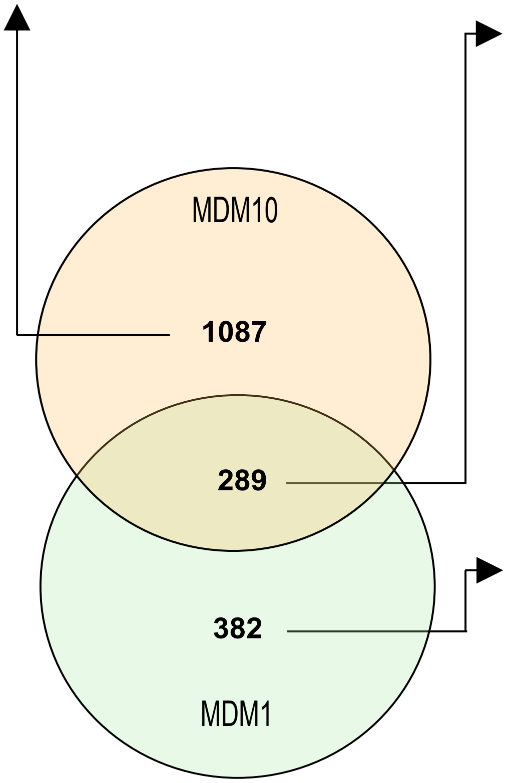


| KEGG TERM | Adjusted p value | GENES |
|---|---|---|
| Hematopoietic cell lineage | 0.0002254542 | CD1E,CD1C,CD1B,ITGA4,IL1B,CSF1R,ANPEP,CD37,TFRC,FCER2,ITGB3 |
| PPAR signaling pathway | 0.0018393227 | SCD,FADS2,HMGCS1,ACSL4,OLR1,ACSL3,PPARD,CYP27A1,PLIN2 |
| Fatty acid metabolism | 0.0018705126 | SCD,FADS2,FADS1,ACSL4,ACSL3,ELOVL5,ACACA,ACADVL |
| Steroid biosynthesis | 0.0031372665 | DHCR24,SQLE,EBP,MSMO1,CYP51A1 |
| Terpenoid backbone biosynthesis | 0.0053098308 | HMGCS1,HMGCR,IDI1,MVK,MVD |
| Ferroptosis | 0.0126550104 | MAP1LC3A,ACSL4,LPCAT3,ACSL3,SLC3A2,TFRC |

| KEGG TERM | Adjusted p value | GENES |
|---|---|---|
| NF-kappa B signaling pathway | 0.0000023458 | PLAU,CCL4L2,CCL4,CD14,CCL19,TRAF1,TLR4,MAP3K14,PTGS2,CD40,BIRC3,CFLAR,CYLD,NFKB1,RELB |
| TNF signaling pathway | 0.0000066557 | CSF1,MMP9,CCL5,IL15,NOD2,PIK3R3,TRAF1,MAP3K14,PTGS2,RPS6KA5,CASP7,BIRC3,MAP2K1,CFLAR,NFKB1 |
| Toll-like receptor signaling pathway | 0.0000133028 | CCL4L2,CCL4,CD14,CXCL11,IL12B,CCL5,PIK3R3,TLR4,CCL3,CXCL9,TLR2,CD40,MAP2K1,NFKB1 |
| Cytokine-cytokine receptor interaction | 0.0001377890 | CSF1,CCL4L2,CCL15,CCL4,CCL19,CXCL11,IL1A,TNFRSF9,IL12B,CCL5,IL2RA,ACVRL1,IL15,CCL3,CXCL9,EBI3,ACVR2A,CXCL16,CD40,IL13RA1,TNFRSF10B,IL27RA |
| Apoptosis | 0.0017643701 | CTSL,PIK3R3,TRAF1,MAP3K14,PMAIP1,CASP7,BIRC3,TNFRSF10B,BAK1,MAP2K1,CFLAR,CTSC,NFKB1 |
| Chemokine signaling pathway | 0.0129764758 | CCL4L2,CCL15,CCL4,CCL19,CXCL11,CCL5,PIK3R3,CCL3,CXCL9,CXCL16,HCK,MAP2K1,NFKB1,ARRB1 |
| Transcriptional misregulation in cancer | 0.0154891784 | MAF,MMP9,PLAU,CD14,TRAF1,FOXO1,SPINT1,CDKN1A,CD40,NFKBIZ,BIRC3,BAK1,ELK4,NFKB1 |

**Fig. S2: Distinct and common gene sets altered by BL001 in T1DM MDM.** A Venn diagram of the DEGs from BL001-treated MDM$1_0$ and MDM1 was generated using InteractiVenn. Individual genes sets were then clustered in KEGG pathways using g:GOSt of g:Profiler.

**Table S1**: Proteins which are significantly decreased in BL001-treated MDM10 obtained from individuals with type 1 diabetes mellitus.

| Protein | -Log(p) | Log2(FC) |
|---|---|---|
| BST2 | 1,66 | -1,55 |
| SLC25A1 | 1,50 | -1,32 |
| SLC35F6 | 1,35 | -1,21 |
| VDAC1 | 2,22 | -1,13 |
| ALCAM | 1,51 | -1,11 |
| UNC93B1 | 1,88 | -1,11 |
| AASDHPPT | 1,75 | -1,09 |
| MIA3 | 1,36 | -1,04 |
| UNC13D | 2,14 | -0,96 |
| LSM5 | 2,01 | -0,92 |
| SLC43A3 | 1,69 | -0,90 |
| DNAJB6;DNAJB7 | 1,64 | -0,87 |
| VDAC3 | 1,31 | -0,86 |
| TM9SF2 | 1,38 | -0,84 |
| S100A10 | 1,47 | -0,83 |
| LMNA | 1,59 | -0,81 |
| NDUFV1 | 1,33 | -0,79 |
| S100A11 | 1,82 | -0,77 |
| TGOLN2 | 1,51 | -0,77 |
| SNRPB2 | 1,36 | -0,77 |
| TM9SF3 | 1,42 | -0,77 |
| COX6B1 | 1,31 | -0,72 |
| LSS | 1,71 | -0,67 |
| NDUFS1 | 1,46 | -0,67 |
| RBM3 | 1,36 | -0,65 |
| RPN2 | 1,47 | -0,64 |
| FASN | 3,03 | -0,62 |
| GIMAP8 | 2,53 | -0,61 |
| CD86 | 1,41 | -0,60 |
| ATP1B3 | 1,98 | -0,60 |
| BCAP29 | 1,41 | -0,57 |
| ZYX | 2,64 | -0,52 |
| ATP1A1 | 2,88 | -0,51 |
| ATP2A3 | 2,44 | -0,51 |
| PSMA2 | 1,32 | -0,51 |
| LRRC8C | 1,95 | -0,49 |
| MRC1 | 2,29 | -0,48 |
| UQCRH | 1,73 | -0,48 |
| HNRNPA1;HNRNPA1L2 | 2,46 | -0,48 |
| CORO7;CORO7-PAM16 | 2,79 | -0,47 |
| PPP1CC | 1,39 | -0,47 |
| IFI35 | 1,40 | -0,47 |
| FTH1 | 1,74 | -0,47 |
| ARHGEF6 | 1,45 | -0,47 |
| VDAC2 | 1,80 | -0,45 |
| DNAJA1 | 2,32 | -0,45 |
| UQCRC2 | 2,02 | -0,45 |

| Protein | -Log(p) | Log2(FC) |
|---|---|---|
| SIGLEC9 | 1,49 | -0,45 |
| EHBP1L1 | 1,63 | -0,44 |
| SF3A2 | 1,47 | -0,44 |
| ERP44 | 1,45 | -0,41 |
| UQCRFS1;UQCRFS1P1 | 1,89 | -0,40 |
| MAGED2 | 1,31 | -0,40 |
| CD97 | 2,69 | -0,38 |
| RPLP2 | 1,64 | -0,37 |
| HADHB | 1,58 | -0,36 |
| GMFG | 2,65 | -0,36 |
| NDUFB3 | 1,38 | -0,36 |
| GBF1 | 1,38 | -0,35 |
| EVL | 1,40 | -0,35 |
| FDPS | 1,65 | -0,34 |
| MED15 | 1,44 | -0,34 |
| EVI2B | 2,20 | -0,34 |
| LCP1 | 1,43 | -0,33 |
| VMP1 | 1,48 | -0,33 |
| APBB1IP | 1,44 | -0,32 |
| PPIA | 1,59 | -0,30 |
| RPRC1;MAP7D1 | 1,39 | -0,30 |
| RNF213 | 2,12 | -0,30 |
| VASP | 1,44 | -0,29 |
| HNRNPA3 | 1,32 | -0,29 |
| PRDX5 | 1,32 | -0,29 |
| TES | 1,38 | -0,25 |
| SH3BP1 | 1,39 | -0,24 |
| APRT | 1,64 | -0,23 |
| DNAJB11 | 1,72 | -0,23 |
| RBMX | 1,51 | -0,23 |
| COX7A2 | 1,31 | -0,23 |
| PTPN12 | 1,92 | -0,22 |
| SAMHD1 | 1,38 | -0,21 |
| PNPLA6 | 1,38 | -0,20 |
| ERH | 1,45 | -0,19 |
| HCK | 2,33 | -0,19 |
| S100A9 | 1,32 | -0,18 |
| PICALM | 1,89 | -0,18 |
| TUBA1B | 2,12 | -0,17 |
| HNRNPD | 1,49 | -0,11 |
| DNAJC3 | 1,84 | -0,11 |
| UBAP2L | 1,30 | -0,11 |
| Sep-02 | 1,33 | -0,10 |
| PGAM1 | 2,67 | -0,09 |
| TLN1 | 1,69 | -0,09 |

**Table S2**: Proteins which are significantly increased in BL001-treated MDM10 obtained from individuals with type 1 diabetes mellitus.

| Protein | -Log(p) | Log2(FC) |
|---|---|---|
| CCT4 | 1,50 | 0,08 |
| DYNC1H1 | 1,46 | 0,10 |
| RPL23A | 1,83 | 0,10 |
| UBE2N;UBE2NL | 1,40 | 0,12 |
| CCT5 | 1,40 | 0,12 |
| RTCB | 1,31 | 0,12 |
| PLEC | 1,45 | 0,13 |
| FERMT3 | 1,74 | 0,14 |
| CCDC88A | 1,95 | 0,15 |
| SND1 | 2,48 | 0,16 |
| PDCD5 | 1,31 | 0,18 |
| BCAP31 | 1,58 | 0,18 |
| EPRS | 1,91 | 0,18 |
| NSF | 1,86 | 0,18 |
| STIP1 | 2,05 | 0,18 |
| CAPG | 2,15 | 0,18 |
| RARS | 2,12 | 0,19 |
| YARS | 1,68 | 0,20 |
| NME1-NME2;NME2 | 2,75 | 0,20 |
| RUFY1 | 1,74 | 0,20 |
| ALYREF | 1,76 | 0,20 |
| EIF4A1 | 1,46 | 0,21 |
| EIF4G2 | 1,66 | 0,21 |
| AHCY | 3,15 | 0,21 |
| SNX8 | 1,45 | 0,22 |
| PRRC1 | 2,99 | 0,24 |
| PSMA6 | 1,44 | 0,24 |
| KYNU | 1,31 | 0,26 |
| AP2M1 | 2,20 | 0,26 |
| AGPS | 1,33 | 0,26 |
| RPS14 | 2,29 | 0,27 |
| LRPPRC | 2,86 | 0,27 |
| ATP6V1G1 | 2,21 | 0,27 |
| TBXAS1 | 1,67 | 0,28 |
| PHB | 1,37 | 0,28 |
| PDS5B | 1,32 | 0,28 |
| RPN1 | 1,80 | 0,28 |
| FGR | 1,73 | 0,29 |
| ATP6V1D | 1,69 | 0,30 |
| PRKCB | 1,33 | 0,30 |
| UFL1 | 1,52 | 0,30 |
| COPA | 1,44 | 0,30 |
| SULT1A4;SULT1A3 | 1,42 | 0,30 |
| TALDO1 | 1,63 | 0,30 |
| MOB1A | 1,50 | 0,32 |
| SACM1L | 2,20 | 0,33 |

| Protein | -Log(p) | Log2(FC) |
|---|---|---|
| SGPL1 | 1,37 | 0,34 |
| CES1 | 1,46 | 0,34 |
| CHMP4B | 1,66 | 0,34 |
| TARDBP;TDP43 | 1,56 | 0,37 |
| GRPEL1 | 1,51 | 0,37 |
| EMC2 | 2,79 | 0,37 |
| ALDH3A2 | 2,66 | 0,37 |
| TNS3 | 1,69 | 0,37 |
| AARS | 1,57 | 0,38 |
| RAB18 | 1,63 | 0,39 |
| EIF2S3;EIF2S3L | 2,42 | 0,40 |
| GLS | 1,91 | 0,40 |
| ARPC5 | 1,59 | 0,41 |
| OTUB1 | 1,51 | 0,41 |
| DARS | 1,62 | 0,43 |
| HIST1H1E | 1,89 | 0,43 |
| ADD1 | 1,39 | 0,44 |
| DHRS4 | 1,42 | 0,44 |
| CLPB | 2,72 | 0,46 |
| LRRC47 | 1,48 | 0,47 |
| PSAP | 2,58 | 0,50 |
| IRF5 | 1,95 | 0,50 |
| CPPED1 | 1,67 | 0,53 |
| VAT1 | 3,07 | 0,55 |
| FLNB | 1,41 | 0,55 |
| ATXN3 | 1,31 | 0,56 |
| COL4A3BP | 1,47 | 0,57 |
| PIK3R1 | 1,57 | 0,59 |
| IPO9 | 1,64 | 0,62 |
| CTSD | 1,79 | 0,66 |
| PML | 1,66 | 0,68 |
| FCHO2 | 2,33 | 0,70 |
| TBC1D2 | 1,41 | 0,70 |
| SHMT2 | 1,36 | 0,75 |
| PLOD3 | 1,35 | 0,85 |
| PC | 1,33 | 0,86 |
| OAT | 2,68 | 0,92 |
| CALCOCO1 | 1,89 | 0,92 |
| SQSTM1 | 2,29 | 1,12 |
| CHCHD2;CHCHD2P9 | 2,55 | 1,13 |
| CD40 | 1,54 | 1,15 |
| GM2A | 2,47 | 1,42 |
| TFRC | 1,68 | 1,79 |

**Table S3**: Proteins which are significantly ($p<0.05$) decreased in BL001-treated MDM1 obtained from individuals with type 1 diabetes mellitus.

| Protein | -Log p | Log2(FC) |
|---|---|---|
| CD14 | 2,15 | -2,28 |
| FAM175B | 1,33 | -1,18 |
| CD163 | 1,42 | -1,12 |
| CYCS | 1,62 | -1,10 |
| ACO1 | 2,20 | -0,96 |
| LYPLA2 | 1,54 | -0,93 |
| S100A10 | 1,68 | -0,92 |
| UNC13D | 1,58 | -0,88 |
| RPN2 | 1,68 | -0,85 |
| SERPINH1 | 2,59 | -0,84 |
| ADPGK | 1,50 | -0,75 |
| PSMA2 | 1,65 | -0,68 |
| PSMB10 | 1,41 | -0,65 |
| MT-CO3 | 2,24 | -0,60 |
| SIRPA;SIRPB1 | 2,24 | -0,58 |
| GNB2 | 1,63 | -0,58 |
| EHD1 | 2,22 | -0,57 |
| U2AF2 | 1,62 | -0,57 |
| PTGR1 | 1,38 | -0,56 |
| TAPBP | 2,96 | -0,56 |
| PAFAH1B2 | 1,45 | -0,56 |
| PGLS | 1,77 | -0,53 |
| RPS2 | 1,34 | -0,51 |
| TIMM50 | 2,25 | -0,51 |
| HIP1 | 1,82 | -0,50 |
| ACTL6A | 1,36 | -0,49 |
| NDUFA9 | 2,38 | -0,49 |
| SEC11A | 1,37 | -0,48 |
| ERAP1 | 2,13 | -0,46 |
| ACSL4 | 1,76 | -0,45 |
| QARS | 2,25 | -0,44 |
| CRYZ | 1,44 | -0,42 |
| THEMIS2 | 1,67 | -0,40 |
| HNMT | 1,82 | -0,40 |
| CTSZ | 1,54 | -0,39 |
| UBE2N;UBE2NL | 2,12 | -0,38 |
| GBF1 | 1,61 | -0,37 |
| VPS13C | 1,44 | -0,36 |
| DCTN1;DKFZp686E0752 | 1,67 | -0,34 |
| VCL | 1,55 | -0,34 |
| ATP2A3 | 1,46 | -0,33 |
| COTL1 | 1,35 | -0,33 |
| TMEM43 | 1,41 | -0,33 |
| CASP1 | 1,49 | -0,33 |
| MTAP | 1,51 | -0,32 |
| TMEM173 | 1,31 | -0,32 |
| TMCO1 | 1,62 | -0,32 |
| PPA1 | 1,92 | -0,29 |
| PSMA4 | 1,71 | -0,28 |
| RAP1B | 1,61 | -0,26 |
| COPG2 | 1,31 | -0,25 |
| MAPRE1 | 1,55 | -0,24 |
| CCDC124 | 1,44 | -0,23 |
| HPRT1 | 1,57 | -0,19 |
| RNF213 | 1,58 | -0,18 |
| PFN1 | 1,35 | -0,18 |
| PSMD7 | 2,09 | -0,18 |
| ACLY | 1,32 | -0,18 |
| EEF2 | 1,53 | -0,18 |
| AP2A1 | 1,71 | -0,17 |
| ATP5F1 | 1,39 | -0,17 |
| FMNL1 | 1,53 | -0,16 |
| PRDX6 | 1,97 | -0,14 |
| ANXA6 | 2,48 | -0,12 |
| SARS | 1,37 | -0,10 |
| ADSS | 1,75 | -0,05 |

**Table S4**: Proteins which are significantly increased in BL001-treated MDM1 obtained from individuals with type 1 diabetes mellitus.

| Protein | -Log p | Log2(FC) |
|---|---|---|
| RAB14 | 1,36 | 0,08 |
| NUCB1 | 1,55 | 0,08 |
| ANXA11 | 1,60 | 0,09 |
| PHB | 2,06 | 0,10 |
| MVP | 1,60 | 0,13 |
| XRCC6 | 1,51 | 0,14 |
| LRRC47 | 1,70 | 0,17 |
| ACADVL | 1,80 | 0,18 |
| CCDC88A | 1,34 | 0,20 |
| PNN | 1,30 | 0,21 |
| EIF6 | 1,43 | 0,21 |
| RPS18 | 2,29 | 0,24 |
| ECHS1 | 2,16 | 0,24 |
| P4HB | 2,86 | 0,25 |
| CLIP1 | 1,35 | 0,25 |
| NCLN | 1,59 | 0,26 |
| WDR61 | 2,47 | 0,28 |
| RPL7A | 1,34 | 0,28 |
| GRPEL1 | 1,37 | 0,29 |
| TCEA1 | 1,79 | 0,29 |
| HNRNPA3 | 1,33 | 0,30 |
| DDX17 | 1,37 | 0,30 |
| CLASP2 | 1,94 | 0,30 |
| PPIF | 1,45 | 0,30 |
| NUDC | 1,82 | 0,31 |
| EDC4 | 2,40 | 0,33 |
| KHSRP | 2,18 | 0,33 |
| ST13 | 2,17 | 0,33 |
| HCFC1 | 1,61 | 0,33 |
| SMARCE1 | 1,45 | 0,34 |
| FBL | 1,88 | 0,37 |
| XRN2 | 1,38 | 0,38 |
| PPP1CC | 1,81 | 0,39 |
| NUP93 | 1,92 | 0,40 |
| VAMP8 | 1,72 | 0,40 |
| GIMAP1 | 2,09 | 0,41 |
| SLC4A1AP | 3,37 | 0,42 |
| NRP1 | 1,54 | 0,43 |
| POLR2A | 1,91 | 0,46 |
| SART1 | 1,43 | 0,53 |
| NOMO1 | 1,90 | 0,57 |
| NUP62 | 1,63 | 0,58 |
| TDP43 | 1,79 | 0,60 |
| TBL1XR1 | 1,34 | 0,60 |
| GOLGA2 | 1,73 | 0,66 |
| FUS | 2,81 | 0,71 |
| SOAT1 | 1,36 | 0,72 |
| RPL36AL | 1,80 | 0,76 |
| RPL8 | 1,33 | 0,80 |
| GOLGB1 | 1,64 | 0,90 |
| HIST1H1E | 1,50 | 0,93 |
| EIF4A1 | 1,73 | 2,32 |
| SPTBN1 | 1,72 | 2,71 |

**Table S5**: Proteins which are significantly (*p<0.05*) decreased in BL001-treated mDCs obtained from individuals with type 1 diabetes mellitus.

| Protein | -Log p | Log2(FC) |
|---|---|---|
| KRT74 | 1,62 | -3,10 |
| PDCD2 | 1,38 | -1,39 |
| ACTG2 | 1,34 | -1,03 |
| PIK3CB | 1,76 | -1,01 |
| KRT84 | 2,20 | -0,92 |
| ITFG1 | 1,33 | -0,85 |
| COL9A3 | 1,32 | -0,78 |
| IFIT3 | 1,73 | -0,73 |
| CD3E | 1,91 | -0,60 |
| PRNP | 1,93 | -0,60 |
| TMEM209 | 1,38 | -0,57 |
| NOL3 | 1,58 | -0,56 |
| TPST2 | 1,39 | -0,55 |
| RFC1 | 1,64 | -0,54 |
| HLA-A;HLA-H | 2,14 | -0,49 |
| CDC42SE2 | 1,36 | -0,45 |
| MYH1;MYH13;MYH4;MYH8 | 1,57 | -0,42 |
| UQCRH | 1,63 | -0,35 |
| CBX1;CBX3 | 1,42 | -0,35 |
| GATAD2B | 1,41 | -0,35 |
| STT3A | 1,46 | -0,34 |
| ASDURF | 1,59 | -0,30 |
| GGCT | 1,39 | -0,29 |
| DAZAP1 | 3,19 | -0,27 |
| EIF5A;EIF5A2 | 1,47 | -0,26 |
| STX11 | 1,93 | -0,23 |
| RTN3 | 1,66 | -0,22 |
| PEA15 | 1,31 | -0,22 |
| COMT | 1,30 | -0,21 |
| TUBB;TUBB2A;TUBB4B | 1,36 | -0,21 |
| UBL4A | 1,87 | -0,21 |
| BAX | 1,32 | -0,20 |
| ERBIN | 1,51 | -0,20 |
| TUBB4A;TUBB4B | 2,05 | -0,19 |
| PPP4C | 1,68 | -0,18 |
| RPL9 | 1,38 | -0,18 |
| SEC23A | 1,51 | -0,18 |
| APEX1 | 1,52 | -0,16 |
| FAM98B | 1,38 | -0,15 |
| THEMIS2 | 1,81 | -0,12 |
| MATR3 | 1,50 | -0,10 |
| PRDX6 | 1,60 | -0,10 |
| ABCD1 | 1,76 | -0,08 |
| MACROH2A1 | 1,48 | -0,07 |

**Table S6**: Proteins which are significantly (*p<0.05*) increased in BL001-treated mDCs obtained from individuals with type 1 diabetes mellitus.

| Protein | -Log p | Log2(FC) |
|---|---|---|
| GNPDA1 | 1,37 | 0,05 |
| APPL1 | 1,65 | 0,07 |
| CORO1B | 1,42 | 0,07 |
| DNAJC1 | 1,76 | 0,08 |
| ARFGEF1;ARFGEF2 | 1,31 | 0,08 |
| PPIB | 1,41 | 0,08 |
| SRP54 | 1,30 | 0,08 |
| VPS4B | 1,61 | 0,08 |
| GLUD1 | 1,65 | 0,09 |
| RAN | 1,93 | 0,09 |
| SARS2 | 1,97 | 0,09 |
| BUB3 | 2,12 | 0,09 |
| PSMD11 | 1,37 | 0,10 |
| FARSB | 1,43 | 0,10 |
| HS1BP3 | 1,81 | 0,10 |
| CRYZ | 4,01 | 0,10 |
| ELAVL1 | 1,50 | 0,10 |
| HSP90B1 | 1,93 | 0,10 |
| MRPL12 | 1,46 | 0,10 |
| STK24;STK26 | 1,64 | 0,11 |
| ATP6V0D1 | 1,58 | 0,11 |
| VPS16 | 1,40 | 0,11 |
| ACADVL | 1,68 | 0,11 |
| RTRAF | 1,30 | 0,11 |
| SAE1 | 1,31 | 0,11 |
| CTBP1 | 1,71 | 0,11 |
| FMR1;FXR1;FXR2 | 2,69 | 0,11 |
| DLAT | 1,69 | 0,12 |
| RPS3 | 1,48 | 0,12 |
| GANAB | 2,75 | 0,12 |
| SNX1 | 1,43 | 0,12 |
| RPL10A | 1,31 | 0,12 |
| CCT6A;CCT6B | 1,82 | 0,12 |
| RAB1A;RAB1B | 1,99 | 0,12 |
| MRE11 | 1,41 | 0,13 |
| SNU13 | 1,81 | 0,13 |
| TRAPPC3 | 1,52 | 0,14 |
| ELMO1;ELMO2 | 1,50 | 0,14 |
| ADSL | 1,63 | 0,14 |
| ATP5F1A | 1,60 | 0,14 |
| ATP5PO | 1,78 | 0,14 |
| MRPL13 | 1,52 | 0,14 |
| MBNL1 | 1,50 | 0,14 |
| NIT1 | 1,42 | 0,14 |
| LILRA1;LILRB1 | 1,32 | 0,15 |
| COMMD7 | 1,34 | 0,15 |
| DCPS | 2,19 | 0,15 |
| TAOK1;TAOK3 | 2,10 | 0,15 |
| POFUT1 | 1,72 | 0,15 |
| HEATR3 | 1,34 | 0,15 |
| P4HA1 | 2,08 | 0,15 |
| TRAPPC11 | 1,58 | 0,16 |
| ATP6V1G1 | 2,27 | 0,16 |
| ERP29 | 1,40 | 0,16 |
| ACOT1;ACOT2 | 1,34 | 0,16 |
| PIK3CG | 1,36 | 0,16 |
| WDR81 | 1,45 | 0,16 |
| ACADSB | 1,67 | 0,17 |
| EXOC8 | 1,63 | 0,18 |
| RPS4X;RPS4Y1;RPS4Y2 | 1,49 | 0,18 |
| POLR2B | 1,39 | 0,18 |
| ELMO1 | 2,17 | 0,18 |
| MESD | 1,45 | 0,18 |
| RPL15 | 1,36 | 0,18 |
| ARL3 | 2,34 | 0,19 |
| EIF1;EIF1B | 1,43 | 0,19 |
| GSDMD | 1,30 | 0,19 |
| ALDH18A1 | 1,38 | 0,19 |
| TM9SF3 | 1,65 | 0,20 |
| MCCC2 | 1,69 | 0,21 |
| RABGAP1 | 2,52 | 0,21 |
| OPTN | 1,51 | 0,22 |
| AIMP2 | 2,61 | 0,23 |
| AGAP3 | 1,37 | 0,24 |
| SRP9 | 2,22 | 0,24 |
| ARSB | 1,52 | 0,25 |
| NCOR2 | 1,40 | 0,25 |
| PWP1 | 3,12 | 0,25 |
| NOP10 | 1,70 | 0,25 |

| Protein | -Log p | Log2(FC) |
|---|---|---|
| CIAO1 | 1,90 | 0,26 |
| GMFB;GMFG | 1,97 | 0,26 |
| OXSR1 | 1,34 | 0,26 |
| RMC1 | 1,34 | 0,27 |
| TAF4 | 2,00 | 0,27 |
| MMAB | 1,58 | 0,27 |
| MRPL11 | 1,91 | 0,27 |
| CTNNA1;CTNNA3 | 1,39 | 0,27 |
| RPL11 | 1,93 | 0,30 |
| BUD31 | 2,64 | 0,31 |
| RRM2B | 1,65 | 0,31 |
| AKT2 | 2,08 | 0,31 |
| AVEN | 1,33 | 0,32 |
| MRPL23 | 1,48 | 0,33 |
| CBR4 | 1,67 | 0,34 |
| UBASH3A | 2,10 | 0,34 |
| SEPTIN3 | 1,70 | 0,36 |
| EARS2 | 2,70 | 0,37 |
| COX17 | 1,35 | 0,39 |
| IGHG2;IGHG4 | 1,43 | 0,39 |
| MRPL1 | 1,31 | 0,40 |
| EED | 2,27 | 0,40 |
| TMEM245 | 1,76 | 0,40 |
| MAOB | 1,33 | 0,40 |
| TANGO6 | 1,46 | 0,41 |
| PTTG1IP | 1,43 | 0,41 |
| FYN;SRC;YES1 | 1,67 | 0,41 |
| ALKBH7 | 1,75 | 0,42 |
| SP1 | 1,77 | 0,47 |
| SNX1;SNX2 | 1,80 | 0,47 |
| RILP | 1,44 | 0,48 |
| UQCC2 | 1,35 | 0,50 |
| CSNK1E | 1,87 | 0,54 |
| LCK | 1,50 | 0,54 |
| ATF6 | 1,54 | 0,55 |
| CASTOR1;CASTOR2 | 1,39 | 0,56 |
| ERVK-6;HERVK_113 | 2,18 | 0,56 |
| EEPD1 | 1,63 | 0,58 |
| UXS1 | 1,36 | 0,58 |
| ATAD2B | 1,39 | 0,58 |
| PUM3 | 1,42 | 0,59 |
| BZW1;BZW2 | 1,40 | 0,62 |
| AIDA | 1,46 | 0,64 |
| KNG1 | 1,32 | 0,64 |
| OXSM | 1,36 | 0,64 |
| NT5C3A | 1,33 | 0,67 |
| ISY1 | 1,96 | 0,68 |
| NELFCD | 1,60 | 0,73 |
| VAMP2 | 1,38 | 0,74 |
| WDR74 | 1,61 | 0,75 |
| GNPDA2 | 1,64 | 0,55 |
| APPL2 | 1,64 | 0,56 |
| CORO1B | 1,64 | 0,56 |
| DNAJC2 | 1,64 | 0,57 |
| ARFGEF1;ARFGEF3 | 1,64 | 0,57 |
| PPIB | 1,63 | 0,57 |
| SRP55 | 1,63 | 0,58 |
| VPS4B | 1,63 | 0,58 |
| GLUD2 | 1,63 | 0,59 |
| RAN | 1,63 | 0,59 |
| SARS3 | 1,63 | 0,60 |
| BUB4 | 1,63 | 0,60 |
| PSMD12 | 1,63 | 0,61 |
| FARSB | 1,63 | 0,61 |
| HS1BP4 | 1,63 | 0,61 |
| CRYZ | 1,63 | 0,62 |
| ELAVL2 | 1,63 | 0,62 |
| HSP90B2 | 1,63 | 0,63 |
| MRPL13 | 1,63 | 0,63 |
| STK24;STK27 | 1,62 | 0,64 |
| ATP6V0D2 | 1,62 | 0,64 |
| VPS17 | 1,62 | 0,65 |
| ACADVL | 1,62 | 0,65 |
| RTRAF | 1,62 | 0,66 |
| SAE2 | 1,62 | 0,66 |
| CTBP2 | 1,62 | 0,66 |
| FMR1;FXR1;FXR3 | 1,62 | 0,67 |
| DLAT | 1,62 | 0,67 |

**Table S7**: GO enrichment terms associated to mitochondria for up-regulated proteins in BL001-treated mDCs as compared to untreated mDCs.

| GO Biological processes | Proteins |
|---|---|
| mitochondrial translation | SARS2/MRPL12/MRPL13/MRPL11/MRPL23/EARS2/MRPL1/UQCC2/MRPL3 |
| mitochondrial gene expression | SARS2/MRPL12/MRPL13/MRPL11/MRPL23/EARS2/MRPL1/UQCC2/MRPL3 |
| mitochondrial translational elongation | MRPL12/MRPL13/MRPL11/MRPL23/MRPL1/MRPL3 |
| mitochondrial translational termination | MRPL12/MRPL13/MRPL11/MRPL23/MRPL1/MRPL3 |
| mitochondrial RNA metabolic process | SARS2/MRPL12/EARS2 |
| mitochondrial ATP synthesis coupled proton transport | ATP5F1A/ATP5PO |
| inner mitochondrial membrane organization | ATP5F1A/ATP5PO |
| mitochondrial respiratory chain complex III assembly | UQCC2 |
| positive regulation of mitophagy in response to mitochondrial depolarization | OPTN |
| positive regulation of mitochondrial membrane potential | AKT2 |
| mitochondrial membrane organization | ATP5F1A/ATP5PO/ALKBH7 |
| mitochondrial DNA replication | RRM2B |
| positive regulation of autophagy of mitochondrion in response to mitochondrial depolarization | OPTN |
| regulation of autophagy of mitochondrion in response to mitochondrial depolarization | OPTN |
| mitochondrial transcription | MRPL12 |
| positive regulation of mitochondrial translation | UQCC2 |
| mitochondrial transport | SAE1/ATP5F1A/ATP5PO/ALKBH7 |
| response to mitochondrial depolarisation | OPTN |
| positive regulation of autophagy of mitochondrion | OPTN |
| mitochondrial cytochrome c oxidase assembly | COX17 |
| mitochondrial transmembrane transport | ATP5F1A/ATP5PO |
| mitochondrial respiratory chain complex assembly | COX17/UQCC2 |
| regulation of mitochondrial translation | UQCC2 |
| regulation of mitochondrial gene expression | UQCC2 |
| positive regulation of protein targeting to mitochondrion | SAE1 |
| positive regulation of mitochondrion organization | SAE1/OPTN |
| regulation of autophagy of mitochondrion | OPTN |
| regulation of protein targeting to mitochondrion | SAE1 |
| positive regulation of establishment of protein localization to mitochondrion | SAE1 |
| regulation of mitochondrion organization | SAE1/OPTN |
| regulation of establishment of protein localization to mitochondrion | SAE1 |
| regulation of mitochondrial membrane potential | AKT2 |
| regulation of mitochondrial membrane permeability | ALKBH7 |
| autophagy of mitochondrion | OPTN |
| mitochondrion disassembly | OPTN |
| protein targeting to mitochondrion | SAE1 |
| establishment of protein localization to mitochondrion | SAE1 |
| protein localization to mitochondrion | SAE1 |
